# From Phenomics to Genomics: Macro-GWAS of Almond Morphology and Quality

**DOI:** 10.64898/2026.07.06.736816

**Authors:** Jorge Mas-Gómez, Manuel Rubio, Henri Duval, Federico Dicenta, Pedro José Martínez-García

## Abstract

In plant breeding and genetics, recent advances in high-throughput phenotyping are beginning to meet the growing demand for large-scale, high-quality phenotypic data that emerged after the development of next-generation sequencing technologies. Recent developments in phenomics have been incorporated into almond breeding programs, facilitating the large-scale acquisition of quantitative phenotypes and the dissection of the genetic architecture underlying morphological and quality-related traits. The implementation of a high-throughput phenotyping platform integrating RGB and hyperspectral imaging with genotyping using the 60K almond SNP array enabled the large-scale characterization of almond populations and the identification of 567 robust marker–trait associations across 66 traits. These analyses revealed two major genomic hotspots on chromosomes 2 and 5 associated with morphological and quality-related traits. These regions harbored biologically relevant candidate genes, including genes associated with OVATE family proteins, brassinosteroid signaling, protein ubiquitination, and acyl-CoA metabolism, as well as other regulators of organ growth, cell proliferation, hormone signaling, and seed development. Furthermore, a novel candidate gene encoding a COMT-like O-methyltransferase involved in lignin biosynthesis was identified and proposed to contribute to shell hardness, a major genetically controlled trait in almond. Together, these findings demonstrate the potential of integrating high-throughput phenomics and genomics to dissect complex traits, identify candidate genes, and accelerate genomics-informed breeding in almond.

## Introduction

Almond [*Prunus dulcis* (Mill.) D.A. Webb] is a diploid stone fruit species of the *Rosaceae* family, with a genome organized into eight chromosomes, as occurs in other *Prunus* crops such as peach (*P. persica*) and sweet cherry (*P. avium*) [1]. The development of new cultivars in *Prunus* species is costly and time-consuming. Their extended juvenile phase means that breeding programs typically require approximately 12–15 years to release an improved cultivar [2]. Consequently, enhancing the efficiency of the breeding process is a major priority. In this context, elucidating the genetic architecture underlying the traits of interest can provide valuable biological insights and facilitate their effective use in the development of more efficient breeding programs [3]. Among the traits of greatest relevance in almond breeding, kernel morphology and compositional quality have a direct influence on both market value and industrial suitability [4]. Kernel size and shape affect consumer preferences, processing requirements, final use, and resistance to breakage, while compositional traits such as fat, protein, and sucrose determine processing performance, sensory quality, and suitability for food and oil production [5].

Despite the importance of these traits in almond, their underlying genetic basis remains poorly understood, as previous studies have generally relied on reduced populations and low-density marker datasets [6, 7]. More recently, two studies have used high-density maps to investigate almond morphological and quality traits [8, 9]. In general, these studies suggest that most of these traits have a complex quantitative architecture and are controlled by multiple genes [6–9]. Shell hardness is a notable exception and has been studied more extensively because it is strongly correlated with two easily measured traits, shell weight and crack-out percentage [10]. These traits have been associated with a major-effect region on chromosome 2 [6, 10–12]. However, for the rest of the traits, phenotyping remains the main bottleneck, limiting the evaluation of the large populations required to identify the genes underlying these traits [13].

The limitations associated with conventional phenotyping represent a widespread challenge in plant breeding programs [13]. To overcome this constraint, breeders are increasingly adopting high-throughput phenotyping platforms (HTPPs) that integrate computer vision, advanced sensor technologies, and artificial intelligence [14]. These technologies facilitate the rapid and efficient evaluation of large breeding populations. Their potential has also been demonstrated in almond breeding showing that image-based phenotyping could identify the same quantitative trait loci (QTLs) associated with fruit size and shape as conventional manual measurements [8]. In line with these developments, we established a phenomics platform that combines RGB and hyperspectral imaging and validated its application in large breeding populations and studied the heritability of the traits [15, 16].

The expansion of phenotyping capacity and trait coverage is expected not only to increase breeding efficiency but also to generate new biological knowledge about the genetic factors underlying complex traits [17]. In the almond tree, however, relatively few studies have proposed candidate genes associated with morphological and quality-related traits [9]. One of the most compelling candidates identified to date is a NAC-domain regulator orthologous to NST1, which has been associated with shell hardness through its role in secondary cell-wall formation and lignification [11, 18]. By contrast, the genetic regulation of seed size, shape, and chemical composition has been more extensively investigated in model and other crop species. Members of the OVATE family are particularly relevant to shape-related traits because they regulate organ morphology by modifying patterns of cell division and growth [19, 20]. Seed and organ size may also be influenced by transcription factors involved in developmental patterning and hormonal responses, including AP2 and NAC proteins and brassinosteroid-responsive regulators such as BZR1, BIM, and related bHLH transcription factors [21–23]. Protein ubiquitination also plays a major role in organ growth, with the DA1–DA2–BIG BROTHER pathway restricting seed and organ size through the regulation of cell proliferation [24–26]. In addition, genes involved in cell-wall biosynthesis and modification, including pectin-related enzymes, glycosyltransferases, wall-associated kinases, arabinogalactan proteins, and WAT1-related transporters, may contribute to variation in tissue expansion, morphology, and fiber-related traits [27–31]. In addition, studies in major oilseed crops have shown that variation in oil content is related to carbon allocation, fatty-acid biosynthesis and lipid assembly [32, 33]. In this context, central components of fatty-acid biosynthesis, such as acetyl-CoA, have been shown to play a fundamental role in determining carbon flux towards lipid production [33]. Similarly, seed protein content and composition are influenced by amino-acid biosynthesis and by the synthesis, processing, and mobilization of storage proteins [34, 35].

Here, we used a high-throughput phenotyping platform integrating RGB and hyperspectral imaging to evaluate large almond populations. These populations were also genotyped using the Axiom™ Almond 60K SNP array. The resulting large-scale phenotypic and genotypic dataset was used to investigate the genetic architecture of kernel and shell morphological traits, as well as kernel quality traits, through a macro GWAS framework based on multilocus methods. SNP effects were examined, and genes located near significant markers were identified and functionally annotated to reveal potential candidate genes underlying these traits.

## Results

### RGB Imaging Analyses

The fine-tuned AI-based object detection models performed strongly for both kernel and shell detection (Appendix 1: Figures 1 and 2). By the end of training, precision and recall were close to or higher than 0.93, suggesting limited false detections and missed objects. Both models reached mAP@50 values near 0.99, defined as the average precision computed at an intersection over union (IoU) threshold of 0.50, and mAP@50–95 values above 0.97, defined as the mean average precision averaged across IoU thresholds from 0.50 to 0.95, confirming accurate object localization across both moderate and stringent IoU thresholds.

Elliptic Fourier Analysis (EFA) was applied independently to the binary masks of kernels and shells. After a preliminary visual inspection, 10 harmonics were retained for both datasets. PCA was then carried out using the EFA coefficients (EF-PCs). In the kernel dataset, the first three principal components explained 67.60%, 12.10%, and 4.19% of the total variation, respectively, whereas in the shell dataset they explained 70.80%, 13.50%, and 4.58%, respectively (Appendix 2: Figures 3 to 8). The first 10 principal components were retained for downstream analyses, and their effects on shape variation are shown graphically in Appendix 2: Figures 5 and 8.

### Hyperspectral Imaging Analyses

The pre-processed mean spectrum was obtained for all samples from the Santomera 2023 and 2024 environments (Appendix 3: Figure 9). DOP was applied to the Reference and Santomera 2023 datasets by employing one EPO component and an EPS value of 0.19. The Reference set spectra were subsequently used to develop PLS models for quality traits (Appendix 3: Figure 10 and 11-14, and Supplementary Table 3). Fat showed the highest predictive performance, with a cross-validation R^2^ of 0.767 and RMSE of 1.753, and an external validation R^2^ of 0.687 and RMSE of 1.651. Protein showed similar performance in cross-validation, with an R^2^ of 0.758 and RMSE of 0.890, and external R^2^ reached 0.607 and RMSE 1.038. Fiber and sucrose showed a lower predictive capacity. The fiber reached a cross-validation R^2^ of 0.392 and external R^2^ of 0.310, with RMSE values of 1.064 and 1.171, respectively. Sucrose showed moderate predictive performance, with a cross-validation R^2^ of 0.277 and RMSE of 0.415, while external validation resulted in an R^2^ of 0.352 and RMSE of 0.426. In addition, the latent variable scatterplots showed that samples were generally well distributed across the PLS space for all quality traits.

### Phenotypic Data Analysis

After processing and curating all pictures, 46,083 kernels and 32,136 shells were analysed, reaching a total of 78,219 elements (Supplementary Table 4 and Appendix 4: Figure 15-30). Overall, Santomera 2022 and 2024 showed the largest kernel and shell sizes, whereas Santomera 2023 consistently presented the lowest values. Kernel length and area were highest in Santomera 2022 and 2024, with mean values of around 22.50 mm and 234 mm^2^, respectively, followed by France 2022. The smallest kernels were observed in Santomera 2023, with a mean area of 196.28 mm^2^ and a mean length of 20.63 mm. Kernel width followed a similar trend, with the highest mean value in Santomera 2024 (14.28 mm), while mean predicted kernel thickness was highest in Santomera 2022 (mean value = 8.14 mm) and lowest in Santomera 2023 (mean value = 6.84 mm). Kernel shape traits showed no major differences among Santomera environments, with circularity mean value ranging from 0.75 to 0.77, although France 2022 showed a lower value (mean value = 0.73), likely due to its different genotype composition. Shell size traits followed the same pattern as kernel traits, with the largest values in Santomera 2022 and 2024 and the lowest in Santomera 2023. Some shell shape and morphometric traits showed moderate environmental variation, particularly shell width_25-75_ ratio and shell EF-PC1, the latter being highest in Santomera 2024 (mean EF-PC1 = 2.00) and the lowest in Santomera 2023 (mean EF-PC1 = -3.44). Kernel-to-shell (KS) size ratios showed slight differences among environments, except for KS width_75_, which was higher in Santomera 2022 (mean value = 0.68) than in 2023 and 2024 (mean value = 0.62). Greater variation was observed for KS shape ratios, especially KS ellipse ratio and ratios related to width_25_. The weight traits showed clearer environmental differences, with the highest kernel and shell weights in Santomera 2022 (mean values 3.64 g and 1.11 g respectively), followed by Santomera 2024 (mean values 3.51 g and 0.97 g respectively), while Santomera 2023 showed the lowest values (mean values 2.73 g and 0.75 g respectively). Finally, quality traits showed differences between Santomera 2023 and 2024, with a higher fat content in 2023 (mean value = 61.61%) than in 2024 (59.28%), with the opposite pattern in protein (19.64% and 20.77%, respectively). Fiber was similar in both years (11.51% and 11.69% respectively), while sucrose was slightly higher in Santomera 2024 (3.98% vs 3.54%).

When BLUEs were described by family, differences were observed between genetic backgrounds, mainly for size- and weight-related traits. For kernel size traits, Marcona-related families generally showed the highest values, particularly Antoñeta *×* Marcona and Florida *×* Marcona, with mean kernel areas of 240.25 and 232.70 mm^2^, respectively, and high width values of 15.01 and 14.20 mm. In contrast, Antoñeta *×* Penta and Antoñeta *×* Tardona showed smaller kernels, with mean areas of 197.9 and 193.8 mm^2^. R1000 *×* Desmayo showed a different pattern, with relatively high kernel length (22.17 mm) but lower width-related values. Germplasm accessions also showed high kernel area and length values, 235.05 mm^2^ and 23.03 mm. For predicted kernel thickness, Marcona-related families again showed the highest values, especially Antoñeta *×* Marcona, Florida *×* Marcona, and Marcona *×* S4017, with mean values of 8.02, 7.77, and 7.85 mm, respectively, whereas Antoñeta *×* Tardona showed the lowest value (6.94 mm). Kernel shape and morphometric traits mainly reflected differences in elongation, with R1000 *×* Desmayo and germplasm showing higher length-to-width ratios, around mean values of 1.72, compared with Marcona-related families, around 1.5. Differences among Marcona-related families were also observed for kernel EF-PC4, with Antoñeta *×* Marcona showing lower values (mean value = -0.95) than Florida *×* Marcona (mean value = 0.73), while Marcona *×* S4017 was intermediate (mean value = 0.07). Regarding shell size traits, a similar pattern was observed. Antoñeta *×* Marcona and Florida *×* Marcona showed the largest shell areas, with mean values of 561.31 and 549.09 mm^2^, respectively, and high shell width values of 24.12 and 23.25 mm. In contrast, Antoñeta *×* Tardona showed the lowest shell area (435.49 mm^2^). R1000 *×* Desmayo and germplasm presented relatively long shells, 32.01 and 32.25 mm, respectively. For shell shape and morphometric traits, the main differences were also associated with elongation. R1000 *×* Desmayo showed higher length-to-width ratios, mean value around 1.54, whereas Marcona-related families showed wider and less elongated shells with mean values around 1.30. Shell EF-PC3 also separated families, with Florida *×* Marcona showing clearly lower values (mean value = -2.53) than Antoñeta *×* Penta and Antoñeta *×* Tardona, which showed positive mean values of 0.84 and 0.70, respectively. KS size ratios showed only slight differences among families, with most mean values around 0.70 for KS length, 0.62 for KS width, and 0.43 for KS area. KS shape ratios also showed limited variation. For weight traits, Antoñeta *×* Marcona showed the highest mean kernel and shell weights, with values of 1.16 g and 4.38 g, respectively, followed by Florida *×* Marcona and Marcona *×* S4017. In contrast, Antoñeta *×* Tardona and Antoñeta *×* Penta showed lower kernel weights, mean values around 0.8–0.85 g, consistent with their smaller kernel and shell sizes.

Broad-sense heritability estimates showed a strong genetic contribution for several traits (Figure 1). The three most heritable traits were KS weight (H^2^ = 0.88), KS symmetry_h_ (H^2^ = 0.71), and shell weight (H^2^ = 0.70), indicating that these traits were largely controlled by genotypic differences. In contrast, the traits most affected by the environment were predicted kernel thickness, kernel weight, and KS length-width-related traits, with environmental components of 0.42, 0.40, and 0.29, respectively. Within kernel morphology traits, kernel length and kernel perimeter showed the highest heritability values, with H^2^ = 0.57 and H^2^ = 0.51, respectively. For kernel shape and morphometric traits, the highest values were observed for kernel EF-PC1 (H^2^ = 0.67) and kernel ellipse ratio (H^2^ = 0.65), followed closely by kernel length-width ratio (H^2^ = 0.64). In shell morphology traits, shell length and shell perimeter were the most heritable traits, with H^2^ = 0.59 and H^2^ = 0.52, respectively. For shell shape and morphometric traits, shell EF-PC1 showed the highest heritability (H^2^ = 0.65), followed by shell length-width ratio and shell ellipse ratio, both with H^2^ = 0.62. For KS ratios, the highest heritability values were observed for KS symmetry_h_ (H^2^ = 0.71) and KS length (H^2^ = 0.66), while several width-related KS ratios also showed high values around H^2^ = 0.65. Among weight-related traits, KS weight and shell weight showed the highest heritability, with H^2^ = 0.88 and H^2^ = 0.70, respectively, whereas kernel weight showed a lower heritability value (H^2^ = 0.41) and a strong environmental component (0.40). Finally, among quality traits, predicted fiber showed the highest heritability (H^2^ = 0.55), followed by predicted sucrose and predicted protein, both with H^2^ = 0.48, while predicted fat showed a slightly lower value (H^2^ = 0.47).

**Figure 1.**
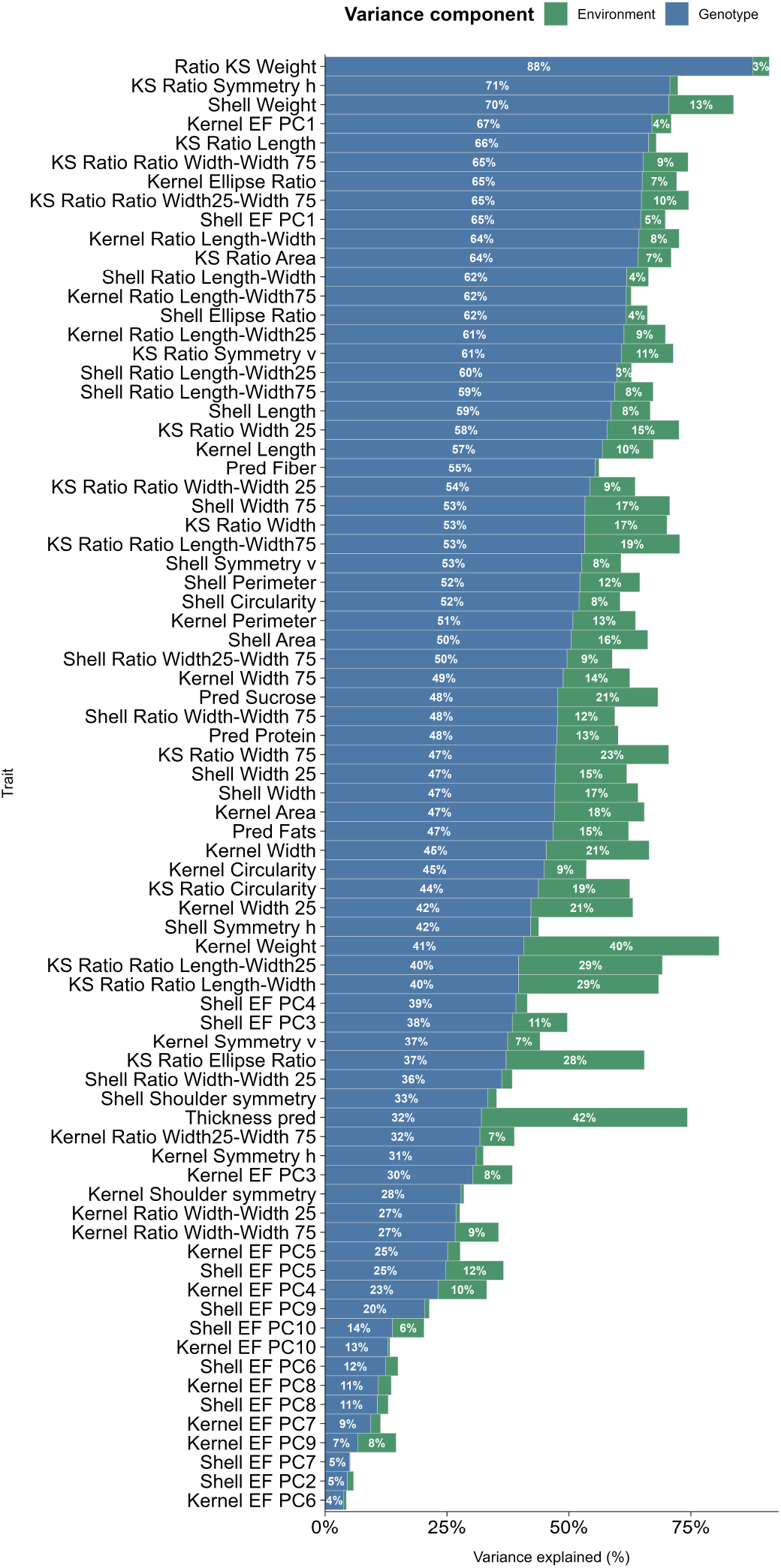
Variance components of the traits studied. Genotype values correspond to H^2^.

The correlation matrix revealed several well-defined trait clusters, mainly corresponding to quality traits, size-related traits, shape descriptors, and kernel-to-shell ratios (Appendix 4: Figure 31). Quality traits formed one of the strongest clusters, with predicted fat showing a highly negative correlation with predicted protein (r = -0.99) and fiber (r = -0.97), whereas protein, fiber, and sucrose were positively associated. A second major cluster grouped kernel and shell size traits, where area, perimeter, width, length, and weight were strongly correlated within and between kernel and shell. Shape-related traits formed another clear cluster, mainly driven by the contrast between rounded and elongated morphologies, as shown by the strong negative correlations between ellipse ratio and length-to-width ratio in both kernel and shell. Additionally, the KS ratios formed a cluster that included ratios between length and width measurements, as well as ratios among the different width measurements.

### Genotyping

A total of 60,581 SNPs were processed with Axiom Analysis Suite v5.1.1.1 and assigned to different quality classes. The largest group corresponded to PolyHighResolution markers, with 42,820 SNPs (70.68%), followed by Other (7,600 SNPs; 12.55%), OTV (3,643 SNPs; 6.01%), NoMinorHom (3,638 SNPs; 6.01%), CallRateBelowThreshold (2,721 SNPs; 4.49%) and MonoHighResolution (159 SNPs; 0.26%) (Table 1). Markers classified as PolyHighResolution or NoMinorHom were retained as the starting high-quality dataset, yielding 46,458 SNPs. This set was further filtered by minor allele frequency (MAF *≥* 5%), leaving 37,251 SNPs. After excluding 3,105 markers showing Mendelian inconsistencies, 34,146 SNPs were retained for downstream analyses. Only for population structure analyses, the final high-quality SNPs set was pruned, achieving a set of 4,808 SNPs.

**Table 1.**
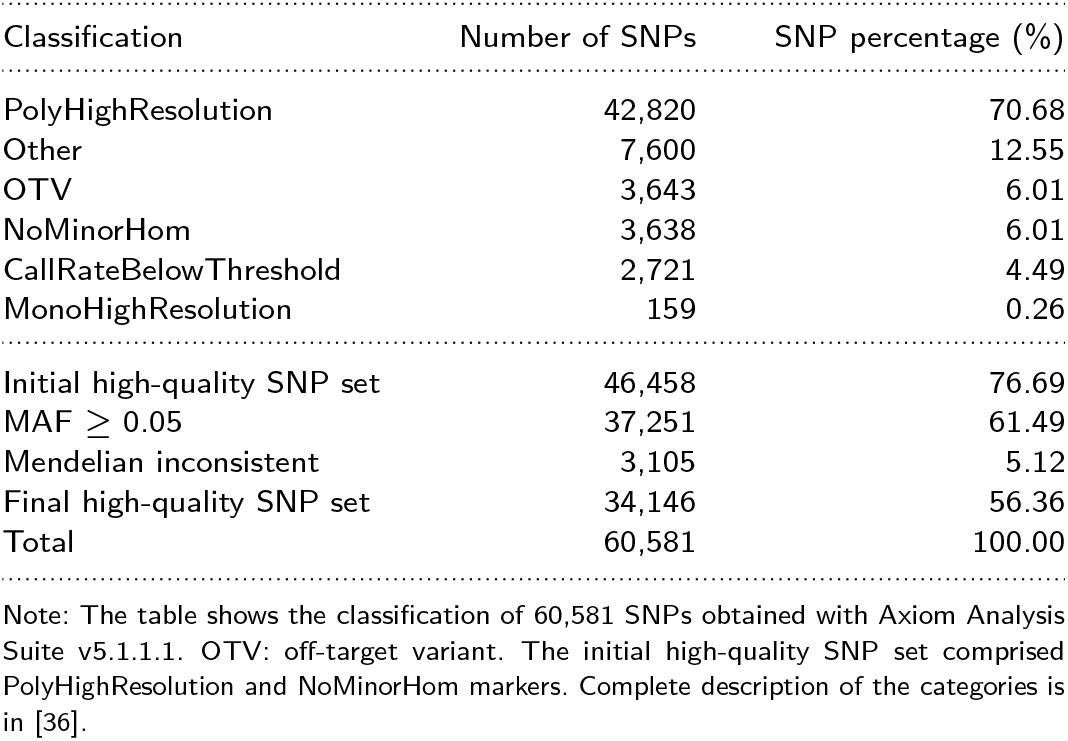
SNP classification.

### GWAS

To account for the potential effects of population structure, ancestry proportions were estimated using *fastSTRUCTURE*, which identified nine genetic clusters (Appendix 5: Figure 32). The first cluster was mainly represented by the Marcona *×* S4017 family, together with germplasm accessions such as Genco, Belona, and Soleta. The second cluster was largely associated with the Marcona genetic background, including the Florida *×* Marcona and Antoñeta *×* Marcona families, as well as related germplasm accessions such as ITAP. The third cluster included the Antoñeta *×* Penta and Antoñeta *×* Tardona families, together with related cultivars and varieties such as Penta, Antoñeta, Tuono, Makako, Lauranne, Marta, and Languedoc. The fourth cluster was mainly composed of Spanish germplasm accessions, including Peraleja, Vivot, Malagueña, and Del Cid. The fifth and sixth clusters were primarily associated with the R1000 *×* Desmayo family, with Q5 mainly reflecting the Desmayo genetic background and Q6 the R1000 background. Cluster Q7 comprised a mixed group of cultivars from France, Spain, and Italy, including Ferragnes, Glorieta, and Cristomorto. Cluster Q8 was mostly represented by cultivars related to the USA genetic background, such as Butte, Nonpareil, and Wood Colony, together with French cultivars such as Fournat and Bartre. Finally, Q9 grouped several *Prunus*-related species, including *Prunus webbii, Prunus vavilovii*, and interspecific *Prunus* hybrids.

Genome-wide association analyses included 66 traits and identified a total of 567 unique marker–trait associations supported by at least two methods (Supplementary Table 5 and Figure 2). The largest number of associations was detected for shell shape traits, with 155 associations, followed by kernel-to-shell ratios with 128 associations, kernel shape with 93 associations, shell morphology with 76 associations, kernel morphology with 73 associations, and both weight-related and predicted quality traits with 21 associations each. The strongest association overall was detected for KS weight with marker AX-586052448 (Pd02: 19,329,225 bp), which showed the highest LOD score (LOD = 19.37). Several genomic regions showed a high concentration of associations (hotspots), especially on Pd02 and Pd05. For kernel morphology, a strong cluster was observed around AX-586093174 (Pd05: 4,251,871 bp) and AX-586094611 (Pd05: 4,159,246 bp), involving kernel area and width-related traits. Kernel shape associations showed a clear concentration on Pd02 and Pd07, especially for aspect ratio-related shape traits. Strong associations were identified around AX-586053067 (Pd02: 20,041,986 bp), AX-586057826 (Pd02: 24,369,150 bp), AX-586057830 (Pd02: 24,370,110 bp), and AX-586135564 (Pd07: 16,396,912 bp).

**Figure 2.**
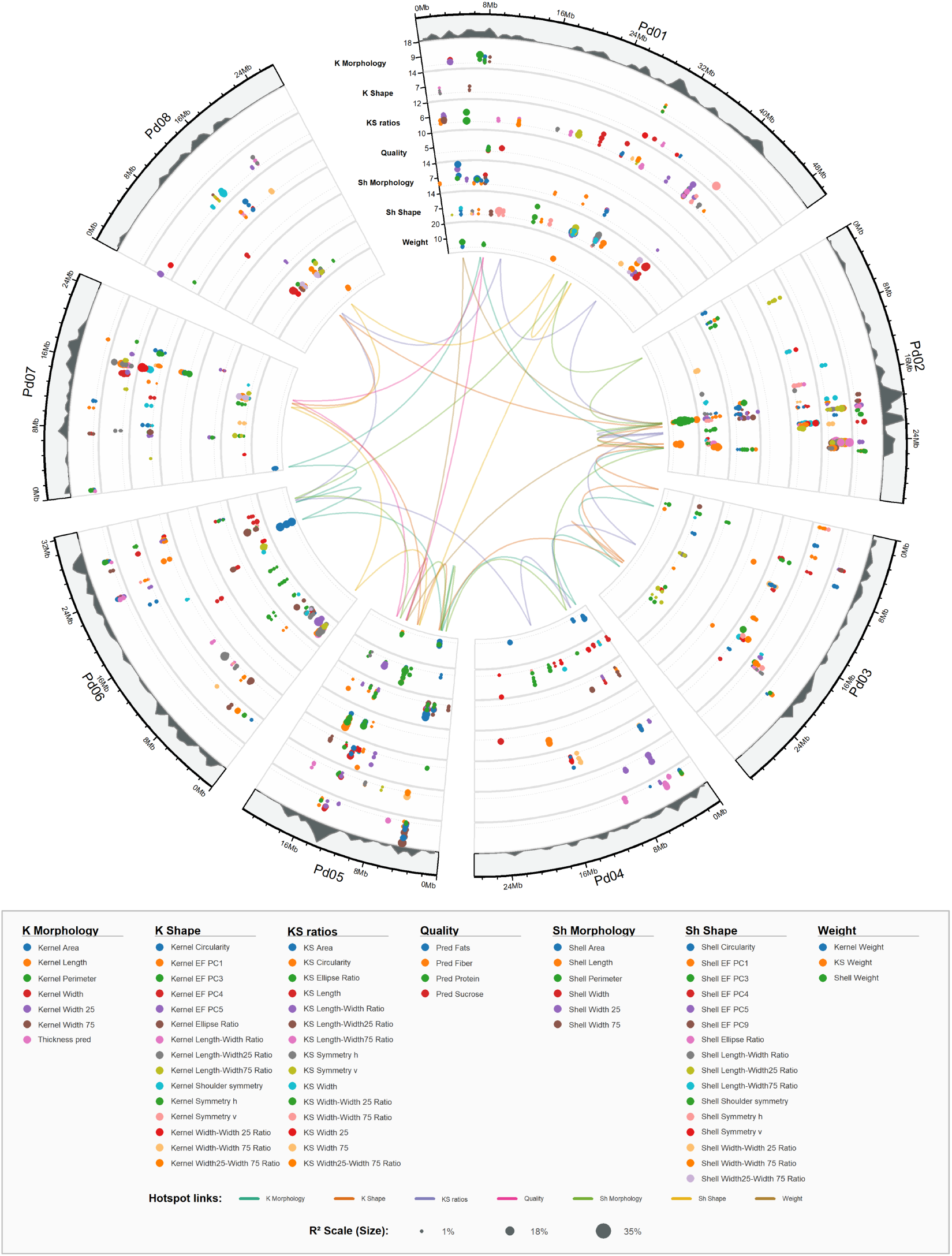
SNP–trait associations identified by at least two methods. Traits are indicated by the colour of each ring, the LOD score is shown, and the point size represents the *r*^2^ value. Linking lines indicate genomic regions containing more than 20% of the traits from the same group within a 200-Mbp window, highlighting group-specific multitrait hotspots.

For shell morphology, hotspots were mainly observed on Pd01, Pd02, and Pd05. On Pd01, relevant associations included shell area with AX-586014654 (Pd01: 3,558,715 bp) and shell width_25_ with AX-586014454 (Pd01: 3,421,529 bp). Pd02 included associations of shell width_25_ with AX-586052397 (Pd02: 19,206,185 bp), shell width_75_ with AX-586053062 (Pd02: 20,040,129 bp), and shell length with AX-586059241 (Pd02: 25,341,803 bp). On Pd05, a multitrait region around AX-586093174 (Pd05: 4,251,871 bp) was associated with shell area, shell width, shell width_25_, and shell perimeter, as also observed for kernel morphology. Shell shape associations were more broadly distributed across the genome, with signals detected on most chromosomes. Relevant associations included shell length-width ratio with AX-586031046 (Pd01: 31,012,190 bp), shell EF-PC3 with AX-586088684 (Pd04: 16,640,153 bp), and shell EF-PC1 with AX-586106712 (Pd06: 2,069,716 bp).

Associations for KS ratios were also widely distributed. However, one of the main multitrait regions was observed on Pd01 around AX-586036742 (Pd01: 38,664,451 bp), which was associated with several ratio traits, including KS circularity, KS ellipse ratio, KS length-width ratio, KS length-width_25_ ratio, KS length-width_75_ ratio, and KS width_75_. Additional important regions were detected on Pd02, particularly around AX-586054062 (Pd02: 21,686,838 bp), which was associated with KS area, KS width, and KS width_25_, and on Pd07 around AX-586132945 (Pd07: 16,511,360 bp), which was associated with KS width and KS width_25_. Weight-related traits showed an important region on Pd02. The strongest signal corresponded to KS weight with AX-586052448 (Pd02: 19,329,225 bp), and the same marker was also associated with shell weight. In contrast, kernel weight showed a relevant association on chromosome 6 with AX-586119655 (Pd06: 24,013,284 bp). For predicted quality traits, associations were relatively distributed throughout the genome, although Pd05 showed a notable concentration of signals. The strongest quality-related association was detected for predicted fiber with AX-586101877 (Pd05: 15,923,656 bp), and this region was also associated with predicted fats and predicted protein.

The effects caused by the significant SNPs identified in the GWAS were explored using SnpEff, detecting 7,198 predicted effects in total (Supplementary Tables 6 and 7). Most effects were classified as modifier impact variants (98.29%), while 84 effects showed low impact and 39 showed moderate impact. Regarding genomic location, most of the effects corresponded to downstream gene variants (43.39%) and upstream gene variants (41.91%). Less frequent but functionally relevant categories included 129 exonic effects (1.79%), seven splice-site region effects (0.10%), 41 effects in 3^*′*^ UTRs (0.57%) and 10 effects in 5^*′*^ UTRs (0.14%).

Functional annotations of genes close to significant markers were explored (Supplementary Table 8), revealing annotations for 2,488 genes. Candidate genes with functional annotations of biological relevance for the traits studied were identified and summarised in Table 2 and Figures 3, and 4. The candidate genes identified were associated with several functional categories of biological relevance to the analysed traits, particularly transcriptional regulation, cell-wall organisation and remodelling, ubiquitin-mediated protein degradation, signalling, and primary and lipid metabolism. Several transcriptional regulators were located near significant markers, including AP2/ERF, BES1/BZR1, bHLH, BIM2, NAC, FEZ-like/NAC-domain, and OVATE family proteins. A second prominent group comprised genes involved in cell-wall structure, biosynthesis, and modification, including wall-associated kinases, pectin methylesterases, fasciclin-like arabinogalactan proteins, a hydroxyproline O-galactosyltransferase, a cell-wall glycosyltransferase, and a WAT1-related transporter. Genes related to protein turnover and regulatory processes were also identified, including BIG BROTHER-like and RING-type E3 ubiquitin ligases and RING-H2 finger ATL proteins. Additional candidates were associated with metabolic functions, such as COMT-like O-methyltransferase, pyruvate dehydrogenase, HAL3-like PPC decarboxylase, GDSL lipase/esterase, cysteine synthase, and aspartic peptidase.

**Figure 3.**
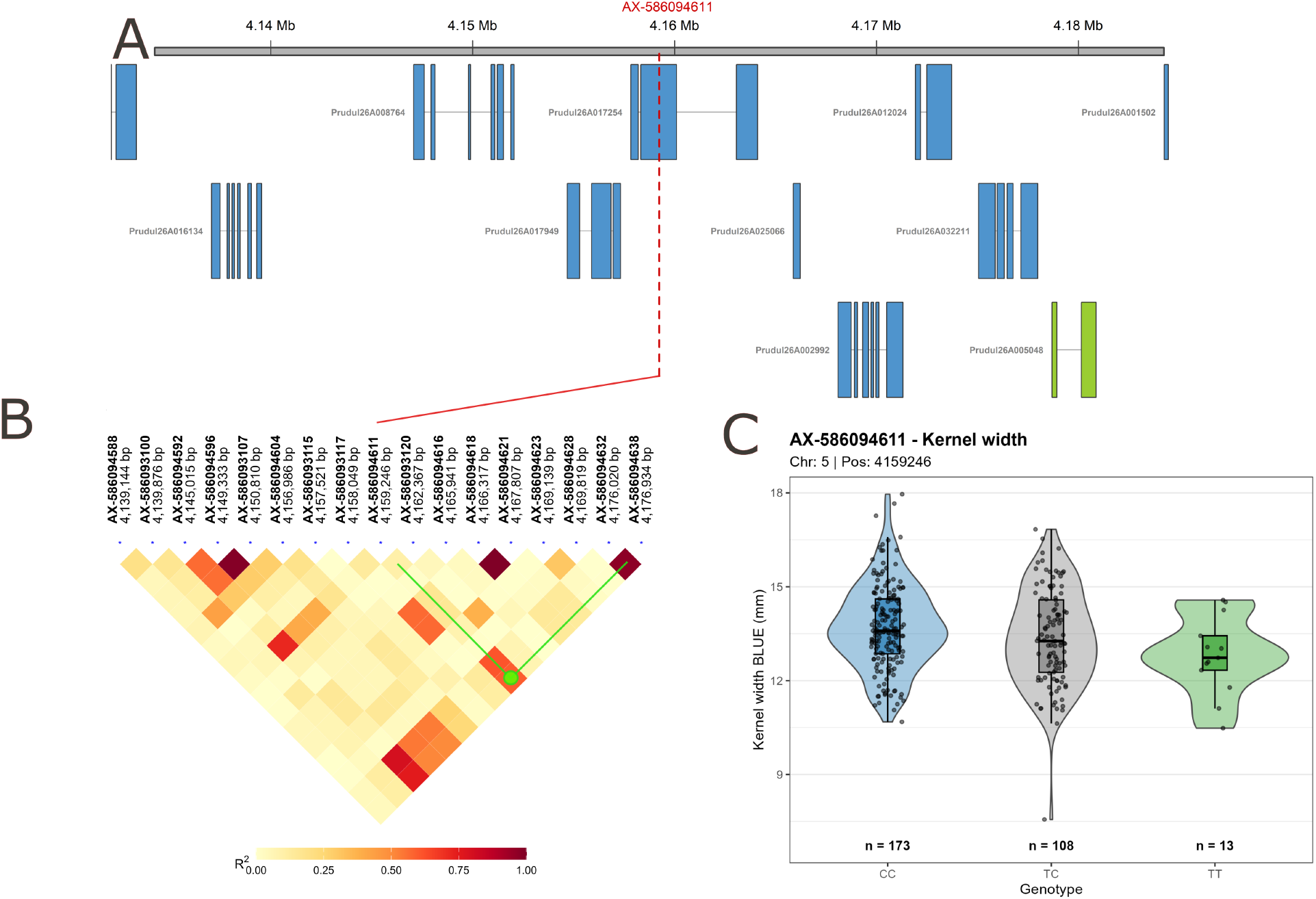
Candidate region for kernel width on chromosome 5 of Prunus dulcis. Panel A shows the genomic context, with the green gene representing the candidate gene; panel B shows that the marker closest to the candidate gene is in high linkage disequilibrium with AX-586094611. Panel C shows the boxplot/violin plot of kernel width BLUEs by genotype at AX-586094611.

**Table 2.**
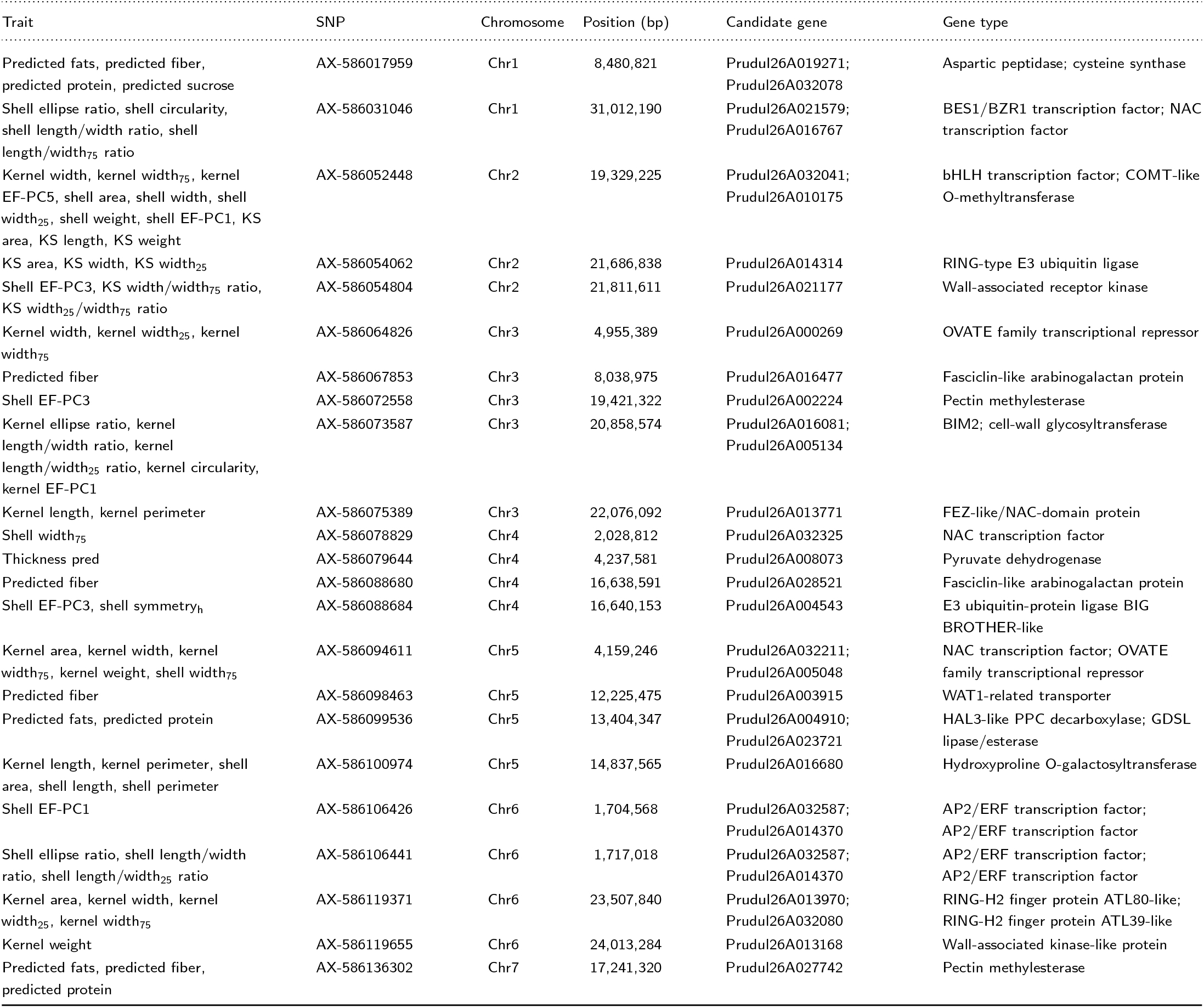
Candidate genes associated with kernel, shell, and predicted quality traits.

| Trait | SNP | Chromosome | Position (bp) | Candidate gene | Gene type |
| --- | --- | --- | --- | --- | --- |
| Predicted fats, predicted fiber, predicted protein, predicted sucrose | AX-586017959 | Chr1 | 8,480,821 | Prudul26A019271;<br>Prudul26A032078 | Aspartic peptidase; cysteine synthase |
| Shell ellipse ratio, shell circularity, shell length/width ratio, shell length/width<sub>75</sub> ratio | AX-586031046 | Chr1 | 31,012,190 | Prudul26A021579;<br>Prudul26A016767 | BES1/BZR1 transcription factor; NAC transcription factor |
| Kernel width, kernel width<sub>75</sub>, kernel EF-PC5, shell area, shell width, shell width<sub>25</sub>, shell weight, shell EF-PC1, KS area, KS length, KS weight | AX-586052448 | Chr2 | 19,329,225 | Prudul26A032041;<br>Prudul26A010175 | bHLH transcription factor; COMT-like O-methyltransferase |
| KS area, KS width, KS width<sub>25</sub> | AX-586054062 | Chr2 | 21,686,838 | Prudul26A014314 | RING-type E3 ubiquitin ligase |
| Shell EF-PC3, KS width/width<sub>75</sub> ratio, KS width<sub>25</sub>/width<sub>75</sub> ratio | AX-586054804 | Chr2 | 21,811,611 | Prudul26A021177 | Wall-associated receptor kinase |
| Kernel width, kernel width<sub>25</sub>, kernel width<sub>75</sub> | AX-586064826 | Chr3 | 4,955,389 | Prudul26A000269 | OVATE family transcriptional repressor |
| Predicted fiber | AX-586067853 | Chr3 | 8,038,975 | Prudul26A016477 | Fasciclin-like arabinogalactan protein |
| Shell EF-PC3 | AX-586072558 | Chr3 | 19,421,322 | Prudul26A002224 | Pectin methylesterase |
| Kernel ellipse ratio, kernel length/width ratio, kernel length/width<sub>25</sub> ratio, kernel circularity, kernel EF-PC1 | AX-586073587 | Chr3 | 20,858,574 | Prudul26A016081;<br>Prudul26A005134 | BIM2; cell-wall glycosyltransferase |
| Kernel length, kernel perimeter | AX-586075389 | Chr3 | 22,076,092 | Prudul26A013771 | FEZ-like/NAC-domain protein |
| Shell width<sub>75</sub> | AX-586078829 | Chr4 | 2,028,812 | Prudul26A032325 | NAC transcription factor |
| Thickness pred | AX-586079644 | Chr4 | 4,237,581 | Prudul26A008073 | Pyruvate dehydrogenase |
| Predicted fiber | AX-586088680 | Chr4 | 16,638,591 | Prudul26A028521 | Fasciclin-like arabinogalactan protein |
| Shell EF-PC3, shell symmetry<sub>h</sub> | AX-586088684 | Chr4 | 16,640,153 | Prudul26A004543 | E3 ubiquitin-protein ligase BIG BROTHER-like |
| Kernel area, kernel width, kernel width<sub>75</sub>, kernel weight, shell width<sub>75</sub> | AX-586094611 | Chr5 | 4,159,246 | Prudul26A032211;<br>Prudul26A005048 | NAC transcription factor; OVATE family transcriptional repressor |
| Predicted fiber | AX-586098463 | Chr5 | 12,225,475 | Prudul26A003915 | WAT1-related transporter |
| Predicted fats, predicted protein | AX-586099536 | Chr5 | 13,404,347 | Prudul26A004910;<br>Prudul26A023721 | HAL3-like PPC decarboxylase; GDLS lipase/esterase |
| Kernel length, kernel perimeter, shell area, shell length, shell perimeter | AX-586100974 | Chr5 | 14,837,565 | Prudul26A016680 | Hydroxyproline O-galactosyltransferase |
| Shell EF-PC1 | AX-586106426 | Chr6 | 1,704,568 | Prudul26A032587;<br>Prudul26A014370 | AP2/ERF transcription factor; AP2/ERF transcription factor |
| Shell ellipse ratio, shell length/width ratio, shell length/width<sub>25</sub> ratio | AX-586106441 | Chr6 | 1,717,018 | Prudul26A032587;<br>Prudul26A014370 | AP2/ERF transcription factor; AP2/ERF transcription factor |
| Kernel area, kernel width, kernel width<sub>25</sub>, kernel width<sub>75</sub> | AX-586119371 | Chr6 | 23,507,840 | Prudul26A013970;<br>Prudul26A032080 | RING-H2 finger protein ATL80-like; RING-H2 finger protein ATL39-like |
| Kernel weight | AX-586119655 | Chr6 | 24,013,284 | Prudul26A013168 | Wall-associated kinase-like protein |
| Predicted fats, predicted fiber, predicted protein | AX-586136302 | Chr7 | 17,241,320 | Prudul26A027742 | Pectin methylesterase |

**Figure 4.**
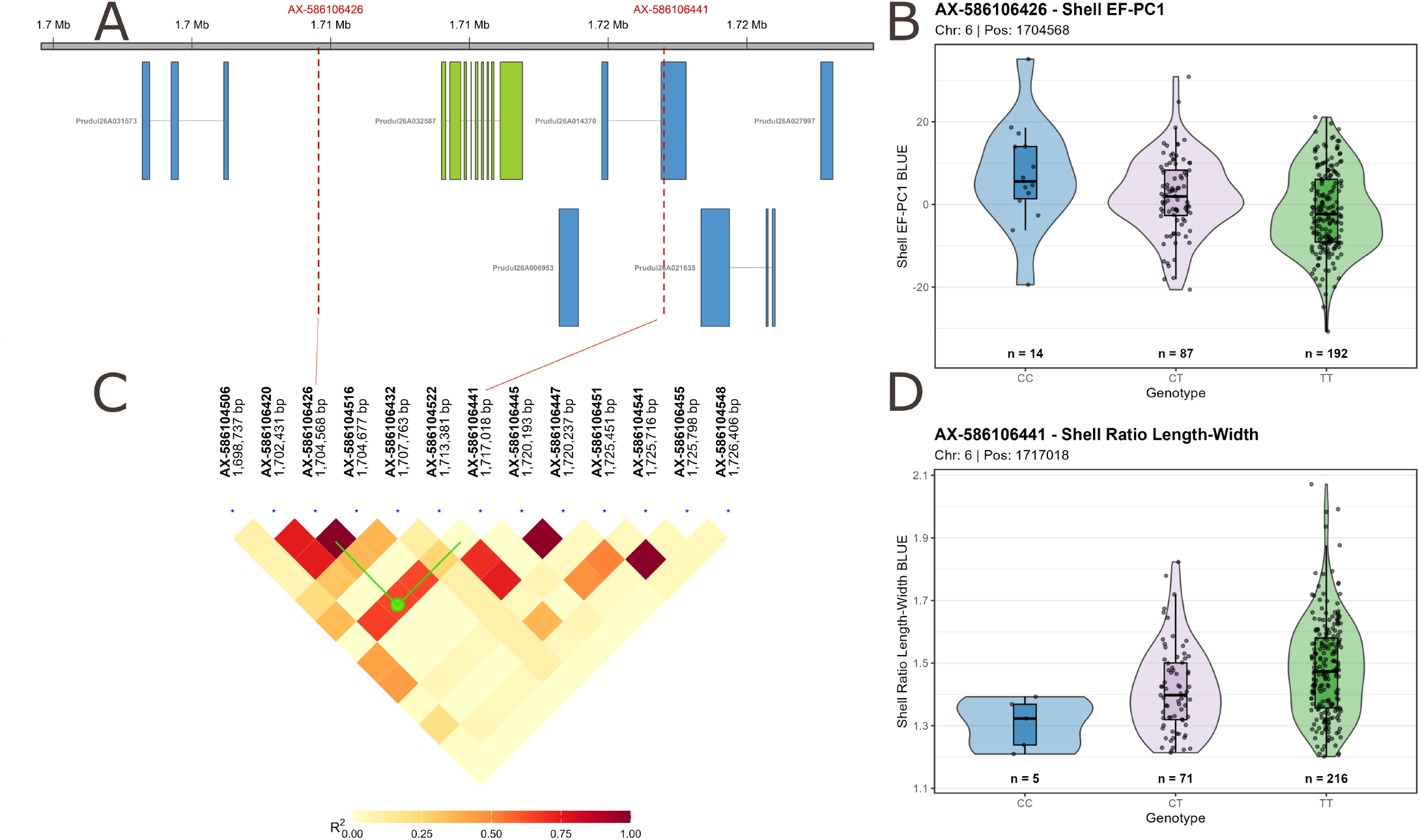
Candidate region for kernel aspect ratio related traits on chromosome 6. Panel A shows the genomic context, with the green gene representing the candidate gene; panel C shows that the markers closing the candidate gene are in high linkage disequilibrium (green lines closing region). Panel B and D shows the boxplot/violin plot of Shell EF-PC1 BLUEs and Shell Ratio Length-Width by genotype at AX-586106426 and AX-586106441, respectively.

## Discussion

The phenomics tools implemented in this work have boosted genomic insights into almond morphology and quality traits. The combination of high-throughput phenotyping and genotyping maximizes efficiency in both areas and has enabled the massive characterization of germplasm collections and breeding families at both the genotypic and phenotypic levels. This has allowed us to generate a large dataset of results, revealing the heritability, genetic architecture, and candidate genes associated with traits that had been scarcely studied before, making this one of the largest genomic association datasets in almond.

The imaging techniques implemented here not only increased the number of individuals and almonds studied, but also expanded the number of traits evaluated. In RGB imaging, the morphometric approach based on elliptic Fourier descriptors revealed new shape traits in both kernel and shell. In both cases, aspect ratio was the main trait influencing PC1, as is commonly observed in fruit morphometric studies [15, 37, 38]. However, additional EF-PC traits were also interpretable, such as kernel EF-PC3, which reflects a pronounced pointed tip for negative values; kernel EF-PC4, whose negative values indicate almond asymmetry on one side; and kernel EF-PC5, for which positive values correspond to almonds with a more rounded top. Indeed, strong and meaningful correlations were found between kernel EF-PC3 and kernel width_25_–width_75_ ratio (r = -0.782), reinforcing the idea that this component captures variation in width proportions relative to the tip. Similarly, kernel EF-PC4 was correlated with kernel shoulder symmetry (r = 0.586) and kernel symmetry_h_ (r = 0.565), supporting its association with almond asymmetry. Finally, kernel EF-PC5 showed a strong correlation with kernel width_25_–width ratio (r = 0.702), highlighting the influence of the upper part of the kernel on this component. In hyperspectral imaging, the most accurate predictions were obtained for the major compositional traits, fat and protein content, for which the PLS predictive models achieved low prediction errors and higher accuracies than in previous genomic studies using hyperspectral technologies (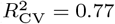 and 0.76, respectively) [39– 41]. Although the predictive accuracy for fiber and sucrose was lower than that obtained for the major components, these models can still provide valuable information for generating genomic insights and, especially, for supporting selection in breeding programs [42].

Phenotypic data analyses revealed substantial genotype and environment effects for the traits evaluated. Environmental effects were particularly pronounced for predicted thickness, kernel weight, and kernel width-related traits, reaching up to 42% for predicted thickness. These traits showed a clear decrease in the Santomera 2023 environment, which corresponded to a drier year than the other Santomera environments. This reduction was likely associated with altered kernel filling under drought conditions [43]. On the other hand, genotype effects were particularly relevant for highly heritable traits such as KS weight and shell weight, which are expected to be strongly genetically determined due to their association with shell hardness [10]. These were followed by aspect ratio-related traits, including kernel/shell EF-PC1, kernel length-width ratio, and KS ellipse ratio, which have previously been reported to exhibit high heritability in almond [44]. In addition, KS ratios emerged as less explored but highly relevant traits, also showing strong genetic determination. In particular, KS ratios related to symmetry_h_, length, width, and area exhibited heritability estimates higher than 0.64. These results suggest that, despite the high correlations observed between kernel and shell morphological traits, such as the correlation between kernel and shell length (r = 0.84), genetic variation may influence their relative proportions. This could reflect the presence of loci with differential effects on kernel and shell morphology, contributing to variation in their relative proportions despite their coordinated development [43]. Strong correlations were also observed among quality traits, particularly between protein and fat content, consistent with previous studies [8, 16]. These traits showed moderate-to-high broad-sense heritability values, ranging from 0.47 to 0.55, in agreement with previous estimates in almond [16, 45].

Genotyping of the studied accessions using the Axiom Almond 60K SNP array [46] was successful, yielding SNP datasets comparable to those reported in previous studies [11, 47]. This high-quality SNP dataset enabled population structure analysis to be successfully incorporated into the GWAS, allowing the identification of clusters according to genetic origin in the breeding families and geographical origin in the germplasm accessions. The population structure results were consistent with previous studies, which reported, for example, the clustering of USA accessions with French accessions [11]. They also agreed with pedigree records from the breeding program, as well as with historical records [48].

The macro-GWAS provided insights into the genetic architecture of multiple traits, generally supporting their quantitative genetic nature. For kernel morphology traits, a hotspot associated with kernel area and width-related traits was detected on chromosome 5, around 4.2 Mbp. This region has not previously been linked to kernel morphology. Although the hotspot was associated with several width-related traits, the strongest association was specifically observed for Width_75_, suggesting a potential role of this genomic region in kernel pointedness. This trait has not been explicitly measured in previous studies, but image-based phenomics tools enabled its detection and revealed a candidate genomic region of interest. Another kernel morphology trait showing a strong association, supported by six GWAS methods, was predicted thickness. This association was located at the beginning of chromosome 4, at 4,237,581 bp, close to a genomic region where a QTL was previously detected in an F1 population derived from the cross Marcona × Marinada [8]. Robust associations for kernel length were also identified on several chromosomes. The signal supported by the highest number of methods was located on chromosome 3, around 22.1 Mbp. In addition, an association detected on chromosome 5, around 14.8 Mbp, may be consistent with a kernel length QTL previously identified in the second half of chromosome 5 in an F1 population from the Vivot × Blanquerna cross [6]. Kernel shape showed two main association hotspots, one on chromosome 2 and another on chromosome 7. On chromosome 2, two subregions showed associations around 20 Mbp and 24 Mbp, both related to kernel aspect ratio traits, whereas the hotspot on chromosome 7 was located around 16.4 Mbp. QTLs related to kernel roundness have previously been detected on chromosomes 2 and 7, although they spanned large genomic regions, with peak positions located at 15.9 Mbp and 21.4 Mbp, respectively [8]. Kernel morphometric traits also showed robust associations supported by five GWAS methods, including EF-PC4 on chromosome 7 at 15.2 Mbp and EF-PC5 on chromosome 4 at 4.3 Mbp. These results provide further evidence for the genetic control of these traits, beyond their moderate heritability estimates presented previously.

The hotspots detected for shell morphology on chromosomes 1, 2, and 5 are novel, as the genomic basis of these traits has been scarcely investigated [6]. Some of these regions overlapped with hotspots identified for kernel morphology. For instance, the region on chromosome 2 around 20 Mbp was associated not only with kernel aspect ratio traits, but also with shell width_75_, which was detected by five GWAS methods and showed a mean LOD value above 7 (AX-586053062). Similarly, the chromosome 5 hotspot around 4.2 Mbp, previously associated with kernel area and width-related traits, also showed robust associations with shell area and width. While most associations detected for shell morphology appear to be novel, among shell shape traits, a region previously detected as a QTL for nut aspect ratio by Fernández i Martí et al. (2013) [6] may coincide with the association region identified in our analysis for shell length–width ratio on chromosome 1, around 31 Mbp. In addition, several novel associations were detected for shell shape traits, including shell EF-PC4 on chromosome 1 around 41 Mbp and shell EF-PC1 at the beginning of chromosome 6. Regarding KS morphology traits, these traits have not previously been explored at the genomic level. Therefore, the widespread hotspots identified here represent novel genomic regions and provide evidence for their genetic control. Together with the high heritability values observed for KS traits, these results support the existence of loci with differential effects on kernel and shell morphology.

Weight-related traits have been more extensively explored at the genomic level. The major region controlling KS weight (crackout) and shell weight has consistently been located on Pd02, although its precise position varies among studies. In a previous work, Pérez de los Cobos et al. (2023) [11] identified the most significant marker at 19,131,932 bp, whereas Sideli et al. (2023) [10] reported the strongest association at 23,264,396 bp (after mapping the primers to the Texas v2 genome). Here, the main association was detected at 19,329,225 bp, (AX-586052448) with a second signal at 19,206,185 bp (AX-586052397), closer to the position reported by Pérez de los Cobos et al. (2023) [11]. However, this does not correspond to the exact same fine-mapped region or marker. Indeed, the marker reported by Pérez de los Cobos et al. (2023) [11] at this position, AX-586051816, was removed from our dataset because it generated 33 mendelian errors. Thus, the germplasm context appears to influence fine mapping resolution, suggesting that integrating complementary approaches, such as gene expression analyses, could help disentangle this region. Regarding kernel weight, a robust association was identified at chromosome 6 around 24 Mbp, which had not been previously detected. Moreover, regions on chromosome 4 at 19.4 Mbp, chromosome 7 at 611 kbp, and chromosome 5 at 4.15 Mbp showed strong and robust associations supported by four or more GWAS methods. For the regions on chromosome 4 and at the beginning of chromosome 7, QTLs on low-density maps had been previously reported supporting these findings [6, 12]. Finally, predicted quality traits showed strong associations on Pd05, around 15.8 Mbp, involving several traits, particularly strong for predicted fiber. Additional robust associations were detected on chromosome 7 around 17.2 Mbp for fat, fiber, and protein, as well as on chromosome 1 around 8.5 Mbp. The region identified on chromosome 5 was located relatively close to a previously reported QTL for protein content, centered at 17,547,713 bp [11]. Similarly, the protein-associated region detected at the distal end of chromosome 7 was located near a QTL identified in a previous low-density QTL mapping study [7]. However, overall, the associations detected here for almond quality traits appear to be novel, given the scarcity of genomic studies on these traits, especially for traits such as sucrose, which have not previously been explored at the genomic level in almond.

The genomic regions identified here revealed several candidate genes potentially related to the traits studied. Among them, many were associated with developmental regulation and organ morphology, including genes annotated as OVATE family transcriptional repressors. Members of the OVATE family are widely recognised as regulators of organ shape, particularly in fruits, where they influence cell division patterns and contribute to shape variation in different plant species [19, 20]. In the present study, OVATE-related candidates were found close to markers AX-586064826 on chromosome 3 at 4,955,389 bp, associated with Prudul26A000269, and AX-586094611 on chromosome 5 at 4,159,246 bp, associated with Prudul26A005048. Notably, these regions were associated with kernel width, suggesting a possible role of these OVATE candidate genes in its regulation. Another relevant candidate region was identified on chromosome Pd06, where two AP2-like ethylene-responsive transcription factors, Prudul26A032587 and Prudul26A014370, were located around 1.71–1.72 Mbp and associated with shell shape-related traits. Moreover, the two associated SNPs were a variant with a low predicted impact located within the coding region (AX-586106441), and an upstream variant (AX-586106426). AP2 transcription factors are well known for their roles in plant developmental morphogenesis [21]. Notably, these genes showed homology with Arabidopsis PLT genes, particularly PLT2, which is involved in auxin-mediated developmental patterning in root stem cells [49]. Thus, these genes represent interesting candidates for organ morphology regulation. In addition, several NAC-related transcription factor candidates were identified, including the FEZ-like/NAC-domain gene Prudul26A013771 (Chr3, 22.08 Mbp), annotated with plant organ development, and the NAC transcription factors Prudul26A016767 (Chr1, 31.01 Mbp), Prudul26A032325 (Chr4, 2.03 Mbp) and Prudul26A032211 (Chr5, 4.16 Mbp). Among these, Prudul26A032325 was annotated with seed development, whereas Prudul26A032211 and Prudul26A016767 showed functional terms related to cell division and auxin response, supporting their relevance as candidates for almond shell and kernel morphology. Notably, Prudul26A032325 was annotated as NAC25, which has been reported to regulate rice endosperm integrity and GA-mediated endosperm cell expansion during Arabidopsis seed germination [50, 51].

Another interesting candidate was Prudul26A021579 (Chr1, 31.00 Mbp), annotated as a BES1/BZR1 brassinosteroid-responsive transcription factor and located in a region associated with shell aspect ratio traits. Brassinosteroid signaling has been shown to regulate seed size and shape in Arabidopsis by modulating integument, endosperm and embryo developmental pathways through BZR1 [22]. Similarly, in soybean, GmBZR1 has been linked to increased seed weight and size through enhanced integument cell expansion [52]. Also in relation to brassinosteroid pathways, Prudul26A016081 (Chr3, 20.86 Mbp), located in a region associated with kernel aspect ratio traits, was annotated as BIM2, a bHLH transcription factor related to BR signaling. BIM proteins have been implicated in fruit growth regulation, pericarp cell expansion and embryonic patterning [23, 53]. An additional brassinosteroid-related candidate gene (Prudul26A032041) was identified in a major hotspot around AX-586052448 (Chr2, 19.33 Mbp), associated with a broad set of kernel and shell morphology traits, including widths, aspect ratio, and weight-related traits. Prudul26A032041 is annotated as bHLH67 and contains FAMA/SPEECHLESS/MUTE-like bHLH domains. Specifically, bHLH67 orthologous transcription factors in rice have been shown to regulate grain morphology and cell expansion through brassinosteroid signaling [54, 55].

The previously mentioned hotspot marker AX-586052448 on chromosome 2, located around 19.3 Mbp, was also reported as the major-effect marker for shell weight (close to shell hardness) [10, 11]. In our analysis, this region was associated with a candidate gene (Prudul26A010175) annotated as a COMT-like O-methyltransferase, showing functional annotations related to methyltransferase activity and the lignin biosynthetic process. COMT is involved in the methylation steps of the monolignol biosynthetic pathway, and its down-regulation can alter lignin composition by promoting the incorporation of 5-hydroxyconiferyl alcohol into lignin [56]. The role of COMT in lignin biosynthesis has been studied in Arabidopsis, where COMT down-regulation combined with F5H1 overexpression strongly altered lignin composition [56]. Similarly, in *Leucaena leucocephala*, reduced COMT activity was reported to decrease lignin content by 28% [57], supporting the relevance of this enzyme in lignin deposition. Shell lignification has also been investigated in *Prunus armeniaca*, where COMT candidate genes on chromosome 2 were identified through QTL analyses and reported to be differentially expressed during shell development [18, 58, 59], further supporting the relevance of COMT-like genes in stone/shell lignification processes. However, the same apricot studies pointed to NST1 as the most plausible candidate based on transcriptomic analyses. Similarly, Pérez de los Cobos et al. (2023) [11] proposed Prudul26A013473, a NAC-domain gene homologous to NST1, as a candidate involved in shell hardness. NST1 has been described as a key regulator of secondary cell wall formation and has been shown to affect lignin accumulation [11, 18]. Based on the results obtained here, the identified COMT gene could be considered an additional candidate gene for further exploration, alongside the NST1 homolog previously reported in earlier studies.

Additional candidate genes related to cell wall formation were identified in genomic regions associated with the studied traits. Among them, a candidate gene annotated as a glycosyltransferase/rhamnogalacturonan I rhamnosyltransferase (Prudul26A005134) was located near the marker AX-586073587 on chromosome 3 at 20,858,574 bp, which was associated with kernel circularity. Glycosyltransferases are involved in the biosynthesis of pectins, major components of the primary cell wall that regulate its mechanical properties and facilitate wall deformation and expansive cell growth through changes in pectin composition and configuration [27, 60]. Additional pectin-related genes were identified (Prudul26A027742 and Prudul26A002224) near markers associated with predicted fiber and shell EF-PC3, respectively. Indeed, SNP AX-586136302 was located relatively close to Prudul26A027742, 1,070 bp upstream of the gene, and was classified as an upstream gene variant with a modifier effect. Pectin methylesterases modify the degree of pectin methylesterification, thereby regulating the mechanical properties of the cell wall and influencing cell expansion, adhesion and tissue morphology [28]. In addition, two wall-associated kinase genes (Prudul26A013168 and Prudul26A021177) were identified near markers associated with kernel weight and shell morphology traits. WAKs have been described as cell wall-associated receptors linked to the pectin fraction of the cell wall, acting as pectin receptors and participating in cell expansion during plant development [29]. Shifting from pectin-related components to cell wall-associated proteins, a candidate gene mainly associated with length-related shell and kernel morphology traits was also identified (Prudul26A016680), annotated as a hydroxyproline O-galactosyltransferase (GALT6). GALT6 catalyses the O-galactosylation of hydroxyproline-rich proteins (HRGPs), including arabinogalactan proteins (AGPs) which are involved in cell division, cell expansion, differentiation and plant organ development [30]. In Arabidopsis, mutant analyses have shown that GALT6 and related GALTs are required for proper AGP glycosylation and affect several growth- and development-related processes [61]. AGP-related genes were also detected in regions associated with predicted fiber content, including two fasciclin-like arabinogalactan proteins, Prudul26A016477 and Prudul26A028521, located near AX-586067853 and AX-586088680, respectively. Given the role of AGPs in cell expansion, differentiation and plant organ development [30], these genes are plausible candidates for fiber-related variation through their involvement in cell wall organization and tissue development. Another candidate gene related to fiber content was a WAT1-related transporter (Prudul26A003915), identified near a marker associated with predicted fiber content. WAT1 encodes a tonoplast-localized protein required for secondary cell wall formation in fibers, and wat1 mutants showed defective cell elongation and strongly reduced secondary wall deposition [31].

A group of candidate genes was related to protein ubiquitination, a regulatory mechanism with a well-established role in the control of seed and organ size [24, 25]. In Arabidopsis, the DA1 pathway restricts maternal seed growth by limiting cell proliferation in the integuments [24]. DA2 and EOD1/BIG BROTHER activate DA1 through monoubiquitination and act as negative regulators of seed and organ size [26]. Consistent with this regulatory mechanism, the SNP AX-586088684, associated with shell EF-PC3 and shell symmetry_v_, was located near Prudul26A004543, which encodes an E3 ubiquitin-protein ligase BIG BROTHER-like, with functional annotations of negative regulation of organ growth. In addition, SNP AX-586119371, associated with kernel area and several kernel-width parameters, identified two RING-H2 finger protein genes, Prudul26A013970 and Prudul26A032080, whereas SNP AX-586054062, associated with kernel-to-shell area and width ratios, was linked to the RING-type E3 ubiquitin ligase Prudul26A014314. Specifically, SNP AX-586054062 was located within the 5^*′*^ UTR of Prudul26A014314, 280 bp upstream of the coding sequence, and was predicted to have a modifier impact. Thus, these candidate genes suggest a possible role for E3 ubiquitin ligase activity in determining almond morphology.

Metabolism-related candidate genes were also identified in association with predicted kernel thickness and quality traits. The region tagged by AX-586099536 contained Prudul26A023721, encoding a GDSL lipase/esterase orthologous to Arabidopsis SFAR4. In Arabidopsis thaliana, SFAR4-like proteins have been shown to participate in fatty-acid degradation and to reduce fatty-acid storage in seeds, suggesting that this region could influence kernel fat content through lipid turnover [62, 63]. The same region also contained Prudul26A004910, encoding a HAL3-like PPC decarboxylase involved in CoA biosynthesis. Because CoA and acyl-CoA thioesters are essential components of fatty-acid metabolism, this gene may also affect lipid accumulation in the kernel [64]. Consistent with the potential involvement of acetyl-CoA metabolism in kernel lipid deposition, the region tagged by AX-586079644 on chromosome 4 contained Prudul26A008073, encoding a plastidial pyruvate dehydrogenase E1 *β* subunit. This enzyme may influence final kernel thickness because the pyruvate dehydrogenase complex converts pyruvate into acetyl-CoA, a key precursor for fatty-acid biosynthesis, thereby potentially affecting lipid deposition and kernel filling [33, 65].Finally, genes related to seed protein metabolism were identified. The marker AX-586017959, associated with predicted fat, fiber, protein, and sucrose contents, was located near Prudul26A019271, encoding an oryzasin-1-like aspartic peptidase. This protein may participate in the processing and mobilization of storage proteins, as disruption of the Arabidopsis aspartic protease ASPG1 delayed the degradation of 12S globulins during germination [34]. Its annotation in lipid metabolic processes and the presence of saposin-like domains also suggest a possible connection with lipid metabolism. The same marker was located near Prudul26A032078, encoding an O-acetylserine (thiol) lyase/cysteine synthase involved in cysteine biosynthesis. Consistently, overexpression of a cysteine synthase in soybean increased protein-bound cysteine and the accumulation of cysteine-rich seed proteins [35]. Variation in these protein metabolism genes could therefore affect seed protein composition and metabolism and indirectly influence the relative proportions of the remaining nutritional components.

## Conclusion

The integration of phenomics and genomics has enabled new insights into the genetic basis of relevant traits in almond, demonstrating its considerable potential for quantitative genetics and breeding. The high-throughput phenotyping approaches employed in this study facilitated the characterisation of traits whose genetic architecture had previously remained poorly understood because of the difficulties associated with their accurate and large-scale measurement. The results revealed genomic regions on chromosomes 2 and 5 that act as association hotspots for multiple kernel, shell, kernel-to-shell ratio, weight-related, and predicted quality traits. Together with the extensive dataset of marker–trait associations identified, these regions provide a valuable foundation for future genetic studies, marker validation, and breeding applications. The biological relevance of the detected associations was further supported by the identification of plausible candidate genes involved in transcriptional regulation, organ morphology, cell-wall development, protein turnover, and lipid metabolism, including genes encoding OVATE family proteins, bHLH and AP2/ERF transcription factors, RING-domain proteins, and GDSL lipases/esterases. Overall, this study provides genomic and phenomic resources that will contribute to a more comprehensive understanding of almond trait variation and support the development of improved cultivars through genomics-assisted breeding.

## Materials and methods

The overall computational workflow, including all layers from data acquisition to functional annotation, is summarized in Appendix 6: Figure 33.

### Plant material

Six F_1_ almond populations, a diversity germplasm panel, and a set of breeding selections were evaluated, comprising a total of 864 individuals (Supplementary Table 1). The F_1_ almond population comprised 696 individuals distributed across six full-sib families: Antoñeta’ × Marcona’ (37 individuals), Antoñeta’ × Penta’ (191 individuals), Antoñeta’ × Tardona’ (73 individuals), Florida’ × Marcona’ (63 individuals), Marcona’ × S4017’ (32 individuals), and R1000’ × Desmayo Largueta’ (300 individuals). The diversity germplasm panel consisted of 138 genotypes originating primarily from Spain, France, the USA, and Italy, while the breeding selection set comprised 30 individuals. Almond trees were evaluated at the CEBAS-CSIC experimental field in Santomera, Murcia, Spain, during 2022, 2023, and 2024. An additional environment was included in 2022 at the INRAE experimental field in France, comprising 67 individuals from the diversity germplasm panel studied, of which 21 were also present in the CEBAS-CSIC germplasm collection. The number of individuals evaluated in each year and the environment is summarized in Supplementary Table 2. For each genotype, 25 in-shell nuts per year were sampled whenever available and individual nut weight was recorded with and without the shell for Santomera environments.

### RGB Imaging

Almond samples were captured on a black-surface template using a Canon EOS 70D camera, achieving a resolution of 6 px/mm in 2022. For the 2022-France dataset and the 2023 and 2024 Santomera environment datasets, pseudo-RGB images were generated using the hyperspectral line-scan camera described in the next section, achieving a resolution of 6.81 px/mm. A detailed description of the complete image analysis workflow is provided in Más-Gómez et al. (2026) [15]. The workflow includes preprocessing, segmentation model development, deployment, morphological measurements, and morphometric analyses, implemented through interactive Python notebooks [15]. For the Santomera 2022 environment dataset captured using an RGB camera, the results have previously been described in Más-Gómez et al. (2026) [15]. For the rest of the environments captured with a line-scan camera, the workflow was updated by fine-tuning YOLO models [66] for kernel and in-shell images, based on pre-trained models on the COCO dataset for detection rather than segmentation. Subsequently, the detected almonds were segmented using the SAM model, with the centre of each bounding box provided as input, as shown in the GitHub repository. Erroneously detected almonds were interactively curated using the SAM model, and almonds that could not be correctly segmented, such as cut almonds in the image, were removed. For the detection model, pseudo-RGB images were split into training and validation sets: 44 and 21 images, respectively, for the kernel dataset and 30 and 17 images, respectively, for the shell dataset. The images were annotated using the CVAT labelling tool [67]. The fine-tuning process was performed for 100 epochs using a pre-trained YOLO detection model, YOLOV26s. Finally, the fine-tuned YOLO model, together with the rest of the workflow, was applied to the complete image dataset.

The complete set of traits extracted from the images by the workflow was divided into two groups: morphological measurements and shape measurements. Morphological measurements included length, defined as the longest axis of the fruit; width, defined as the dimension perpendicular to the length; area, defined as the surface enclosed by the contour of the fruit; perimeter, defined as the length of the outer boundary; width at three different positions along the length; and predicted thickness, estimated only for the kernels using a quadratic model based on area and weight, as described in Mas-Gómez et al. (2026) [15]. Shape measurements included aspect ratio, defined as the ratio of length to width and used as an indicator of elongation (length-width ratio); circularity, calculated as 4*π ×* area*/*perimeter^2^, with values closer to 1 indicating a more circular shape; ellipse ratio, defined as the ratio between the minor and major axes of a fitted ellipse; vertical and horizontal symmetry; symmetry in the upper part of the almond, used to quantify the shoulder; length-to-width ratios at 25% and 75% of the length; width-to-width ratios at 25% and 75% of the length; and morphometric traits. For morphometric traits, Elliptic Fourier Analysis (EFA) was performed using the Momocs v1.4.1 R package [68], followed by principal component analysis (PCA) of the Fourier coefficients, with the resulting Elliptic Fourier Principal Components (EF-PCs) used as traits describing shape variability. Finally, kernel-to-shell (KS) ratio traits were calculated for area, circularity, ellipse ratio, length, length-to-width ratio, length-to-width ratio at 25% and 75% of the length, width-to-width ratio at 25% and 75% of the length, width at 25% and 75% of the length, horizontal and vertical symmetry, and width.

### Hyperspectral Imaging

In Santomera 2023 and 2024 environments, almond samples were imaged using a hyperspectral line-scan camera (Black Industry SWIR 1.7 MAX, HAIP Solutions GmbH, Hannover, Germany) covering the 900–1730 nm spectral range, with 425 spectral bands and 1280 pixels per line. The camera was installed on a linear stage (X-LRT0750DL-BE08C, Zaber Technologies, Vancouver, Canada) to ensure controlled motion during acquisition. Sample illumination was achieved with a halogen bar light consisting of eight 20 W bulbs (SIMTRUM Photonics, Singapore). Images were processed following [16] using a Python workflow that included calibration, deep-learning-based segmentation using the masks obtained in the previous section, spectral preprocessing, and prediction of quality traits. Standard normal variate (SNV) preprocessing was applied, the spectral range was shortened to 950–1,668 nm, and the median spectrum was obtained for each genotype. In addition, to reduce the potential impact of sensor-related effects and differences in acquisition conditions between the 2023 and 2024 seasons (e.g., temperature variations), Dynamic Orthogonal Projection (DOP) was applied as a calibration transfer method using fat content as the response variable for the projection [69]. For this purpose, 17 samples scanned in 2024 were also analyzed to obtain reference quality values, following the procedures described in [16]. These 17 samples were subsequently used to project the Santomera 2023 and Reference samples using the DOP approach [69]. Different combinations of External Parameter Orthogonalization (EPO) components and Gaussian kernel bandwidth parameter (EPS) values were optimized by selecting the configuration that minimized the prediction error of the response variable within the 17-sample reference set. The code used to implement the DOP procedure is publicly available in a GitHub repository. Finally, Partial least squares (PLS) models were recalibrated using the 112 reference samples characterised in [16], to predict fat, fiber, protein, and sucrose contents, expressed as percentage (% dry weight basis), and scatterplots of the latent components were examined to assess the effectiveness of the DOP. The data were divided into calibration and external validation subsets following a 70:30 ratio. This partitioning was performed using the Kennard–Stone (KS) algorithm, ensuring a representative and well-spread selection of samples across the predictor space. Model optimization was carried out through repeated 8-fold cross-validation in the calibration set, performed 50 times.

### Phenotypic Data Analysis

Best linear unbiased estimates (BLUEs) were calculated for each trait using a linear mixed model fitted with the ASReml-R package. For each phenotypic trait, the model included the environment (location + year combination) and the genotype accession as fixed effects, as follows:

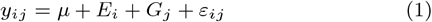

where *y*_*ij*_ is the phenotypic observation of the *j*-th almond accession evaluated in the *i*-th environment, *μ* is the overall mean, *E*_*i*_ is the fixed effect of the *i*-th environment, *G*_*j*_ is the fixed effect of the *j*-th genotype or accession, and *ε*_*ij*_ is the residual error term. Residuals were assumed to be independently and normally distributed as:

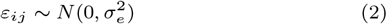

In addition, broad-sense heritability (H^2^) was estimated for each trait using a linear mixed model fitted with the ASReml-R package. For each phenotypic trait, the model included genotype accession and environment as random effects, as follows:

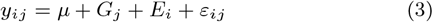

where *y*_*ij*_ is the phenotypic observation of the *j*-th almond accession evaluated in the *i*-th environment, *μ* is the overall mean, *G*_*j*_ is the random effect of the *j*-th genotype or accession, *E*_*i*_ is the random effect of the *i*-th environment, and *ε*_*ij*_ is the residual error term.

Broad-sense heritability was calculated as:

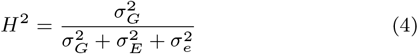

where 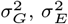, and 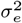 correspond to the genotypic, environmental, and residual variance components, respectively.

### Genotyping

A total of 359 non-redundant almond accessions were genotyped (Supplementary Table 1) using the almond Axiom™ 60K SNP array [46] at the Plateforme Gentyane UMR INRAE facilities in Clermont-Ferrand, France. Genotype calling and initial quality control were performed with `Axiom Analysis Suite v5.1.1.1`, following the recommended Best Practices Workflow. For downstream analyses, only SNP markers assigned to the “Poly High Resolution” (PHR) or “No Minor Homozygotes” (NMH) classes were retained, and markers with a minor allele frequency below 5% were discarded. Additional filtering was carried out to identify Mendelian inconsistencies using the *mendel* function implemented in `PLINK`, based on previously validated parent–offspring relationships. Mendelian errors were defined as alleles observed in the offspring but absent from both parents, thus violating Mendel’s first law. Markers showing inconsistencies in more than 2% of the tested parent–offspring relationships were excluded from the dataset, whereas markers involved in 2% or fewer inconsistencies were set as missing values for the corresponding individuals. In addition, for population structure analyses, linkage disequilibrium (LD) pruning was applied to reduce redundancy among markers. This filtering step was performed in `PLINK` using the `--indep-pairwise` command, with a sliding window of 50 SNPs, a step size of 5 SNPs, and an *r*^2^ threshold of 0.2.

### GWAS

The BLUEs of each accession were obtained from the fitted model and used as adjusted phenotypic values for downstream genome-wide association analyses (GWAS). Due to the potential artifacts and noise associated with morphometric descriptors derived from Elliptic Fourier Analysis, only EF-PC traits with a broad-sense heritability (H^2^) greater than 0.2 were retained for GWAS. Multi-locus genome-wide association analyses were performed using `mrMLM` v4.0 [70], which implements multi-locus GWAS approaches, including mrMLM, FASTmrMLM, FASTmrEMMA, pLARmEB, pKWmEB, ISIS EM-BLASSO, and EM-BLASSO. The analyses were conducted using the filtered genotype matrix, adjusted phenotypic values, a kinship matrix estimated from the marker data, and population structure inferred with `fastSTRUCTURE`. In total, 297 individuals were included for kernel and shell morphology traits, 253 for weight traits, and 245 for quality traits. Marker–trait associations were considered significant when they were detected by at least two independent methods and showed a LOD score of 3 or higher. Putative effects of the final SNP dataset on gene features were predicted using `SnpEff` v4.3e [71]. Variant annotation was performed against the *Prunus dulcis* cultivar ‘Texas’ v2.0 reference genome [1]. The SNP effects were classified by impact and type of variation. Candidate genes were identified by searching within a *±*25 kb window around significant SNPs using the *Prunus dulcis* ‘Texas’ v.2.0 genome annotation. Functional annotation was retrieved from Ensembl Plants using biomaRt, including gene descriptions, Gene Ontology terms, KEGG enzyme annotations, UniProt identifiers, InterPro domains and high-confidence *Arabidopsis thaliana* orthologs.

## Data availability

The code used for the imaging analyses is available in the almondcv2 and almondcv3 hyp GitHub repositories. Supplementary Tables are available on Zenodo.

## Author contributions statement

Conceptualization, J.M.G. and P.J.M.G.; Data curation, J.M.G.; Formal analysis, J.M.G.; Funding acquisition, P.J.M.G. and F.D.; Investigation, J.M.G., P.J.M.G. and M.R.; Methodology, J.M.G. and P.J.M.G.; Project administration, P.J.M.G. and F.D.; Resources, H.D., P.J.M.G., and F.D.; Software, J.M.G.; Supervision, P.J.M.G., F.D. and M.R.; Validation, P.J.M.G., F.D. and M.R.; Visualization, J.M.G.; Writing – original draft, J.M.G.; Writing – review and editing, J.M.G., H.D., P.J.M.G., M.R. and F.D.

## Acknowledgments

We would like to thank María del Mar Gómez Abajo, Camen Jurado, Francisco Gómez Lopez, Lucía Rodríguez Robles, Antonio Moreno Marín, Luis Miguel Serrano Sánchez, and Teresa Cremades Rosado for their assistance with the sampling. P.J.M.G. gratefully acknowledges the support of grant 22838/FLB/24, provided by Fulbright–Fundación Séneca, Región de Murcia (Spain).

## References

1. Tyler Alioto, Konstantinos G. Alexiou, Amélie Bardil, Fabio Barteri, Raúl Castanera, Fernando Cruz, Amit Dhingra, Henri Duval, Ángel Fernández i Martí, Leonor Frias, Beatriz Galán, José L. García, Werner Howad, Jèssica Gómez-Garrido, Marta Gut, Irene Julca, Jordi Morata, Pere Puigdomènech, Paolo Ribeca, María J. Rubio Cabetas, Anna Vlasova, Michelle Wirthensohn, Jordi Garcia-Mas, Toni Gabaldón, Josep M. Casacuberta, and Pere Arús. Transposons played a major role in the diversification between the closely related almond and peach genomes: results from the almond genome sequence. The Plant Journal, 101(2):455–472, 2020.

2. Jorge Mas-Gómez, Francisco José Gómez-López, Ángela Sánchez Prudencio, Manuel Rubio Angulo, and Pedro José Martínez-García. Accelerating Almond Breeding in Post-genomic Era, pages 159–166. Springer International Publishing, Cham, 2023.

3. Luís Felipe V. Ferrão, Rodrigo R. Amadeu, Juliana Benevenuto, Ivone de Bem Oliveira, and Patricio R. Munoz. Genomic selection in an outcrossing autotetraploid fruit crop: Lessons from blueberry breeding. Frontiers in Plant Science, Volume 12 - 2021, 2021.

4. R Socias, O Kodad, JM Alonso, and TM Gradziel. Almond quality: a breeding perspective. Horticultural reviews, 34:197– 238, 2007.

5. A. Verdú, S. Izquierdo, and R. Socias i Company. Processing and industrialization. CABI, page 460–481, 2017.

6. Angel Fernández i Martí, Carolina Font i Forcada, and Rafel Socias i Company. Genetic analysis for physical nut traits in almond. Tree genetics & genomes, 9(2):455–465, 2013.

7. Carolina Font i Forcada, Àngel Fernández i Martí, and Rafel Socias i Company. Mapping quantitative trait loci for kernel composition in almond. BMC genetics, 13(1):47, 2012.

8. Felipe Pérez de los Cobos, Agustí Romero, Leontina Lipan, Xavier Miarnau, Pere Arús, Iban Eduardo, Ignasi Batlle, and Alejandro Calle. Qtl mapping of almond kernel quality traits in the f1 progeny of ‘marcona’ × ‘marinada’. Frontiers in Plant Science, Volume 15 - 2024, 2024.

9. Shashi N Goonetilleke, Michelle G Wirthensohn, and Diane E Mather. Genetic analysis of quantitative variation in almond nut traits. Tree Genetics & Genomes, 19(6):55, 2023.

10. Gina M Sideli, Diane Mather, Michelle Wirthensohn, Federico Dicenta, Shashi N Goonetilleke, Pedro José Martínez-García, and Thomas M Gradziel. Genome-wide association analysis and validation with kasp markers for nut and shell traits in almond (prunus dulcis [mill.] da webb). Tree Genetics & Genomes, 19(2):13, 2023.

11. Felipe Pérez de los Cobos, Eva Coindre, Naima Dlalah, Bénédicte Quilot-Turion, Ignasi Batlle, Pere Arús, Iban Eduardo, and Henri Duval. Almond population genomics and non-additive gwas reveal new insights into almond dissemination history and candidate genes for nut traits and blooming time. Horticulture Research, 10(10):uhad193, 10 2023.

12. Raquel Sánchez-Pérez, Werner Howad, Federico Dicenta, Pere Arús, and P Martínez-Gómez. Mapping major genes and quantitative trait loci controlling agronomic traits in almond. Plant Breeding, 126(3):310–318, 2007.

13. Wanneng Yang, Hui Feng, Xuehai Zhang, Jian Zhang, John H Doonan, William David Batchelor, Lizhong Xiong, and Jianbing Yan. Crop phenomics and high-throughput phenotyping: past decades, current challenges, and future perspectives. Molecular plant, 13(2):187–214, 2020.

14. Mansoor Sheikh, Farooq Iqra, Hamadani Ambreen, Kumar A Pravin, Manzoor Ikra, and Yong Suk Chung. Integrating artificial intelligence and high-throughput phenotyping for crop improvement. Journal of Integrative Agriculture, 23(6):1787–1802, 2024.

15. Jorge Mas-Gómez, Manuel Rubio, Federico Dicenta, and Pedro José Martínez-García. Open rgb imaging workflow for morphological and morphometric analysis of fruits using deep learning: a case study on almonds. GigaScience, 15:giaf157, 01 2026.

16. Jorge Mas-Gómez, Manuel Rubio, Federico Dicenta, and Pedro José Martínez-García. Unlocking almond breeding for nutritional composition with hyperspectral imaging. Plant Phenomics, page 100239, 2026.

17. Robert T Furbank and Mark Tester. Phenomics–technologies to relieve the phenotyping bottleneck. Trends in plant science, 16(12):635–644, 2011.

18. Xiao Zhang, Lijie Zhang, Qiuping Zhang, Jiayu Xu, Weisheng Liu, and Wenxuan Dong. Comparative transcriptome profiling and morphology provide insights into endocarp cleaving of apricot cultivar (prunus armeniaca l.). BMC plant biology, 17(1):72, 2017.

19. Jiping Liu, Joyce Van Eck, Bin Cong, and Steven D. Tanksley. A new class of regulatory genes underlying the cause of pear-shaped tomato fruit. Proceedings of the National Academy of Sciences, 99(20):13302–13306, 2002.

20. Ashley Snouffer, Carmen Kraus, and Esther van der Knaap. The shape of things to come: ovate family proteins regulate plant organ shape. Current Opinion in Plant Biology, 53:98–105, 2020. Growth and development.

21. Kai Feng, Xi-Lin Hou, Guo-Ming Xing, Jie-Xia Liu, Ao-Qi Duan, Zhi-Sheng Xu, Meng-Yao Li, Jing Zhuang, and Ai-Sheng Xiong. Advances in ap2/erf super-family transcription factors in plant. Critical reviews in biotechnology, 40(6):750–776, 2020.

22. Wen-Bo Jiang, Hui-Ya Huang, Yu-Wei Hu, Sheng-Wei Zhu, Zhi-Yong Wang, and Wen-Hui Lin. Brassinosteroid regulates seed size and shape in arabidopsis. Plant physiology, 162(4):1965–1977, 2013.

23. Kentaro Mori, Martine Lemaire-Chamley, Joana Jorly, Fernando Carrari, Mariana Conte, Erika Asamizu, Tsuyoshi Mizoguchi, Hiroshi Ezura, and Christophe Rothan. The conserved brassinosteroid-related transcription factor bim1a negatively regulates fruit growth in tomato. Journal of Experimental Botany, 72(4):1181–1197, 2021.

24. Na Li, Ran Xu, and Yunhai Li. Molecular networks of seed size control in plants. Annual review of plant biology, 70(1):435–463, 2019.

25. Na Li and Yunhai Li. Signaling pathways of seed size control in plants. Current opinion in plant biology, 33:23–32, 2016.

26. Tian Xia, Na Li, Jack Dumenil, Jie Li, Andrei Kamenski, Michael W. Bevan, Fan Gao, and Yunhai Li. The ubiquitin receptor da1 interacts with the e3 ubiquitin ligase da2 to regulate seed and organ size in arabidopsis . The Plant Cell, 25(9):3347–3359, 09 2013.

27. Robert Palin and Anja Geitmann. The role of pectin in plant morphogenesis. Biosystems, 109(3):397–402, 2012. Biological Morphogenesis: Theory and Computation.

28. Jérôme Pelloux, Christine Rusterucci, and Ewa J Mellerowicz. New insights into pectin methylesterase structure and function. Trends in plant science, 12(6):267–277, 2007.

29. Bruce D. Kohorn and Susan L. Kohorn. The cell wall-associated kinases, waks, as pectin receptors. Frontiers in Plant Science, Volume 3 - 2012, 2012.

30. Anna Majewska-Sawka and Eugene A. Nothnagel. The multiple roles of arabinogalactan proteins in plant development. Plant Physiology, 122(1):3–10, 01 2000.

31. Philippe Ranocha, Nicolas Denancé, Ruben Vanholme, Amandine Freydier, Yves Martinez, Laurent Hoffmann, Lothar Köhler, Cécile Pouzet, Jean-Pierre Renou, Björn Sundberg, Wout Boerjan, and Deborah Goffner. Walls are thin1 (wat1), an arabidopsis homolog of medicago truncatula nodulin21, is a tonoplast-localized protein required for secondary wall formation in fibers. The Plant Journal, 63(3):469–483, 2010.

32. Hui Li, Jia Sun, Ying Zhang, Ning Wang, Tianshu Li, Huiying Dong, Mingliang Yang, Chang Xu, Limin Hu, Chunyan Liu, et al. Soybean oil and protein: biosynthesis, regulation and strategies for genetic improvement. Plant, Cell & Environment, 49(7):3730–3746, 2026.

33. Julius Ver Sagun, Umesh Prasad Yadav, and Ana Paula Alonso. Progress in understanding and improving oil content and quality in seeds. Frontiers in plant science, 14:1116894, 2023.

34. Wenzhong Shen, Xuan Yao, Tiantian Ye, Sheng Ma, Xiong Liu, Xiaoming Yin, and Yan Wu. Arabidopsis aspartic protease aspg1 affects seed dormancy, seed longevity and seed germination. Plant and Cell Physiology, 59(7):1415–1431, 07 2018.

35. Won-Seok Kim, Demosthenis Chronis, Matthew Juergens, Amy C Schroeder, Seung Won Hyun, Joseph M Jez, and Hari B Krishnan. Transgenic soybean plants overexpressing o-acetylserine sulfhydrylase accumulate enhanced levels of cysteine and bowman–birk protease inhibitor in seeds. Planta, 235(1):13–23, 2012.

36. Thermo Fisher Scientific. Axiom™ Genotyping Solution Data Analysis User Guide. Thermo Fisher Scientific, Waltham, MA, USA, 2020. Publication Number MAN0018363.

37. Huimin Wang, Hao Yin, Haitao Li, Gengchen Wu, Wei Guo, Kaijie Qi, Shutian Tao, Shaoling Zhang, Seishi Ninomiya, and Yue Mu. Quantitative 2d fruit shape analysis of a wide range of pear genetic resources toward shape design breeding. Scientia Horticulturae, 327:112826, 2024.

38. Bünyamin Demir, Bahadır Sayıncı, NECATİ Çetin, MEHMET Yaman, Ruçan Ç ömlek, Y Aydın, and M Sutyemez. Elliptic fourier based analysis and multivariate approaches for size and shape distinctions of walnut (juglans regia l.) cultivars. Grasas y Aceites, 69(4):e271–e271, 2018.

39. Marcin Grzybowski, Nuwan K Wijewardane, Abbas Atefi, Yufeng Ge, and James C Schnable. Hyperspectral reflectance-based phenotyping for quantitative genetics in crops: Progress and challenges. Plant Communications, 2(4), 2021.

40. Dawei Sun, Haiyan Cen, Haiyong Weng, Liang Wan, Alwaseela Abdalla, Ahmed Islam El-Manawy, Yueming Zhu, Nan Zhao, Haowei Fu, Juan Tang, et al. Using hyperspectral analysis as a potential high throughput phenotyping tool in gwas for protein content of rice quality. Plant methods, 15(1):54, 2019.

41. Hengbiao Zheng, Weijie Tang, Tao Yang, Meng Zhou, Caili Guo, Tao Cheng, Weixing Cao, Yan Zhu, Yunhui Zhang, and Xia Yao. Grain protein content phenotyping in rice via hyperspectral imaging technology and a genome-wide association study. Plant Phenomics, 6:0200, 2024.

42. Cheryl Dalid, Caiwang Zheng, Luis Osorio, Sujeet Verma, Amr Abd-Elrahman, Xu Wang, and Vance M Whitaker. Genetic analysis of predicted vegetative biomass and biomass-related traits from digital phenotyping of strawberry. The Plant Genome, 18(2):e70018, 2025.

43. David Doll and Kenneth Shackel. Drought management for california almonds. Crops & Soils, 49(2):28–35, 2016.

44. P.J. Martínez-García, M. Rubio, T. Cremades, and F. Dicenta. Inheritance of shell and kernel shape in almond (prunus dulcis). Scientia Horticulturae, 244:330–338, 2019.

45. Carolina Font i Forcada, O Kodad, T Juan, G Estopañan, et al. Genetic variability and pollen effect on the transmission of the chemical components of the almond kernel. Spanish journal of agricultural research, 9(3):781–789, 2011.

46. Henri Duval, Eva Coindre, Sebastian E. Ramos-Onsins, Konstantinos G. Alexiou, Maria J. Rubio-Cabetas, Pedro J. Martínez-García, Michelle Wirthensohn, Amit Dhingra, Anna Samarina, and Pere Arús. Development and evaluation of an axiomtm 60k snp array for almond (prunus dulcis). Plants, 12(2), 2023.

47. Jorge Mas-Gómez, Francisco José Gómez-López, Manuel Rubio, María del Mar Gómez-Abajo, Federico Dicenta, and Pedro José Martínez-García. Integration of linkage mapping, qtl analysis, rna-seq data, and genome-wide association studies (gwas) to explore relative flowering traits in almond. Horticultural Plant Journal, 2026.

48. Alejandro Calle, Lidia Aparicio-Durán, Ignasi Batlle, Iban Eduardo, and Xavier Miarnau. Review of agronomic and kernel quality traits of 273 almond cultivars. Genetic Resources and Crop Evolution, 72(4):3783–3828, 2025.

49. Luca Santuari, Gabino F Sanchez-Perez, Marijn Luijten, Bas Rutjens, Inez Terpstra, Lidija Berke, Maartje Gorte, Kalika Prasad, Dongping Bao, Johanna LPM Timmermans-Hereijgers, et al. The plethora gene regulatory network guides growth and cell differentiation in arabidopsis roots. The Plant Cell, 28(12):2937–2951, 2016.

50. Rong Li, Ming-Wei Wu, Jinxin Liu, Xintong Xu, Yiqun Bao, and Chun-Ming Liu. Nac25 transcription factor regulates the degeneration of cytoplasmic membrane integrity and starch biosynthesis in rice endosperm through interacting with mads29. Frontiers in Plant Science, 16:1563065, 2025.

51. Rocío Sánchez-Montesino, Laura Bouza-Morcillo, Julietta Marquez, Melania Ghita, Salva Duran-Nebreda, Luis Gómez, Michael J Holdsworth, George Bassel, and Luis Oñate-Sánchez. A regulatory module controlling ga-mediated endosperm cell expansion is critical for seed germination in arabidopsis. Molecular Plant, 12(1):71–85, 2019.

52. Xiang Lu, Qing Xiong, Tong Cheng, Qing-Tian Li, Xin-Lei Liu, Ying-Dong Bi, Wei Li, Wan-Ke Zhang, Biao Ma, Yong-Cai Lai, et al. A pp2c-1 allele underlying a quantitative trait locus enhances soybean 100-seed weight. Molecular Plant, 10(5):670–684, 2017.

53. John W Chandler, Melanie Cole, Annegret Flier, and Wolfgang Werr. Bim1, a bhlh protein involved in brassinosteroid signalling, controls arabidopsis embryonic patterning via interaction with dornröschen and dornröschen-like. Plant Molecular Biology, 69(1):57–68, 2009.

54. Hyoseob Seo, Suk-Hwan Kim, Byoung-Doo Lee, Jung-Hyun Lim, Sang-Ji Lee, Gynheung An, and Nam-Chon Paek. The rice basic helix–loop–helix 79 (osbhlh079) determines leaf angle and grain shape. International journal of molecular sciences, 21(6):2090, 2020.

55. Shouzhen Teng, Qiming Liu, Guoxin Chen, Yuan Chang, Xuean Cui, Jinxia Wu, Pengfei Ai, Xuehui Sun, Zhiguo Zhang, and Tiegang Lu. Osbhlh92, in the noncanonical brassinosteroid signaling pathway, positively regulates leaf angle and grain weight in rice. New Phytologist, 240(3):1066–1081, 2023.

56. Ruben Vanholme, John Ralph, Takuya Akiyama, Fachuang Lu, Jorge Rencoret Pazo, Hoon Kim, Jørgen Holst Christensen, Brecht Van Reusel, Véronique Storme, Riet De Rycke, Antje Rohde, Kris Morreel, and Wout Boerjan. Engineering traditional monolignols out of lignin by concomitant up-regulation of f5h1 and down-regulation of comt in arabidopsis. The Plant Journal, 64(6):885–897, 2010.

57. Smita Rastogi and Upendra Nath Dwivedi. Down-regulation of lignin biosynthesis in transgenic leucaena leucocephala harboring o-methyltransferase gene. Biotechnology Progress, 22(3):609–616, 2006.

58. Xiao Zhang, Qiuping Zhang, Xinyu Sun, Xiao Du, Weisheng Liu, and Wenxuan Dong. Differential expression of genes encoding phenylpropanoid enzymes in an apricot cultivar (prunus armeniaca l.) with cleavable endocarp. Trees, 33(6):1695–1710, 2019.

59. Qiuping Zhang, Yuping Zhang, Weisheng Liu, Ning Liu, Xiaoxue Ma, Chunjing Lü, Ming Xu, Shuo Liu, and Yujun Zhang. Re-sequencing and morphological data revealed the genetics of stone shell and kernel traits in apricot. Frontiers in Plant Science, Volume 14 - 2023, 2023.

60. Charles T. Anderson. We be jammin’: an update on pectin biosynthesis, trafficking and dynamics. Journal of Experimental Botany, 67(2):495–502, 01 2016.

61. Debarati Basu, Lu Tian, Wuda Wang, Shauni Bobbs, Hayley Herock, Andrew Travers, and Allan M Showalter. A small multigene hydroxyproline-o-galactosyltransferase family functions in arabinogalactan-protein glycosylation, growth and development in arabidopsis. BMC plant biology, 15(1):295, 2015.

62. Mingxun Chen, XUE Du, Yang Zhu, Zhong Wang, Shuijian Hua, Zhilan Li, Wangli Guo, Guoping Zhang, Jinrong Peng, and Lixi Jiang. Seed fatty acid reducer acts downstream of gibberellin signalling pathway to lower seed fatty acid storage in arabidopsis. Plant, cell & environment, 35(12):2155–2169, 2012.

63. Li-Min Huang, Chia-Ping Lai, Long-Fang O Chen, Ming-Tsair Chan, and Jei-Fu Shaw. Arabidopsis sfar4 is a novel gdsl-type esterase involved in fatty acid degradation and glucose tolerance. Botanical studies, 56(1):33, 2015.

64. Thomas Kupke, Pilar Hernandez-Acosta, Stefan Steinbacher, and Francisco A Culianez-Macia. Arabidopsis thaliana flavoprotein athal3a catalyzes the decarboxylation of 4′-phosphopantothenoylcysteine to 4′-phosphopantetheine, a key step in coenzyme a biosynthesis. Journal of Biological Chemistry, 276(22):19190–19196, 2001.

65. Jinshan Ke, Robert H. Behal, Stephanie L. Back, Basil J. Nikolau, Eve Syrkin Wurtele, and David J. Oliver. The role of pyruvate dehydrogenase and acetyl-coenzyme a synthetase in fatty acid synthesis in developing arabidopsis seeds. Plant Physiology, 123(2):497–508, 06 2000.

66. J. Redmon, S. Divvala, R. Girshick, and A. Farhadi. You Only Look Once: Unified, Real-Time Object Detection. In Proceedings of the IEEE Conference on Computer Vision and Pattern Recognition, pages 779–788, Las Vegas, NV, USA, 2016. IEEE.

67. B. Sekachev, N. Manovich, M. Zhiltsov, A. Zhavoronkov, D. Kalinin, B. Hoff, T. Osmanov, D. Kruchinin, A. Zankevich, D. Sidnev, M. Markelov Johannes, M. Chenuet, aandre, telenachos, A. Melnikov, J. Kim, L. Ilouz, N. Glazov Priya, R. Tehrani, S. Jeong, V. Skubriev, S. Yonekura, V. Truong, zliang, lizhming, and T. Truong. opencv/cvat: v1.1.0, 2020.

68. Vincent Bonhomme, Sandrine Picq, Cédric Gaucherel, and Julien Claude. Momocs: Outline analysis using r. Journal of Statistical Software, 56(13):1–24, 2014.

69. Puneet Mishra, Jean Michel Roger, Douglas N. Rutledge, and Ernst Woltering. Two standard-free approaches to correct for external influences on near-infrared spectra to make models widely applicable. Postharvest Biology and Technology, 170:111326, 2020.

70. Ya-Wen Zhang, Cox Lwaka Tamba, Yang-Jun Wen, Pei Li, Wen-Long Ren, Yuan-Li Ni, Jun Gao, and Yuan-Ming Zhang. mrmlm v4.0.2: An r platform for multi-locus genome-wide association studies. Genomics, Proteomics Bioinformatics, 18(4):481–487, 08 2020.

71. Pablo Cingolani, Adrian Platts, Le Lily Wang, Melissa Coon, Tung Nguyen, Luan Wang, Susan J. Land, Xiangyi Lu, and Douglas M. Ruden. A program for annotating and predicting the effects of single nucleotide polymorphisms, snpeff. Fly, 6(2):80–92, 2012. PMID: 22728672.

