## Appendix for "From Phenomics to Genomics: Macro-GWAS of Almond Morphology and Quality"

*Pre-print*, 2026, pp. 1–32
**Paper**

### From Phenomics to Genomics: Macro-GWAS of Almond Morphology and Quality

Jorge Mas-Gómez,[1,*] Manuel Rubio,[1] Henri Duval,[2] Federico Dicenta[1] and Pedro José Martínez-García[1,*]

[1]Fruit Breeding Group, CEBAS-CSIC, Campus Universitario Espinardo, 30100, Murcia, Spain
[2]Unité de Génétique et Amélioration des Fruits et Légumes (GAFL), INRAE, F-84143, Montfavet, France

### Appendix 1

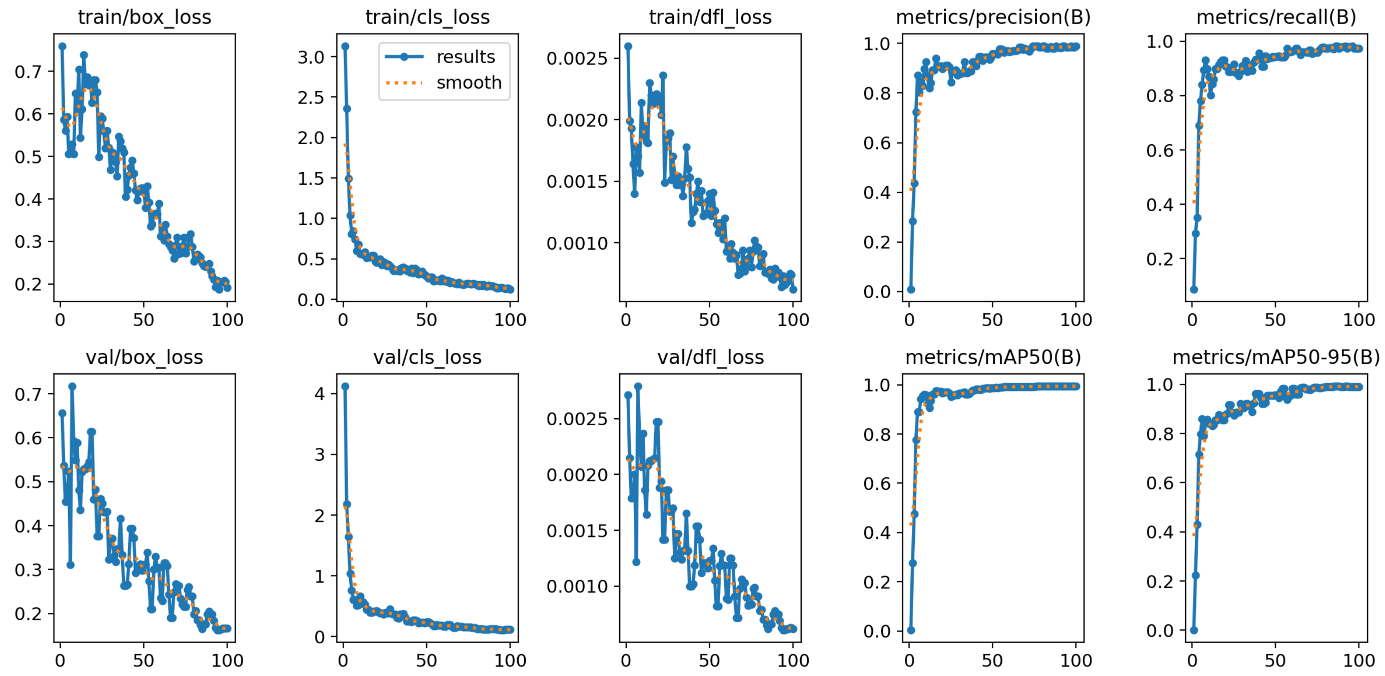


**Figure 1** Training results for kernel detection.

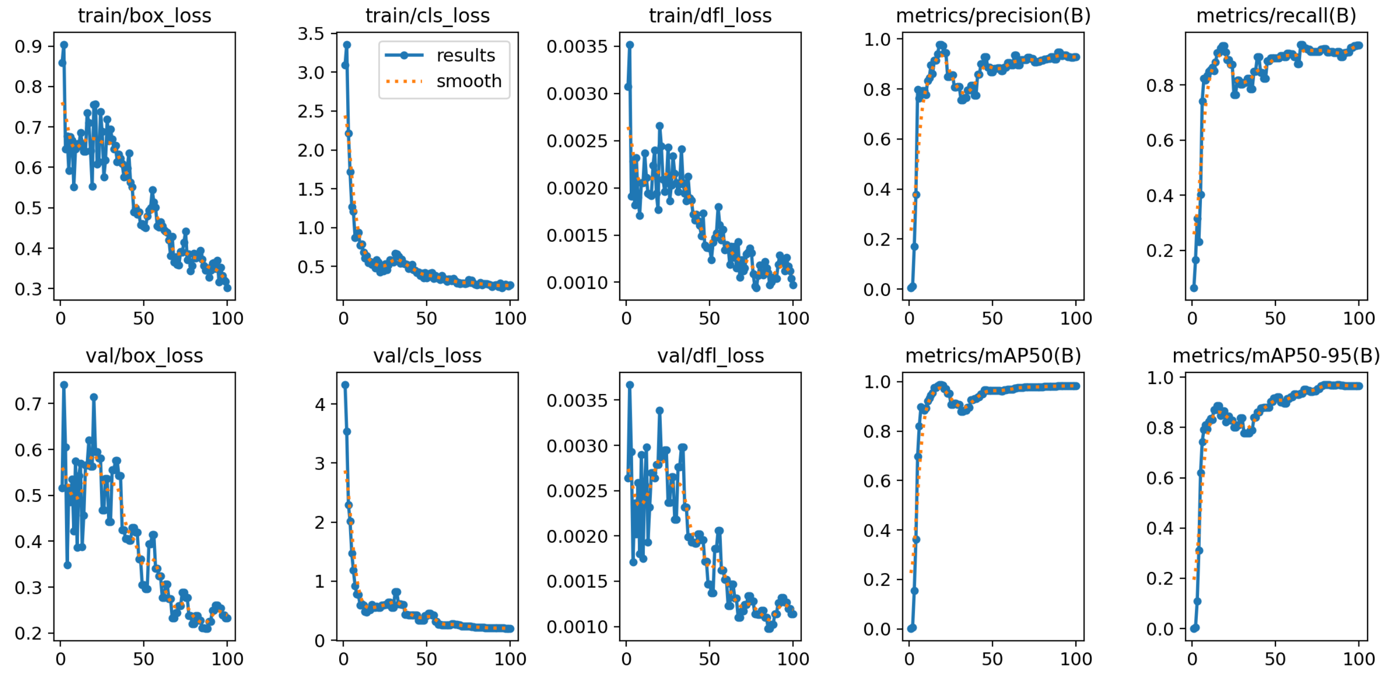


**Figure 2** Training results for shell detection.

#### Appendix 2

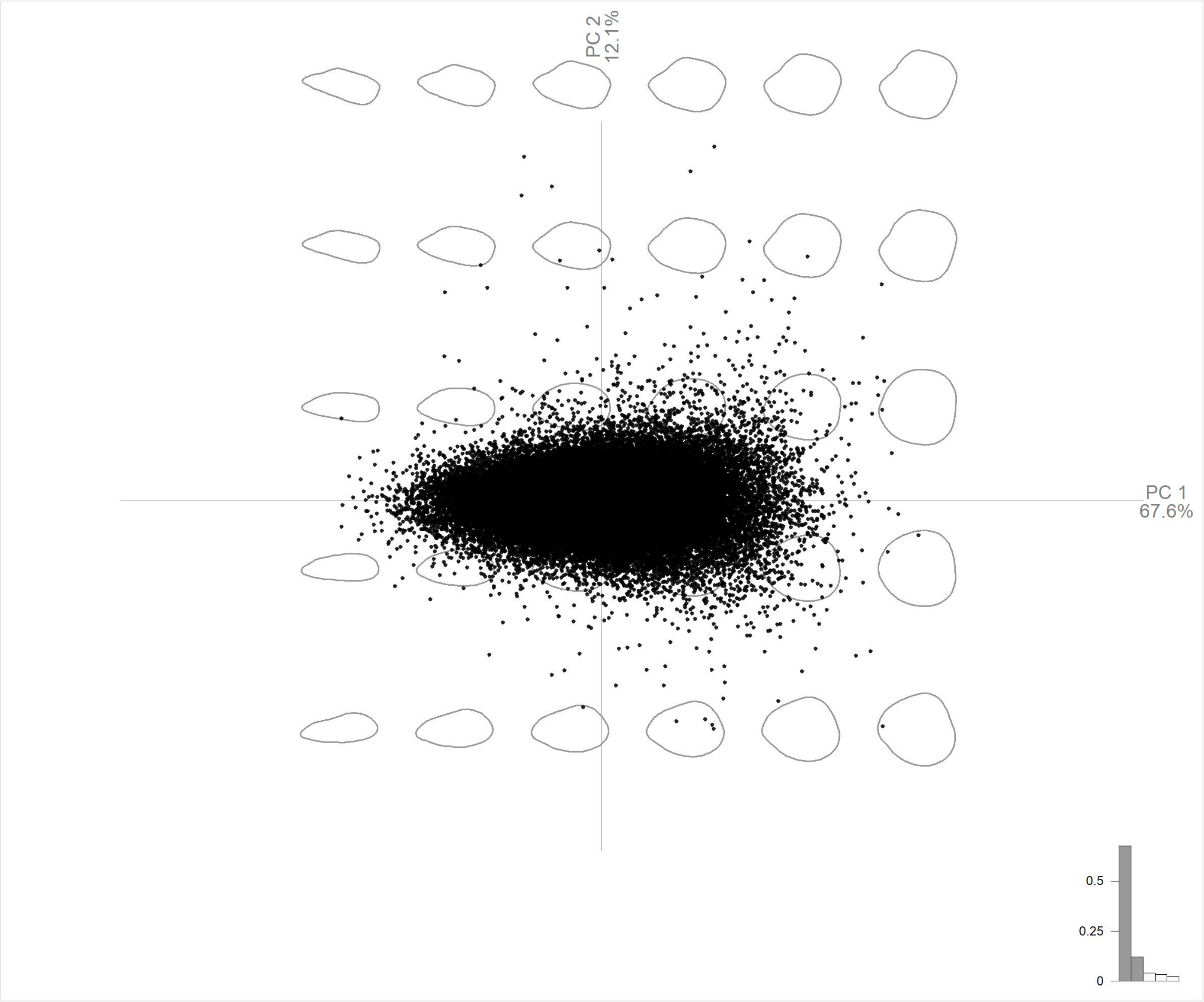


**Figure 3** Training results for shell detection.

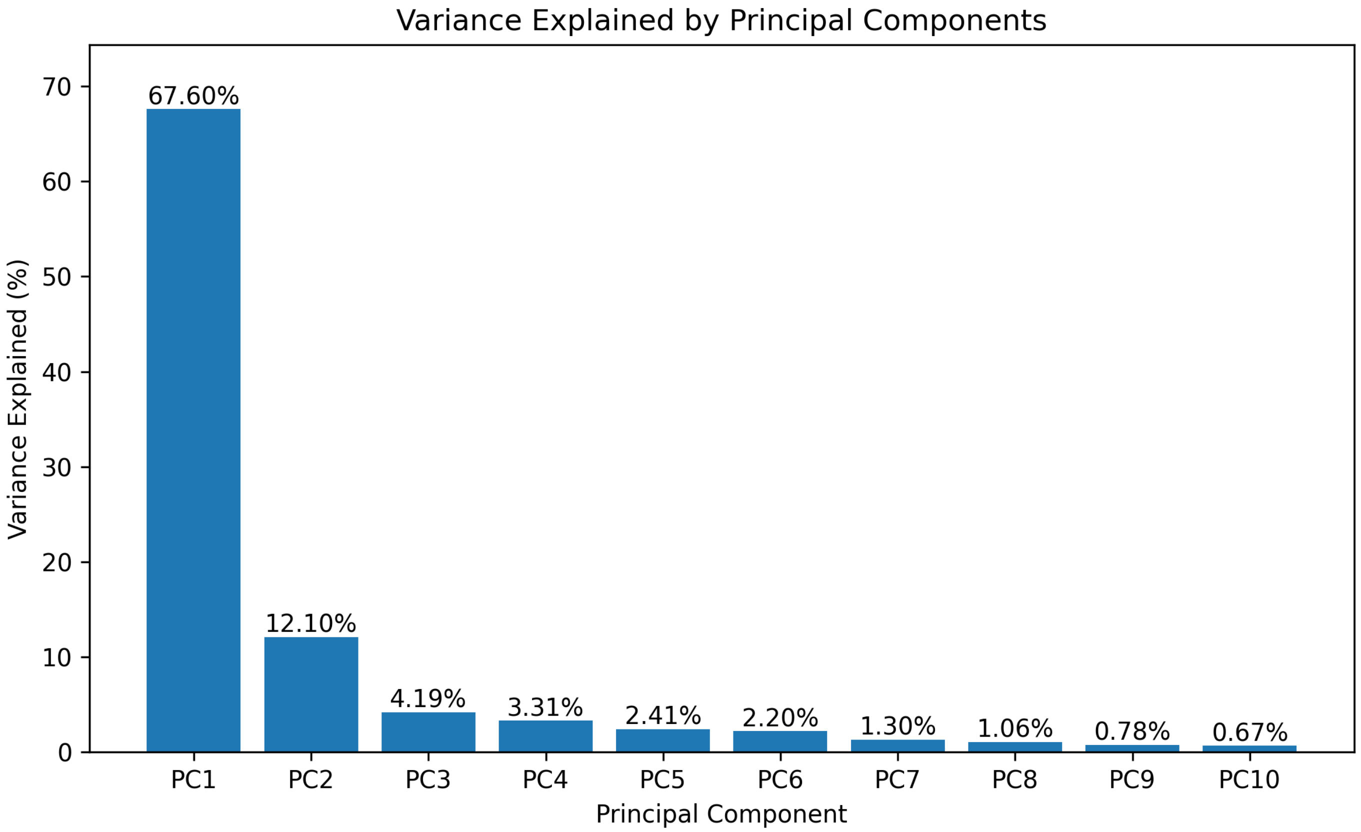


**Figure 4** Training results for shell detection.

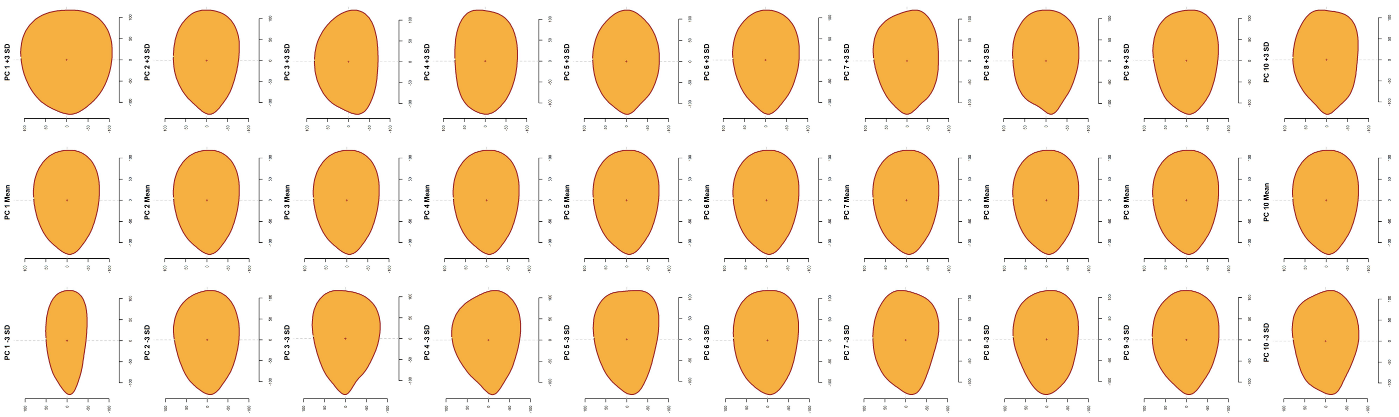

**Figure 5** Training results for shell detection.

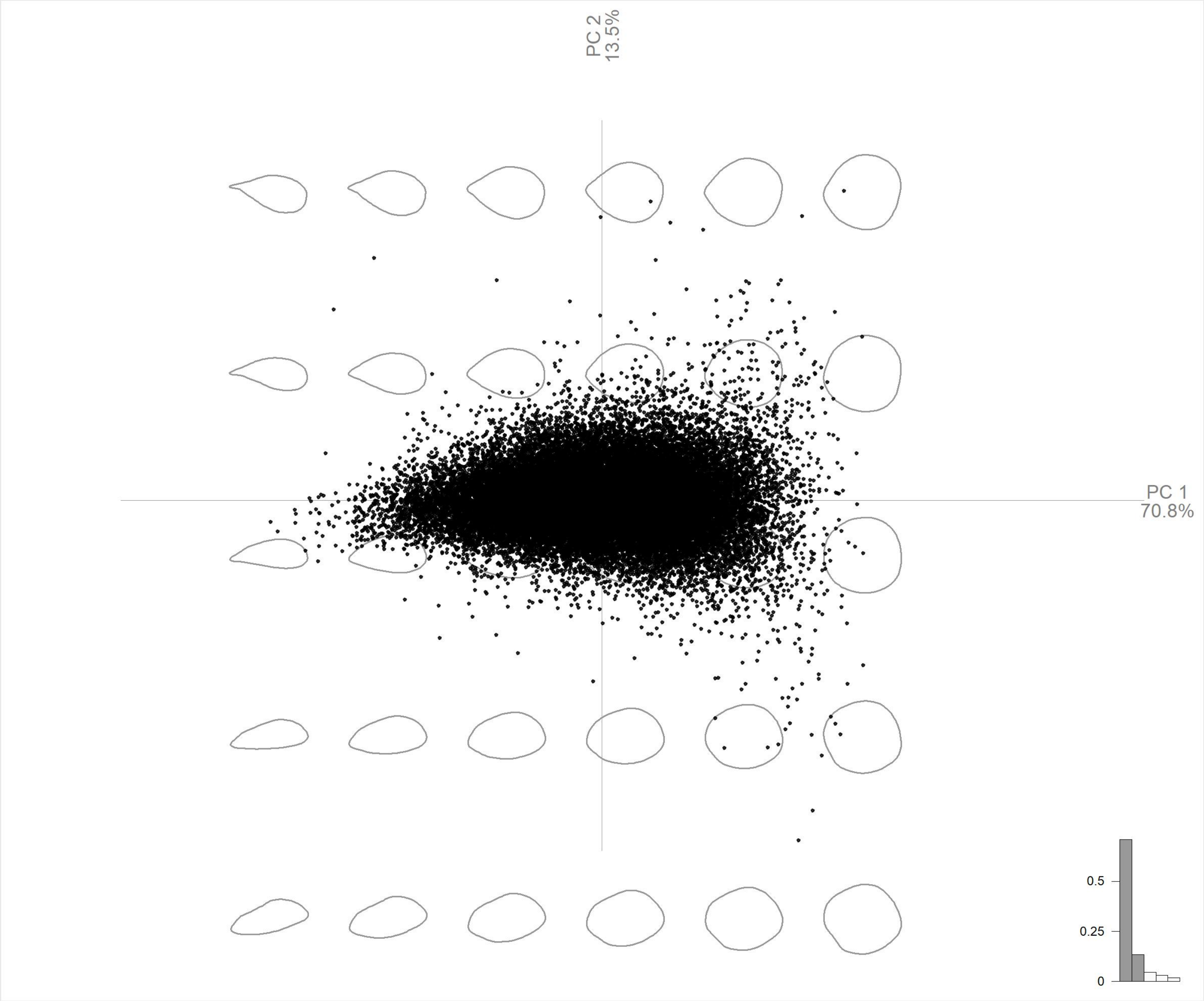


**Figure 6** Training results for shell detection.

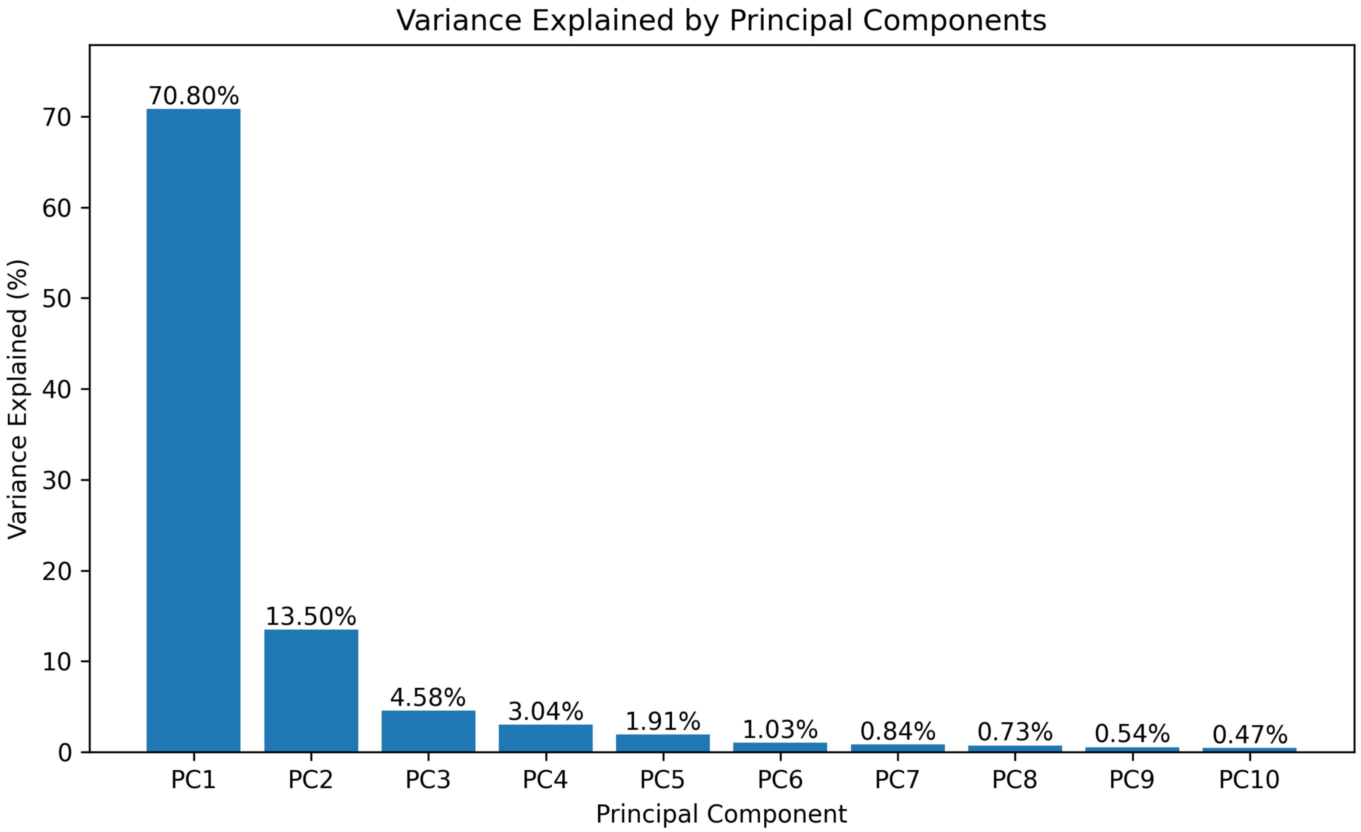


**Figure 7** Training results for shell detection.

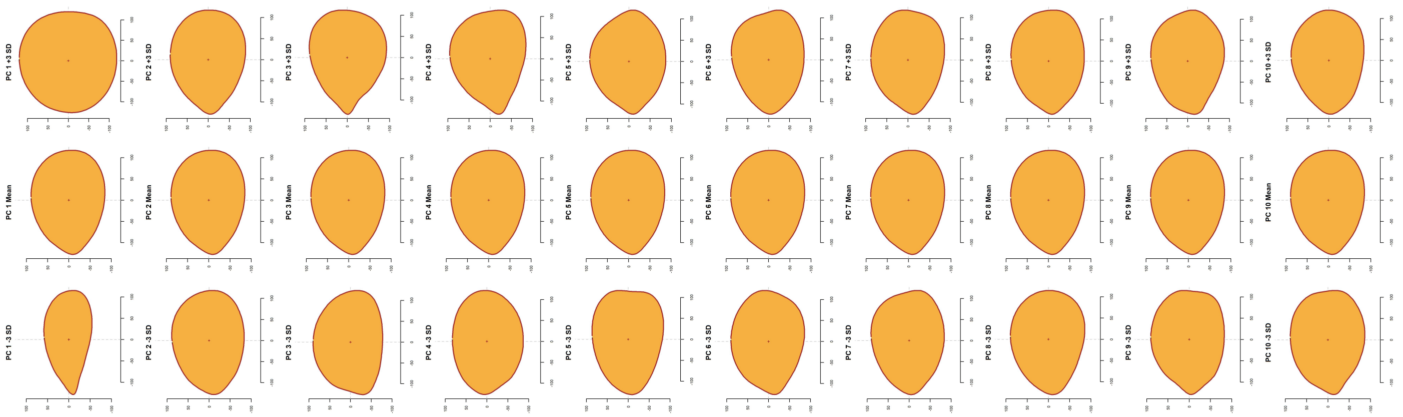

**Figure 8** Training results for shell detection.

#### Appendix 3

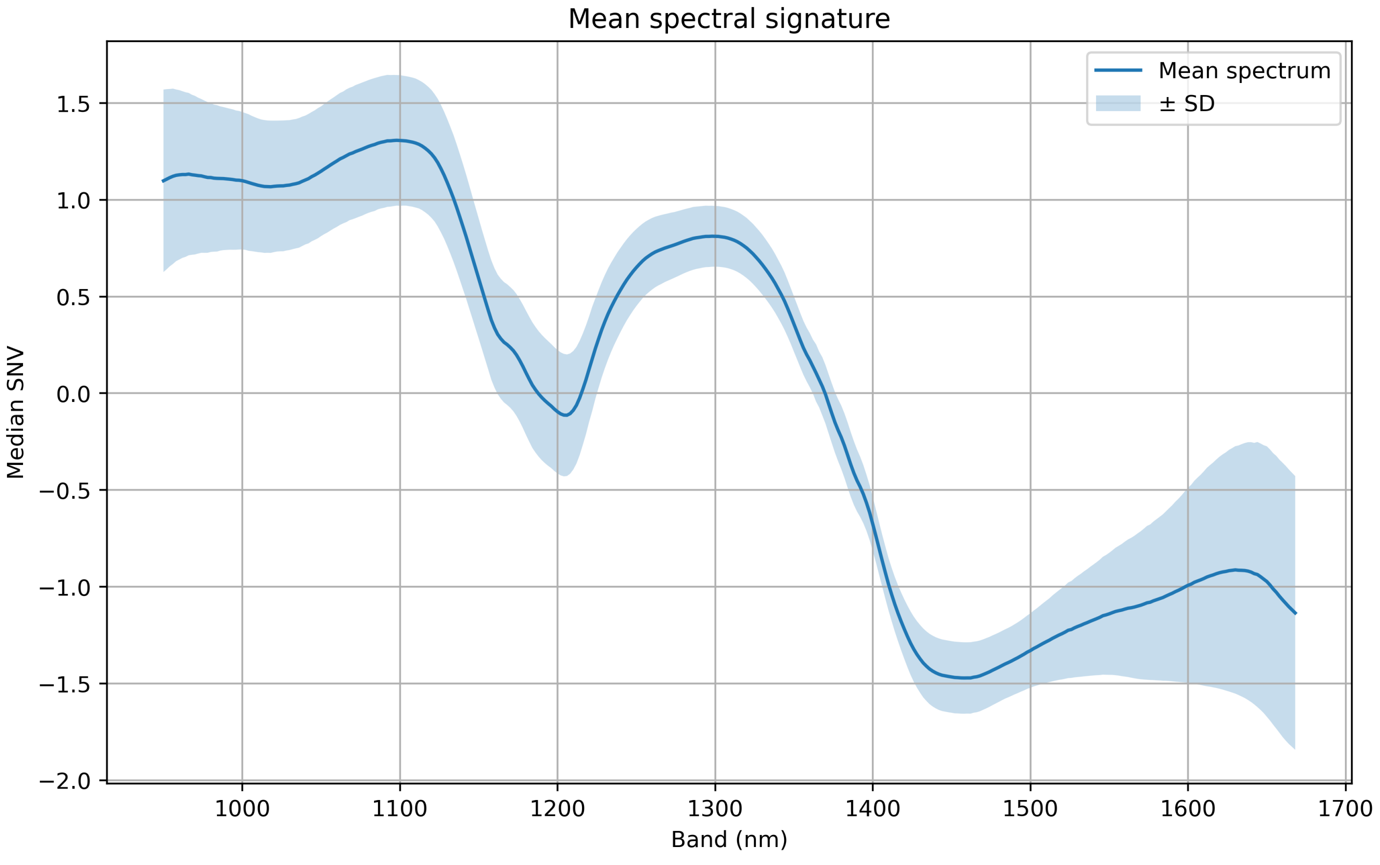


**Figure 9** Training results for shell detection.

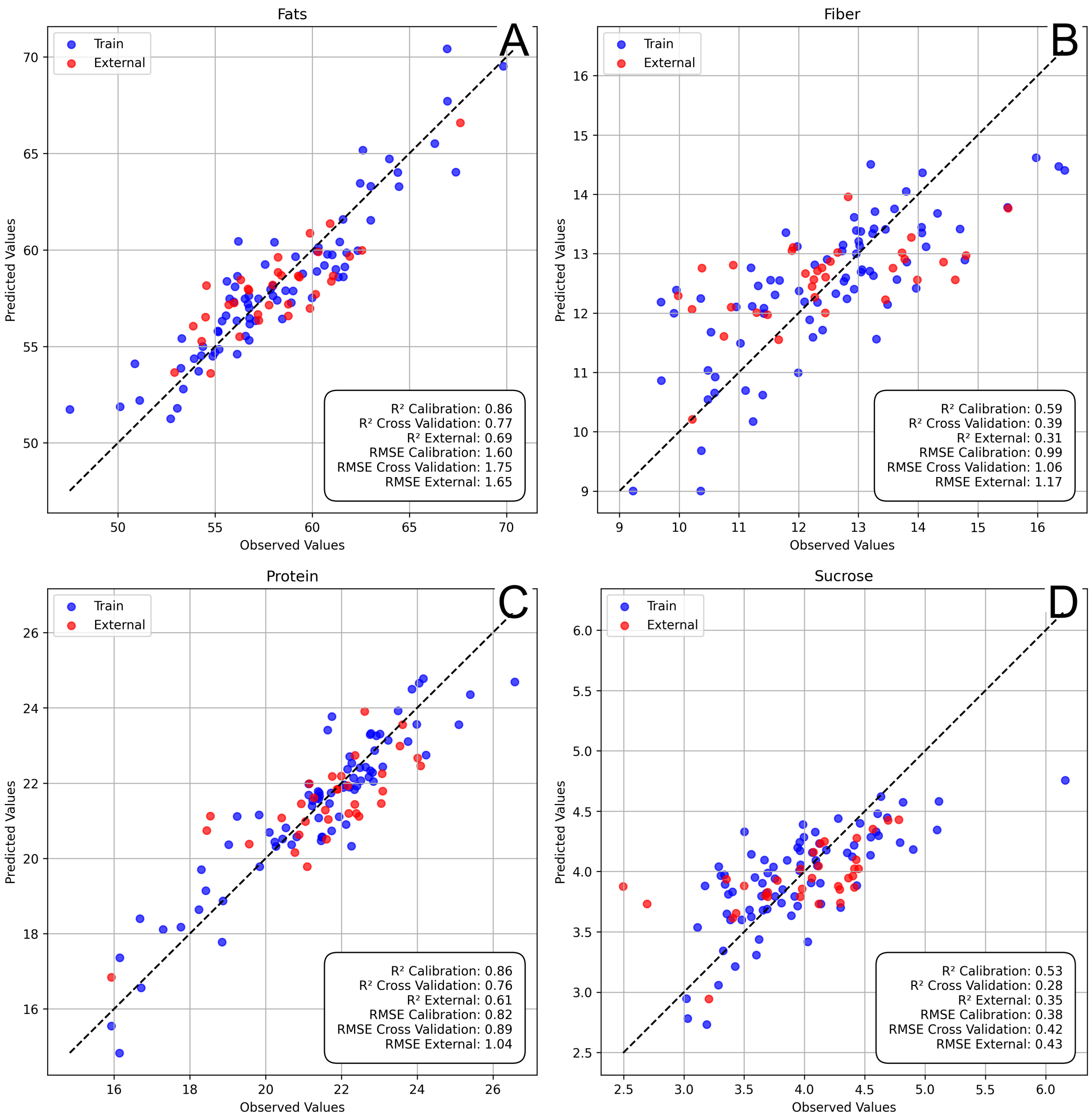


**Figure 10** Training results for shell detection.

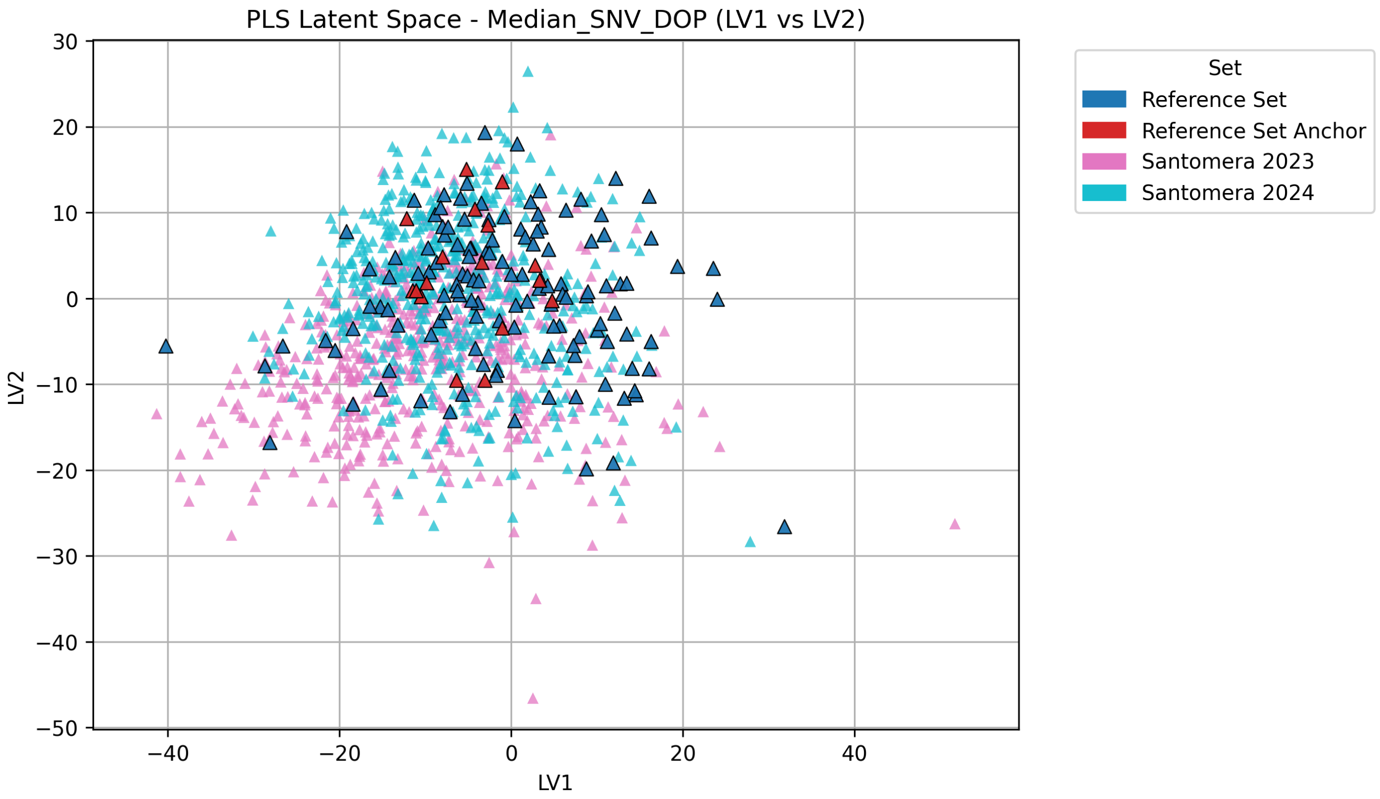
PLS Latent Space - Median_SNV_DOP (LV1 vs LV2)
LV1
LV2
Set
Reference Set
Reference Set Anchor
Santomera 2023
Santomera 2024

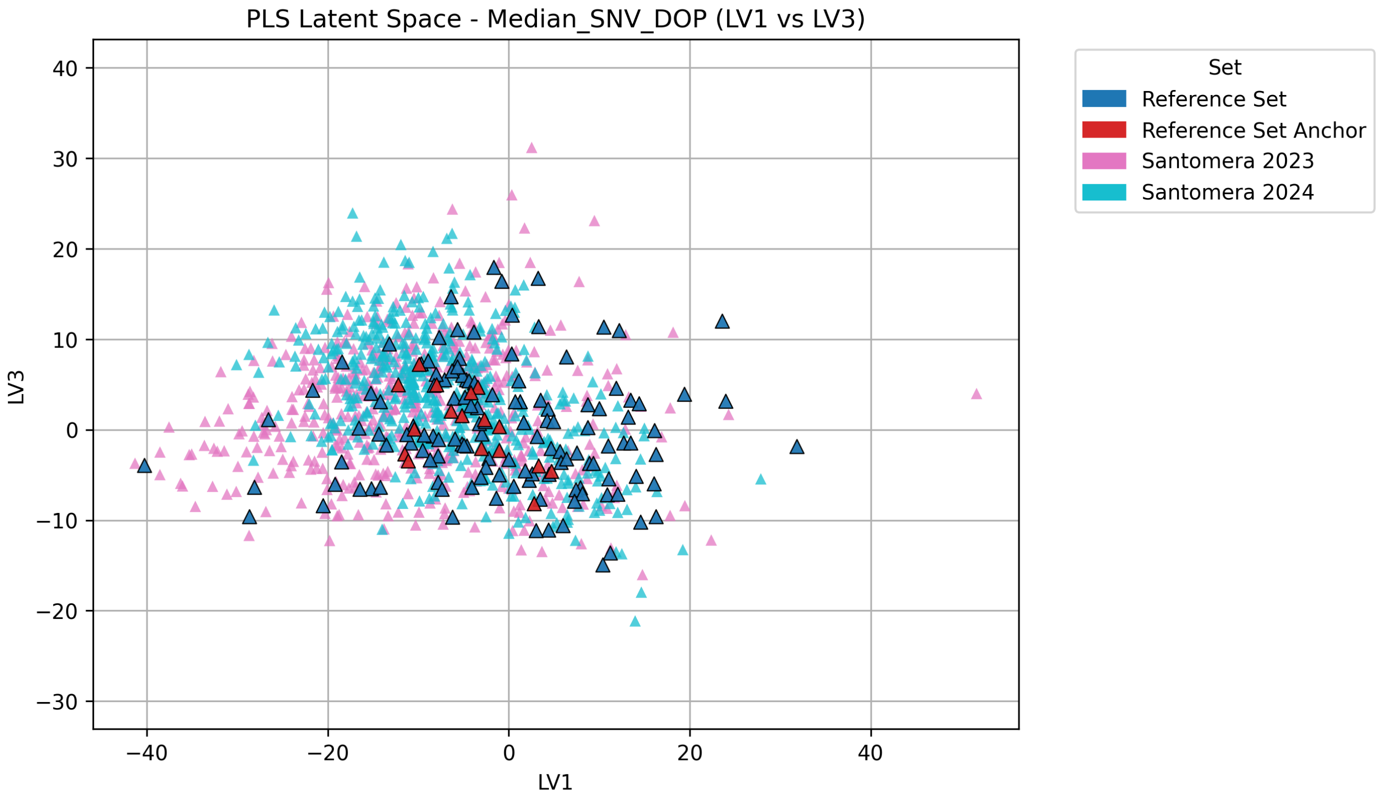
PLS Latent Space - Median_SNV_DOP (LV1 vs LV3)
LV1
LV3
Set
Reference Set
Reference Set Anchor
Santomera 2023
Santomera 2024

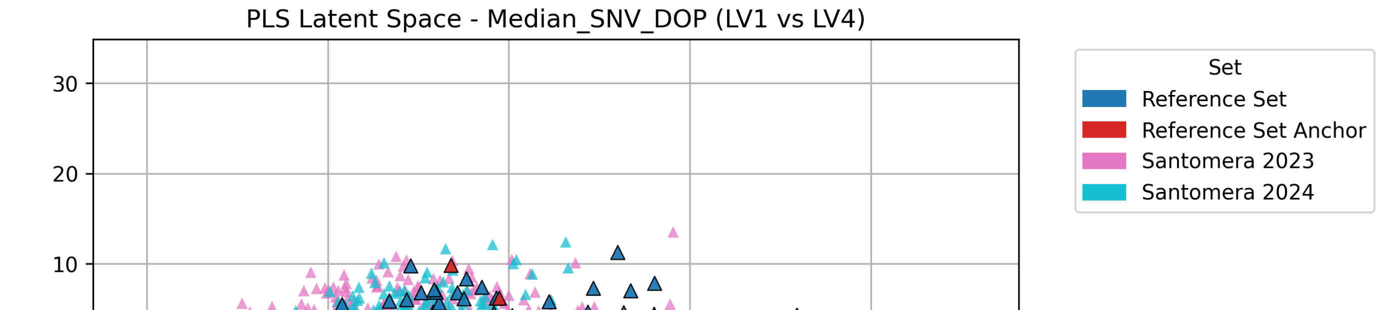
PLS Latent Space - Median_SNV_DOP (LV1 vs LV4)
Set
Reference Set
Reference Set Anchor
Santomera 2023
Santomera 2024

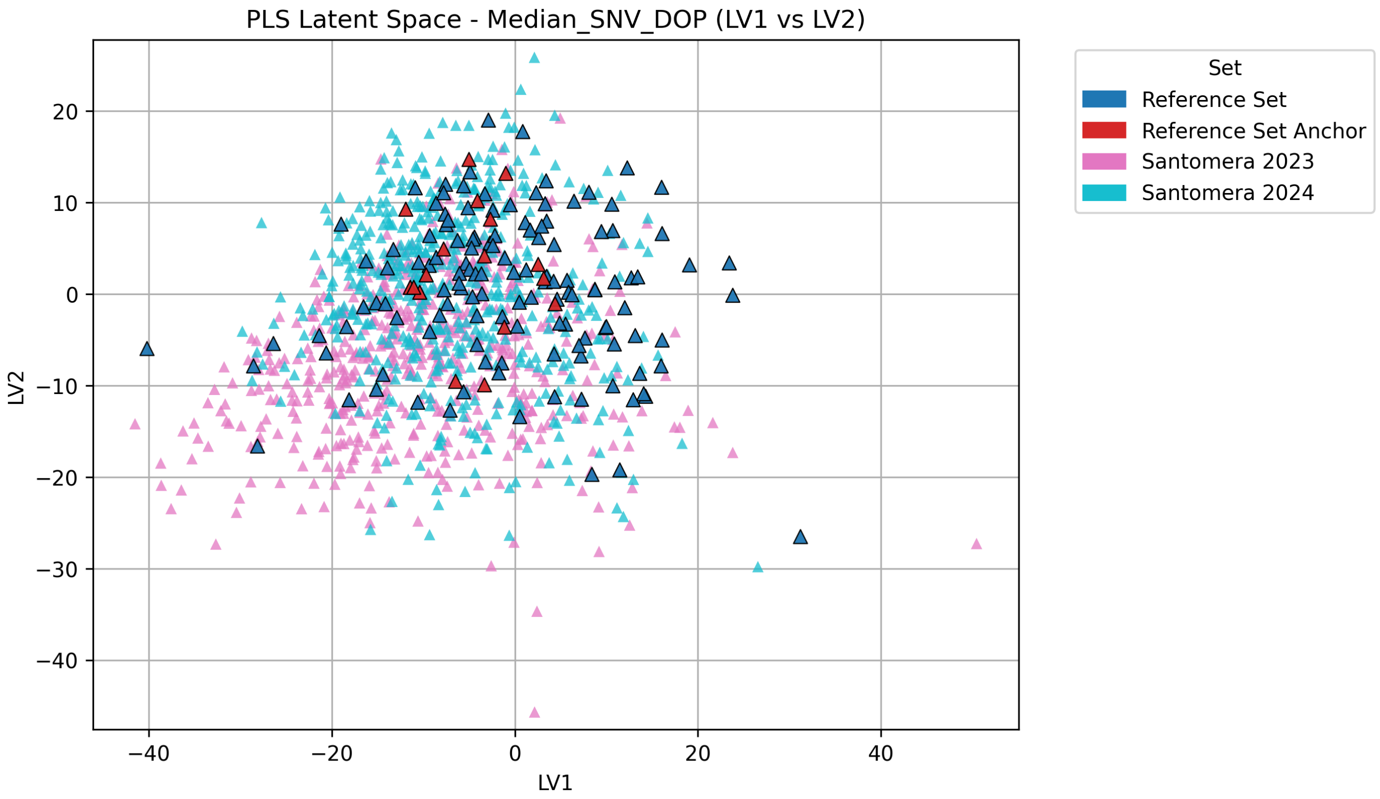
PLS Latent Space - Median_SNV_DOP (LV1 vs LV2)
LV2
LV1
Set
Reference Set
Reference Set Anchor
Santomera 2023
Santomera 2024

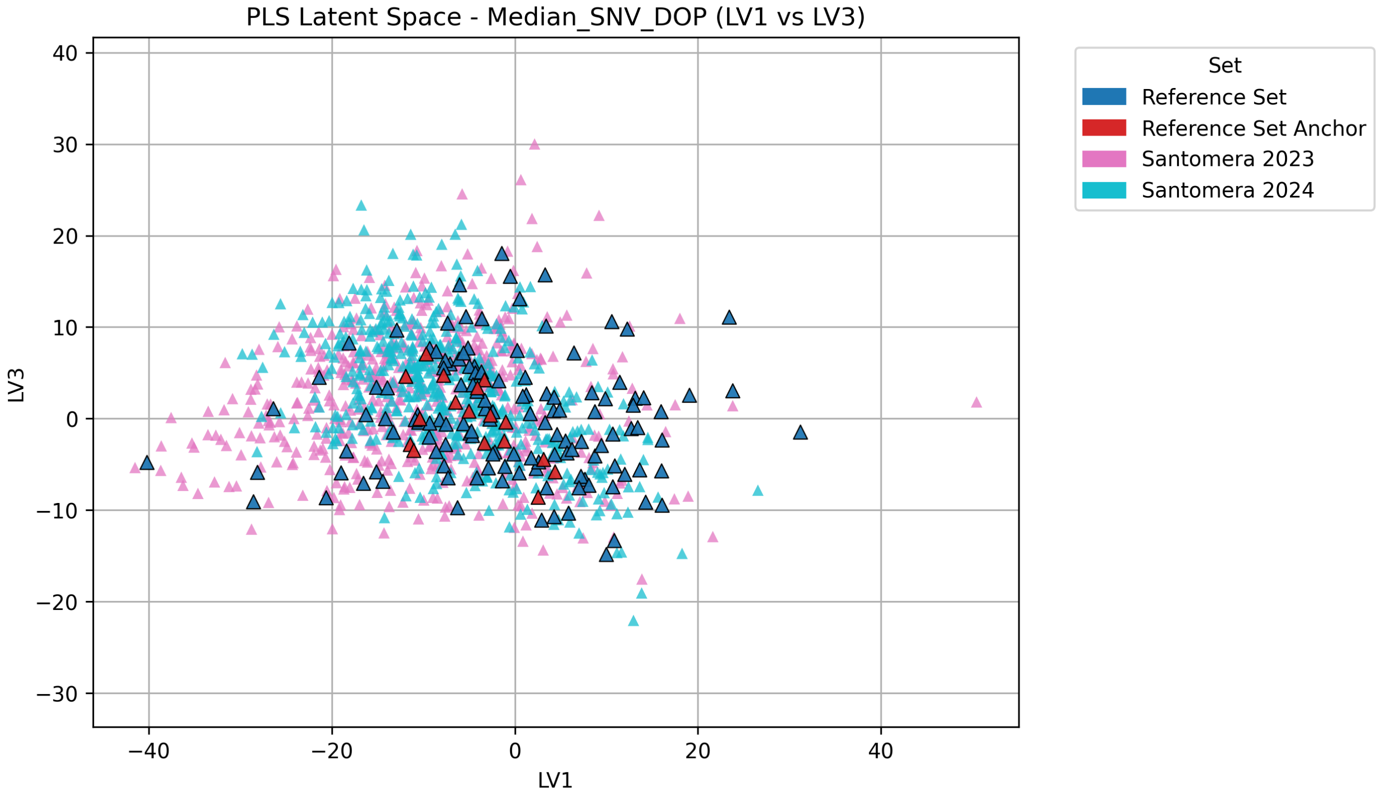
PLS Latent Space - Median_SNV_DOP (LV1 vs LV3)
LV3
LV1
Set
Reference Set
Reference Set Anchor
Santomera 2023
Santomera 2024

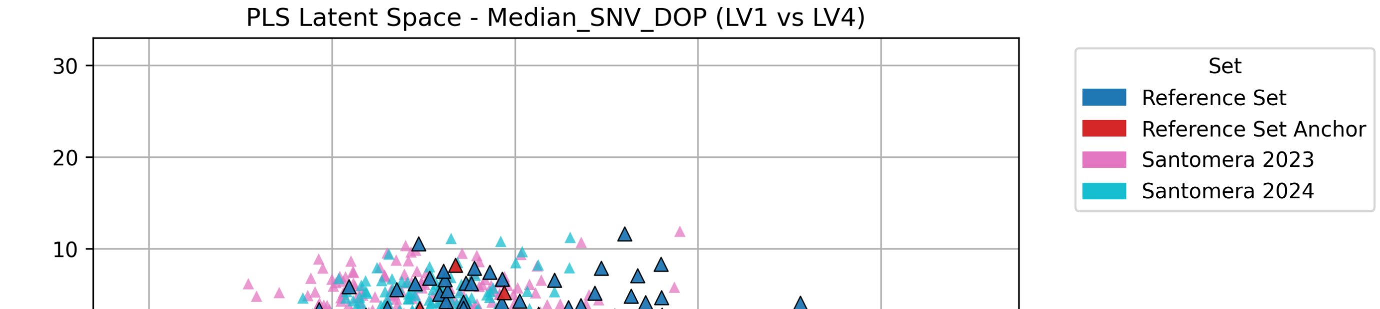
PLS Latent Space - Median_SNV_DOP (LV1 vs LV4)
Set
Reference Set
Reference Set Anchor
Santomera 2023
Santomera 2024

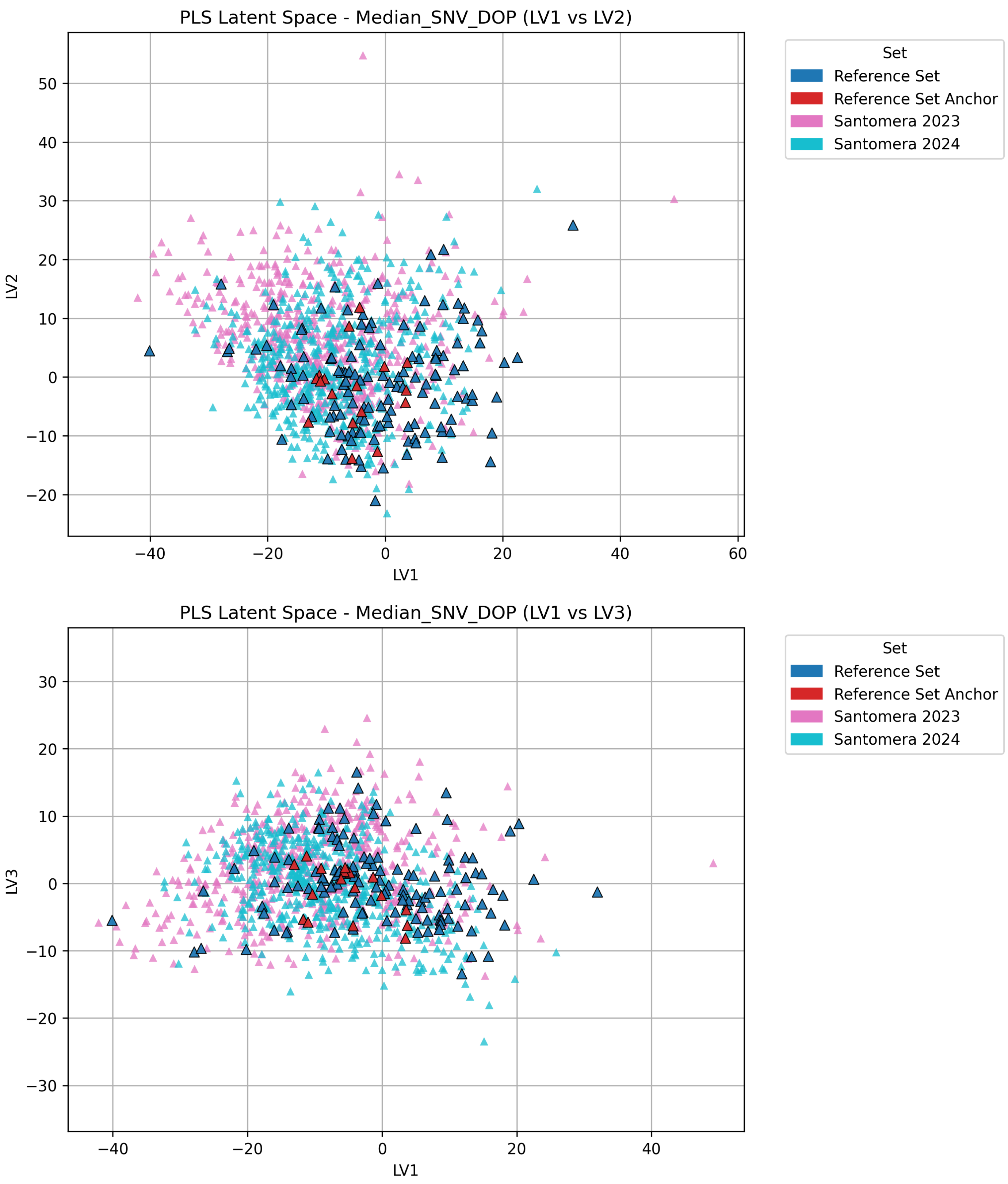


**Figure 13** Training results for shell detection.

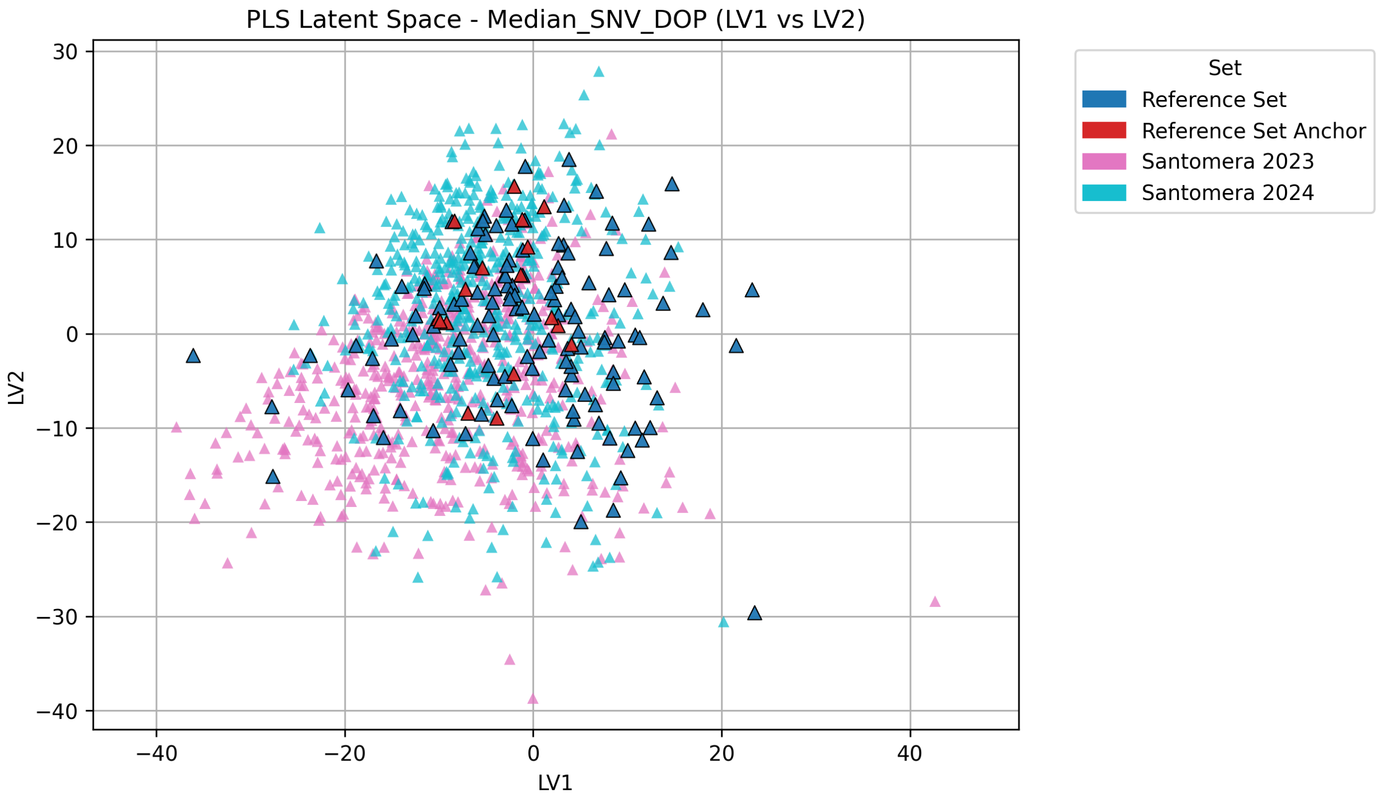
PLS Latent Space - Median_SNV_DOP (LV1 vs LV2)
LV1
LV2
Set
Reference Set
Reference Set Anchor
Santomera 2023
Santomera 2024

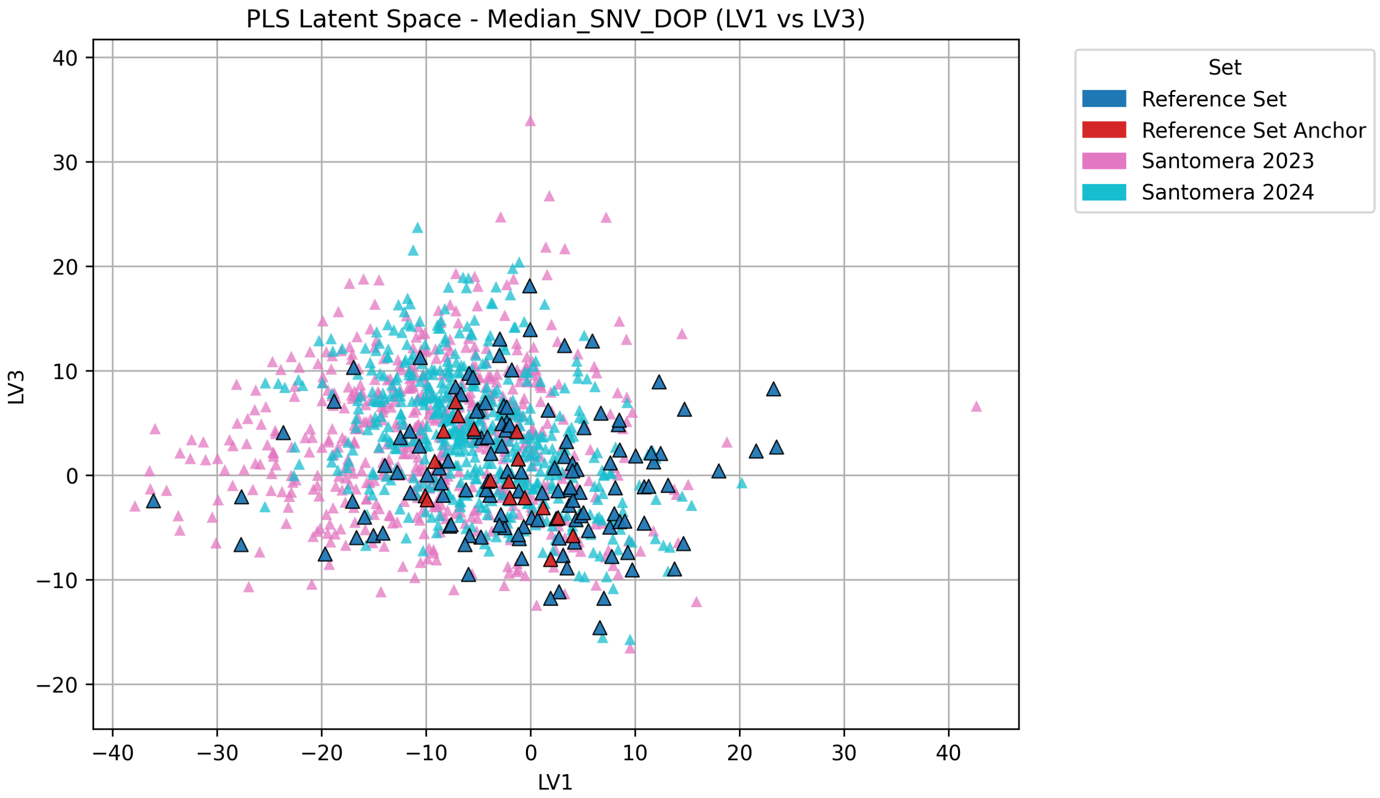
PLS Latent Space - Median_SNV_DOP (LV1 vs LV3)
LV1
LV3
Set
Reference Set
Reference Set Anchor
Santomera 2023
Santomera 2024

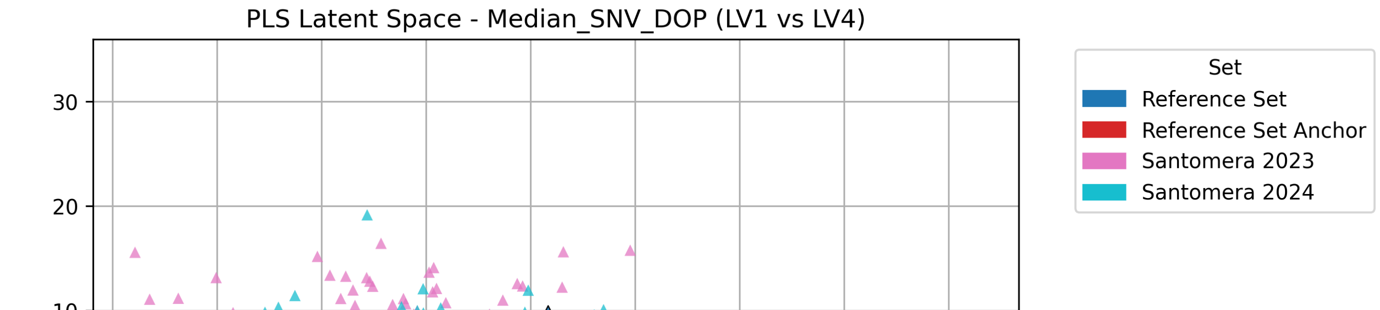
PLS Latent Space - Median_SNV_DOP (LV1 vs LV4)
Set
Reference Set
Reference Set Anchor
Santomera 2023
Santomera 2024

### Appendix 4

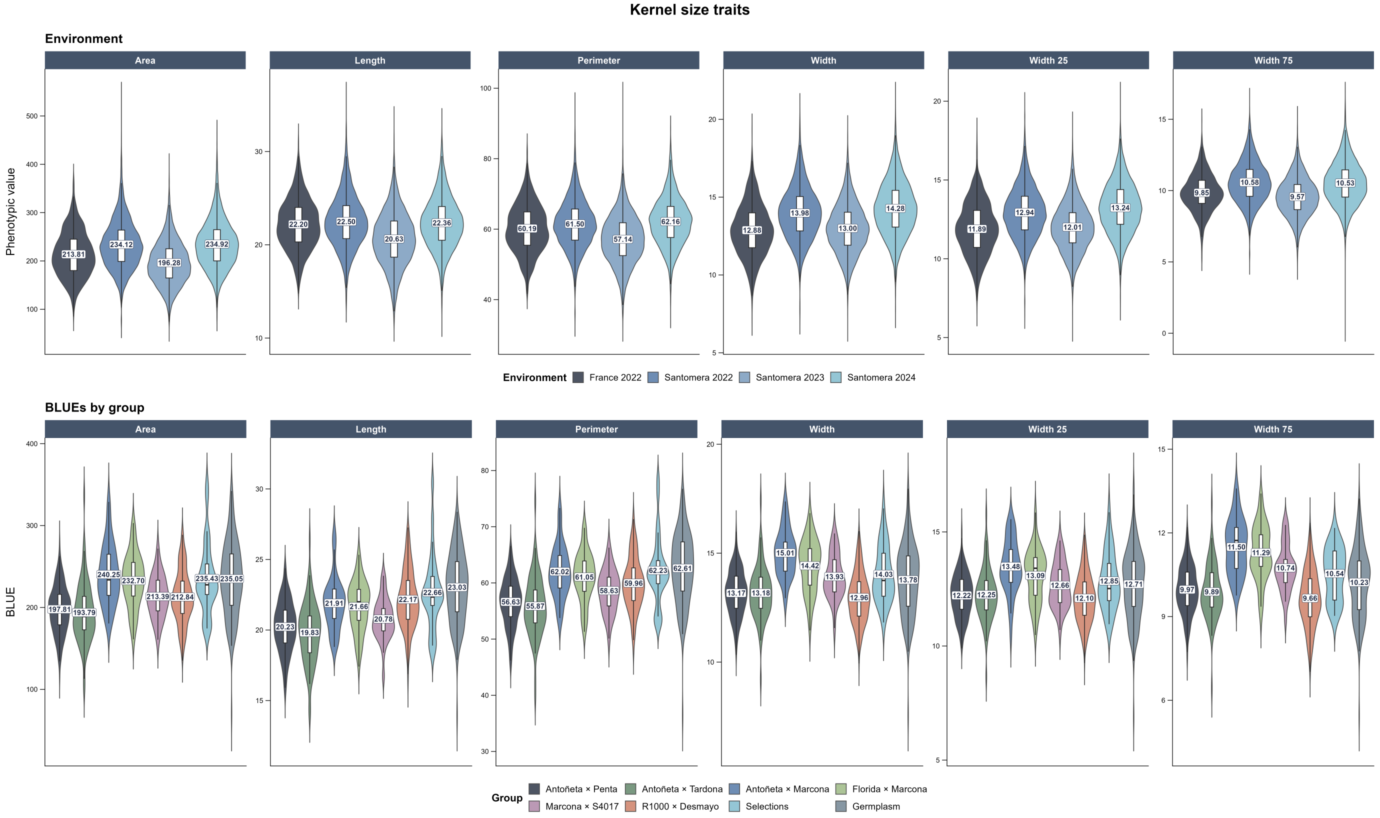


**Figure 15** Distribution of BLUEs by group and environment for kernel size traits.

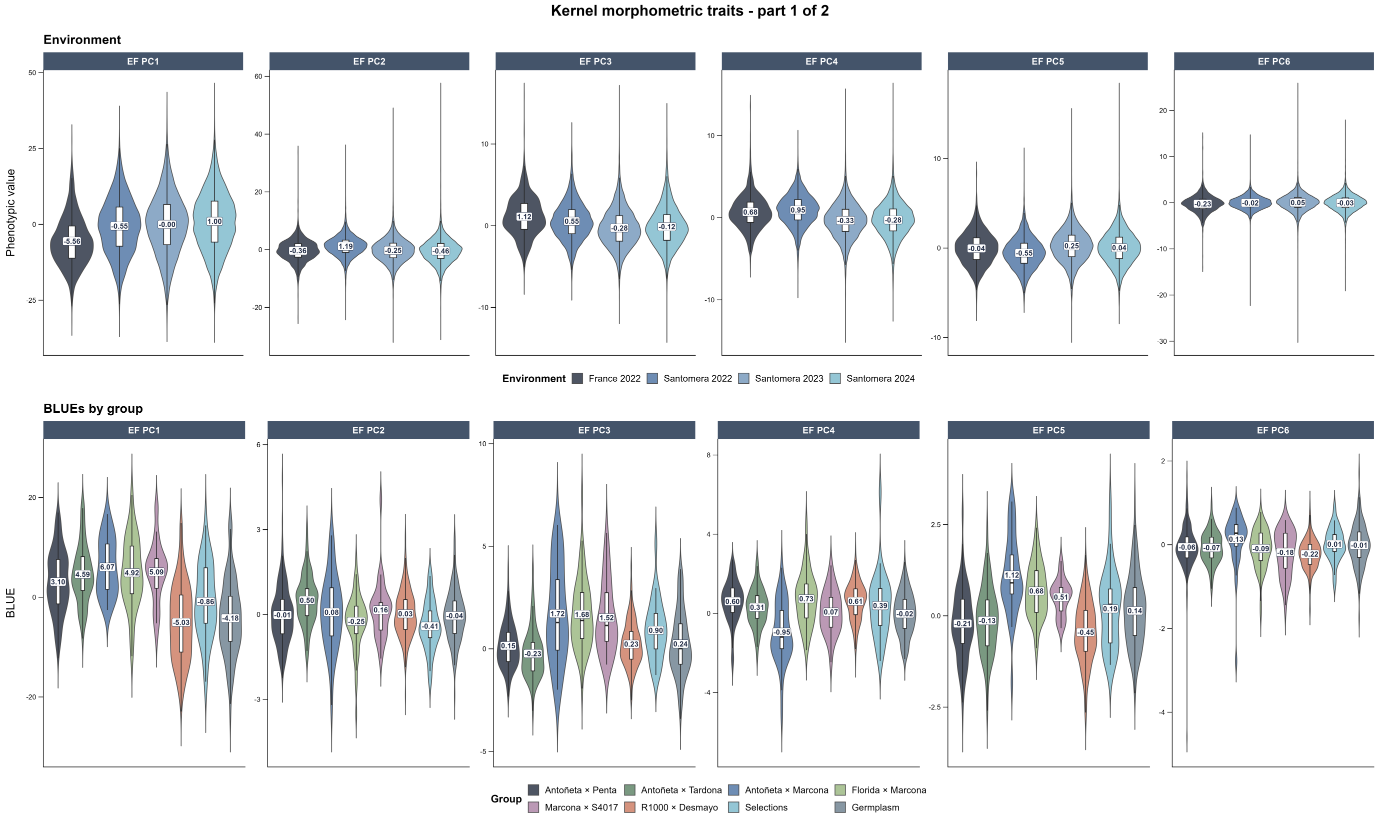


**Figure 16** Distribution of BLUEs by group and environment for kernel morphometric traits, part 1.

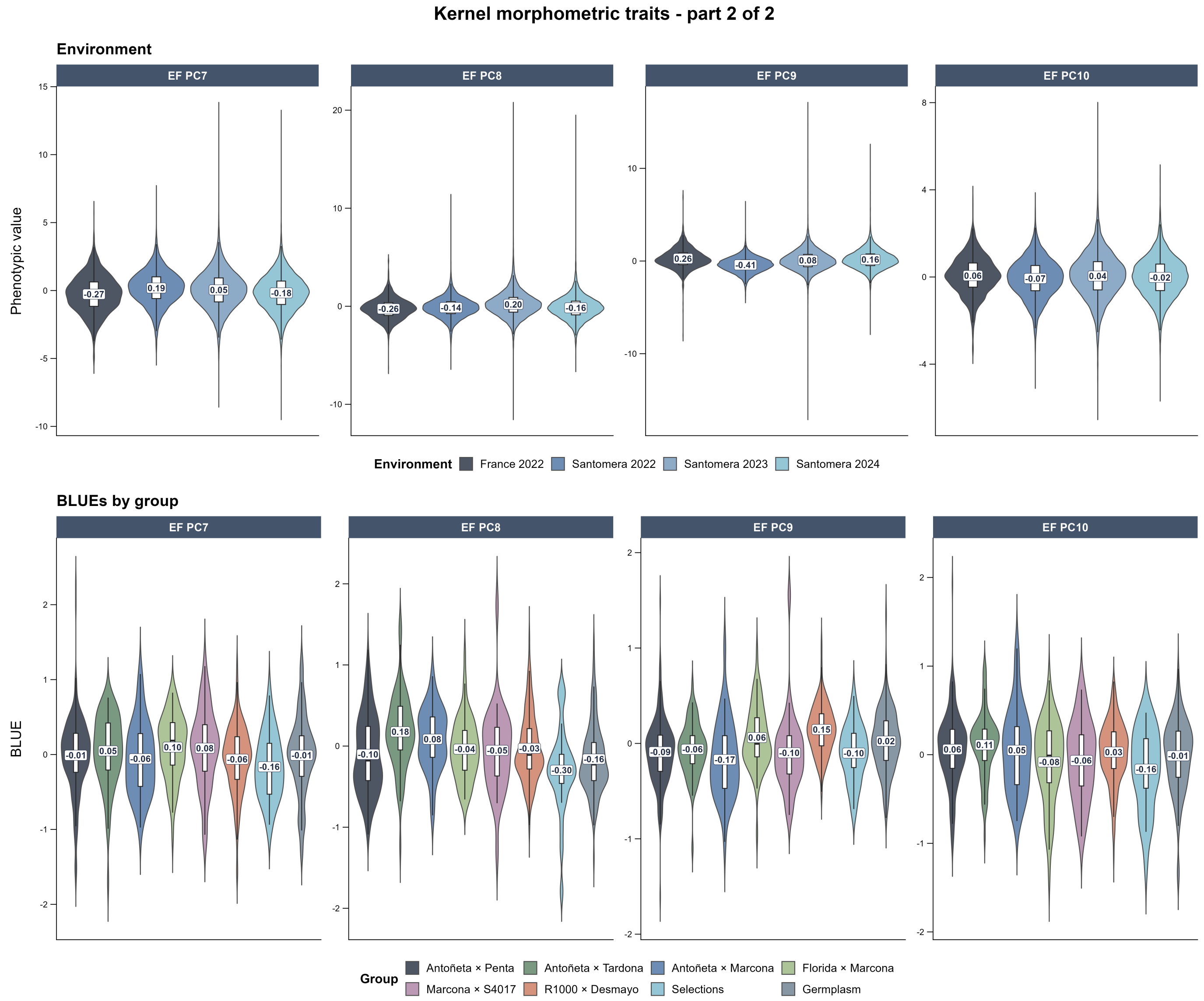


**Figure 17** Distribution of BLUEs by group and environment for kernel morphometric traits, part 2.

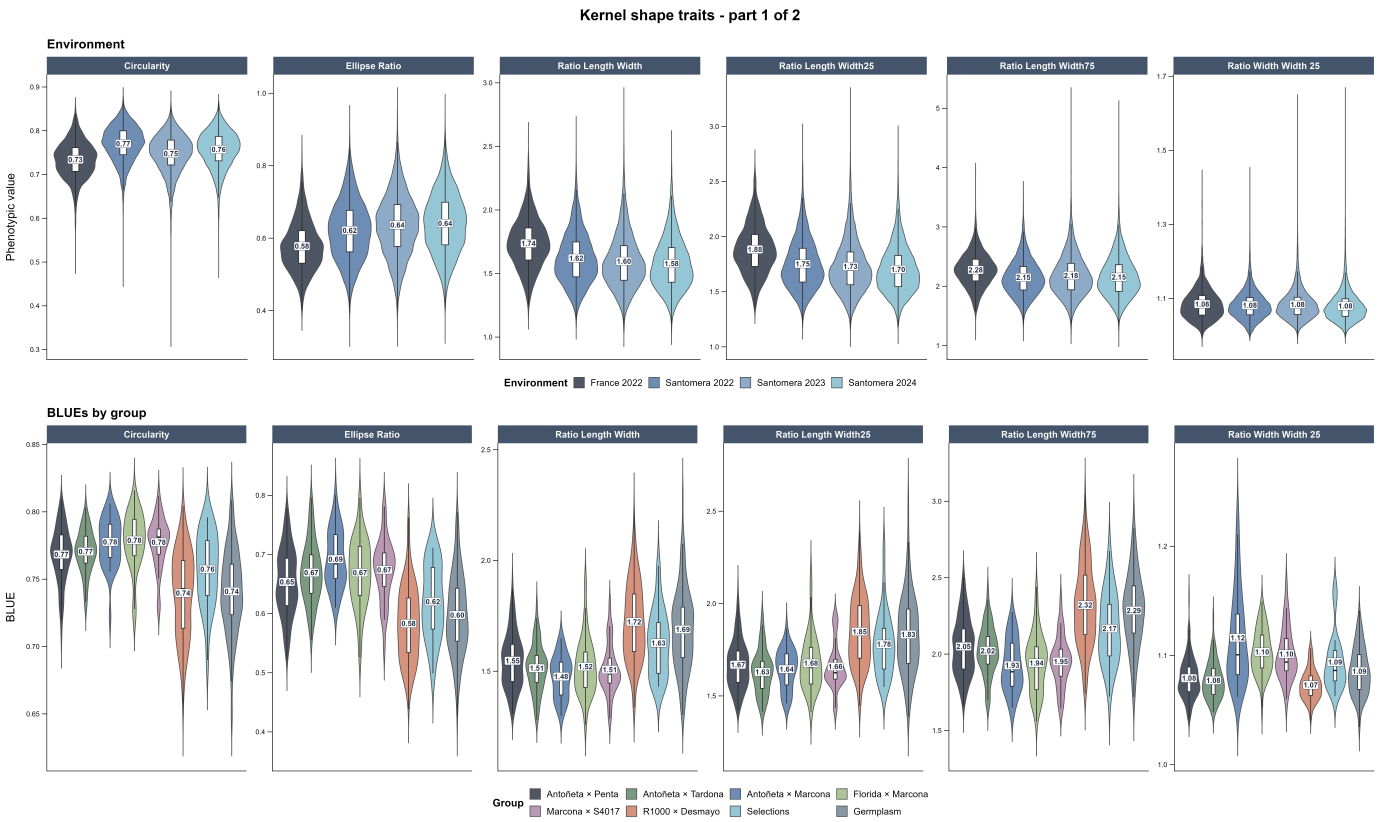


**Figure 18** Distribution of BLUEs by group and environment for kernel shape traits, part 1.

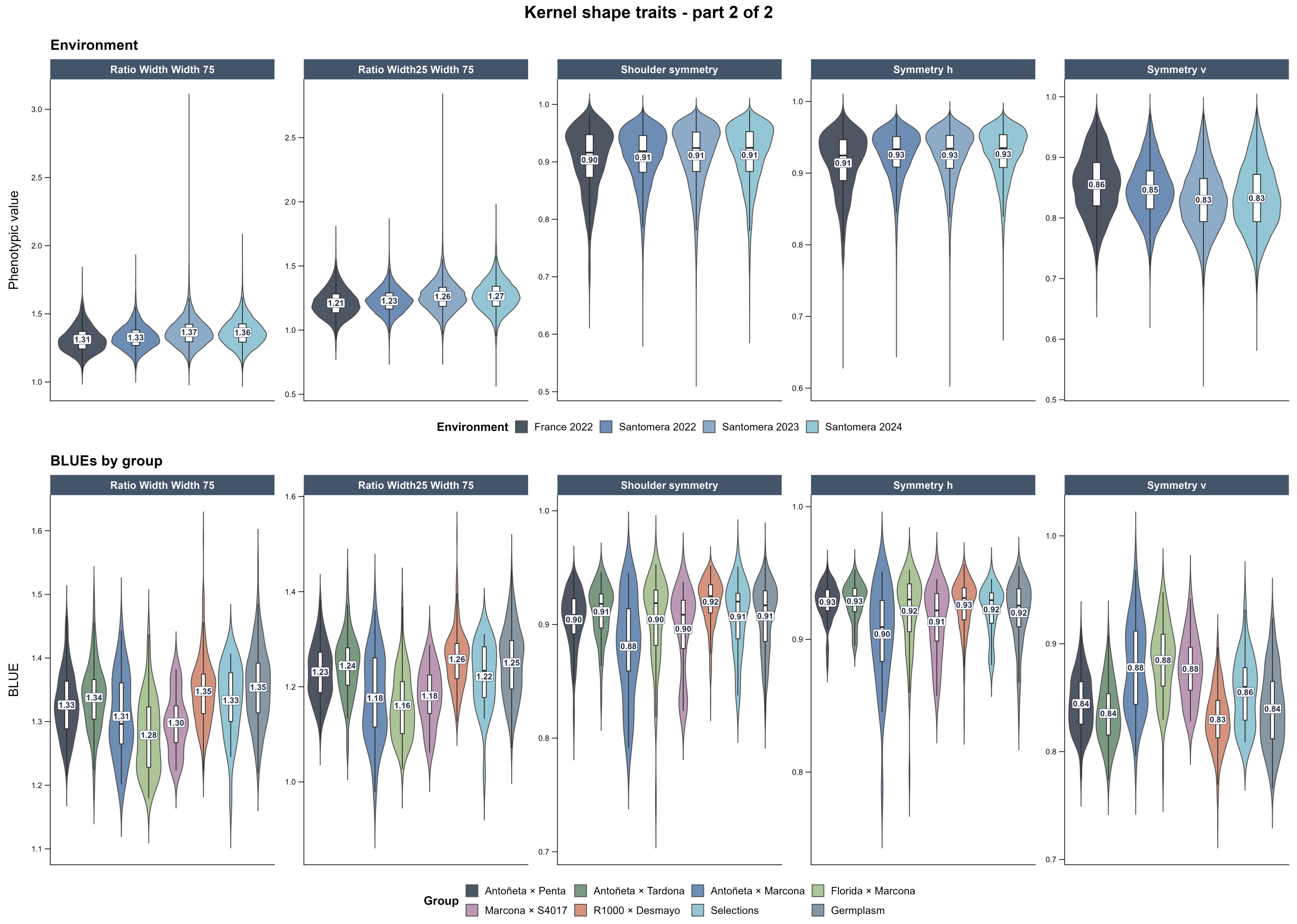


**Figure 19** Distribution of BLUEs by group and environment for kernel shape traits, part 2.

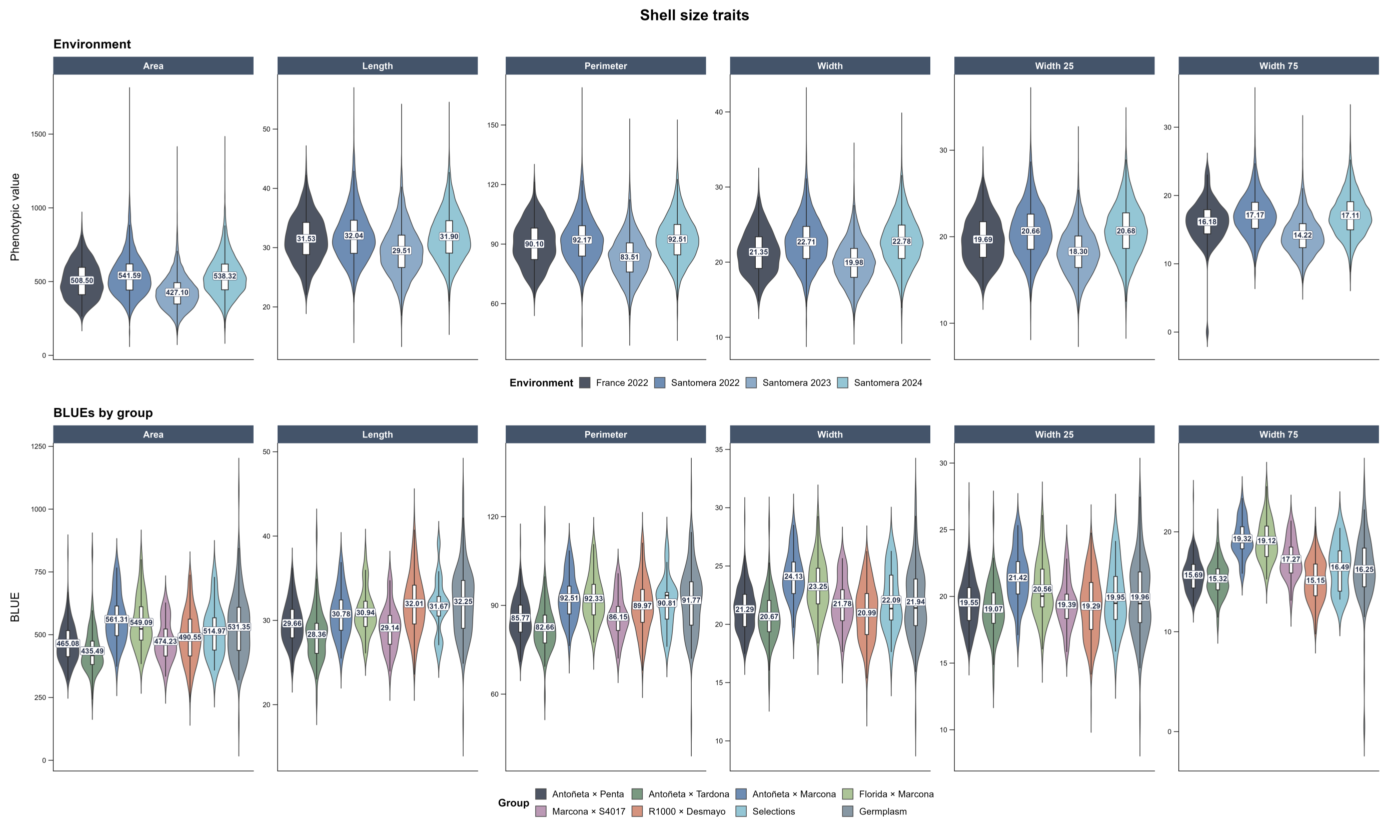


**Figure 20** Distribution of BLUEs by group and environment for shell size traits.

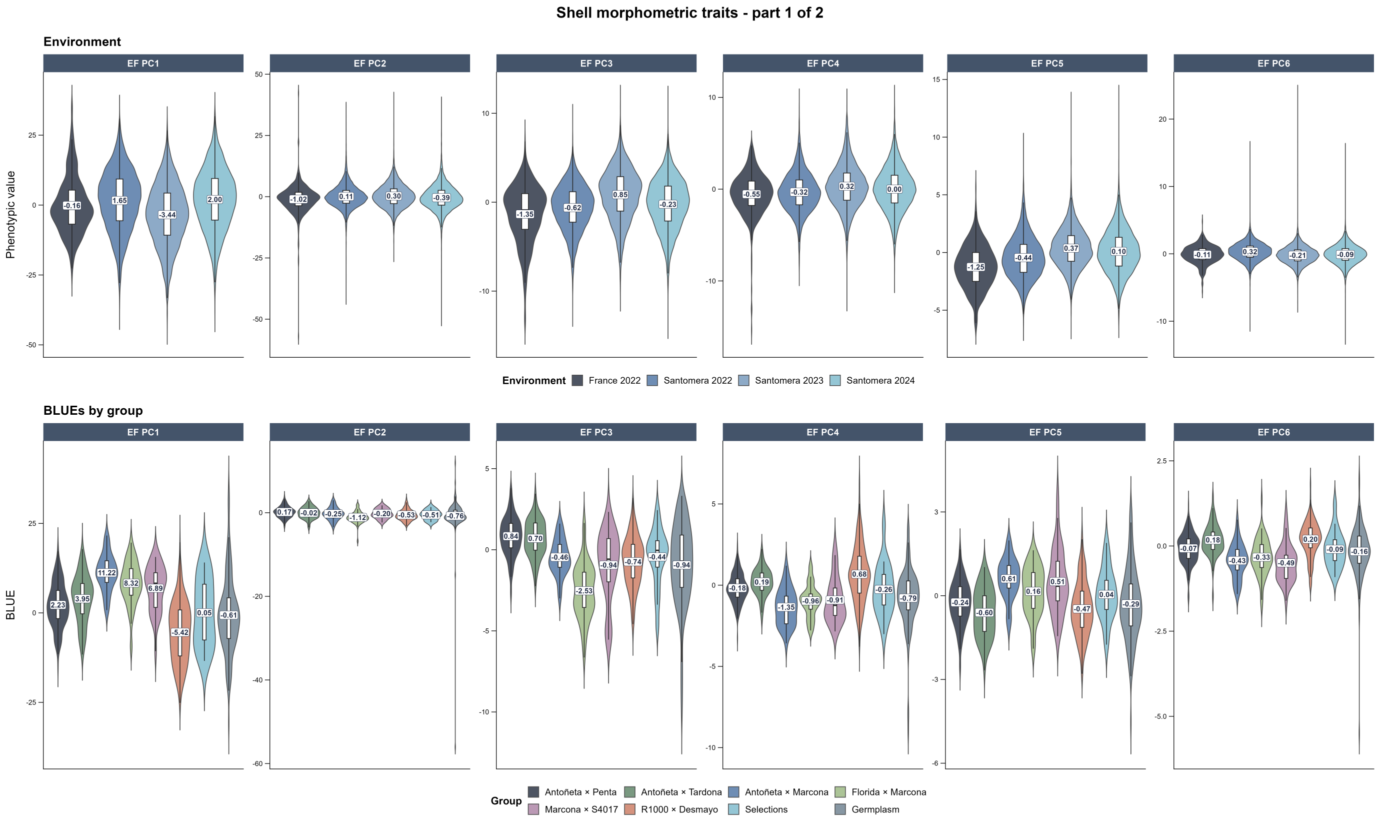


**Figure 21** Distribution of BLUEs by group and environment for shell morphometric traits, part 1.

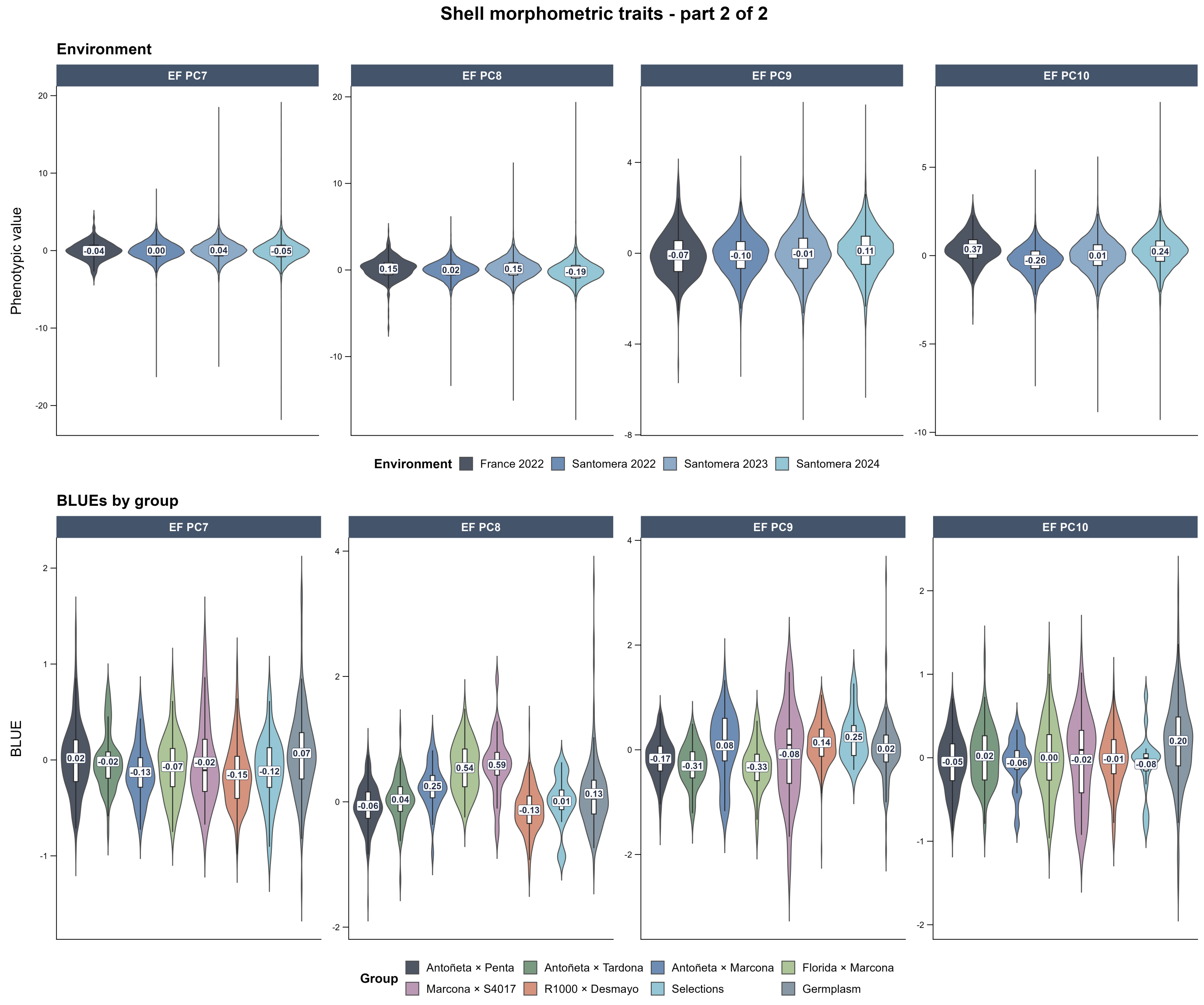


**Figure 22** Distribution of BLUEs by group and environment for shell morphometric traits, part 2.

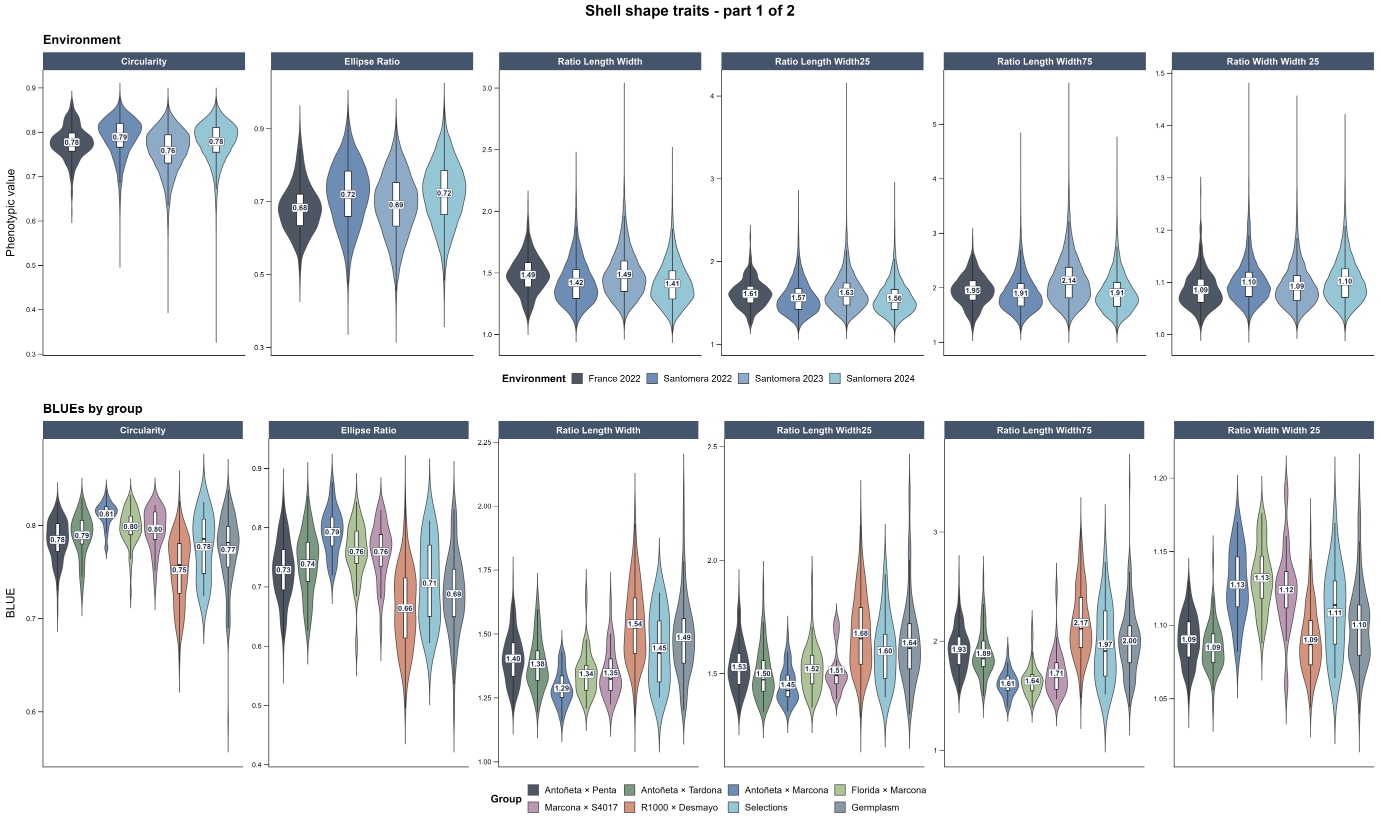


**Figure 23** Distribution of BLUEs by group and environment for shell shape traits, part 1.

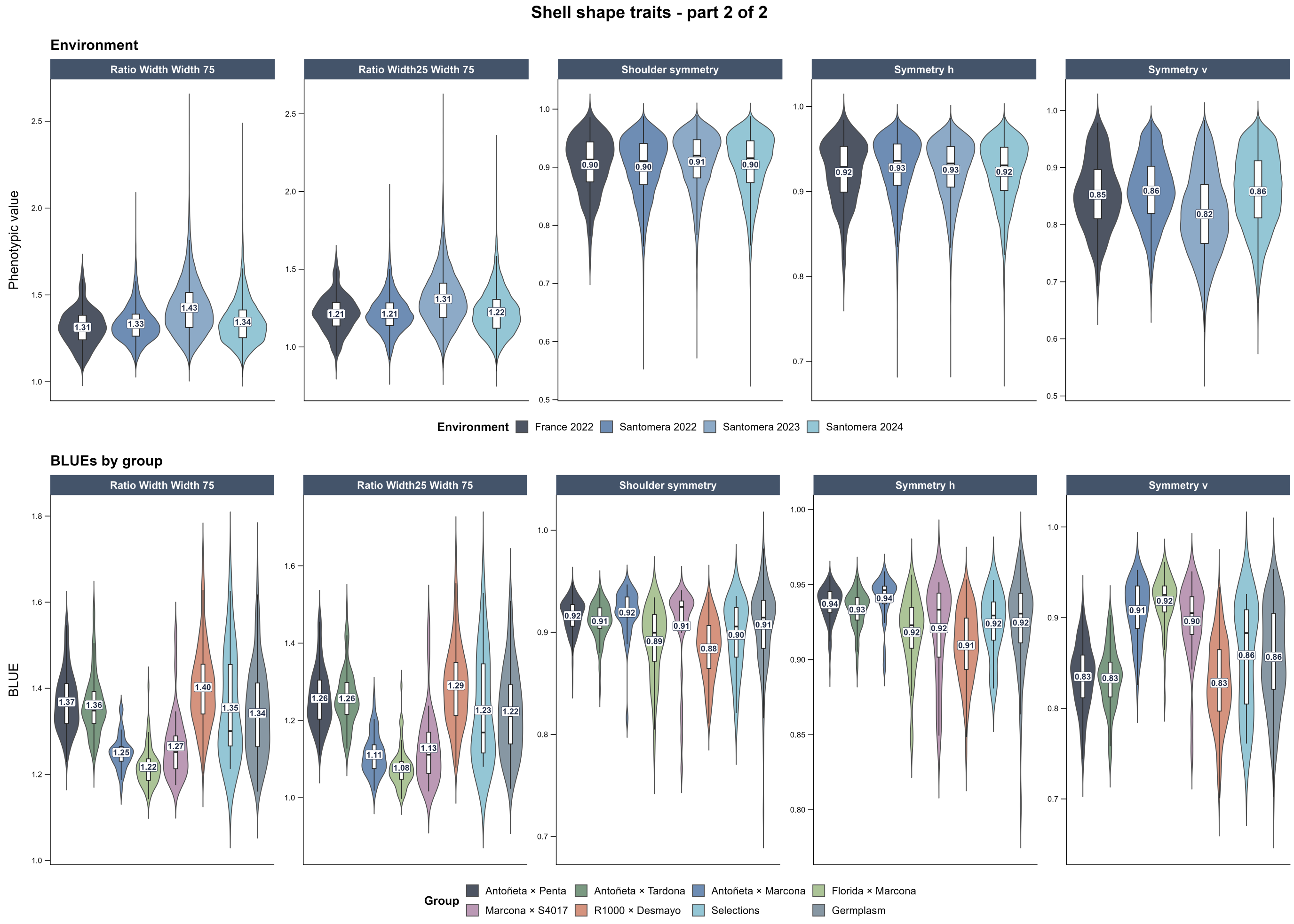


**Figure 24** Distribution of BLUEs by group and environment for shell shape traits, part 2.

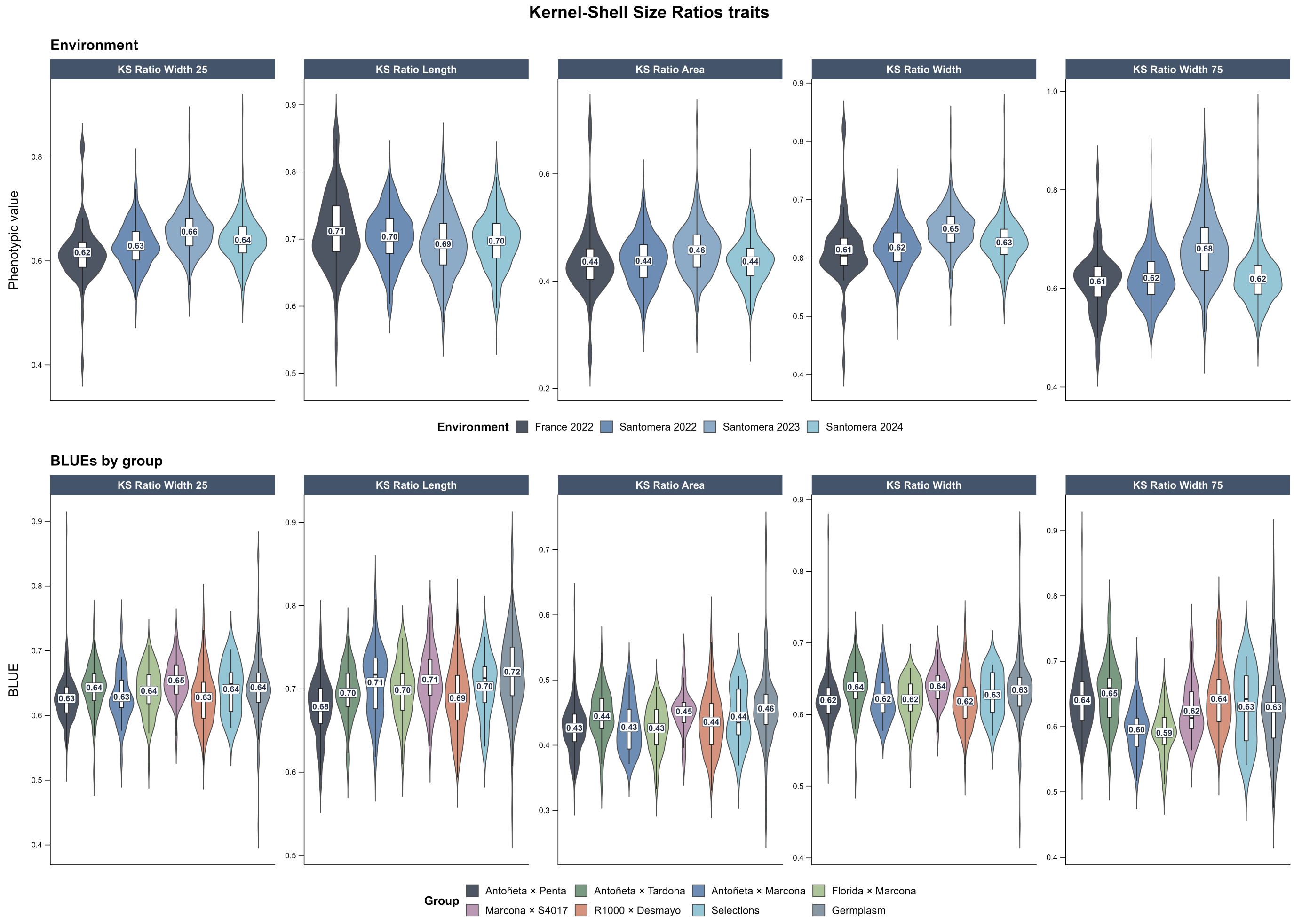


**Figure 25** Distribution of BLUEs by group and environment for kernel-to-shell size ratios.

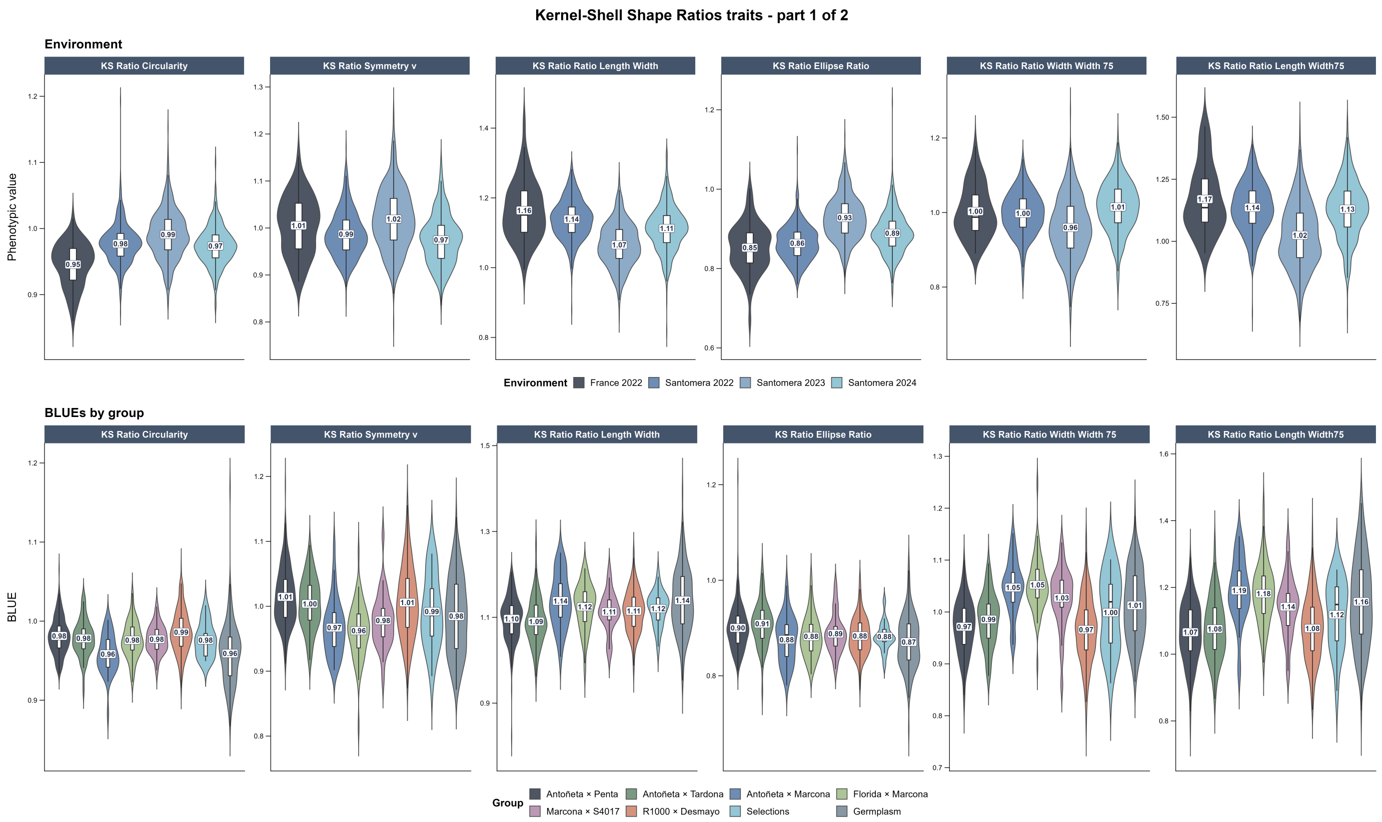


**Figure 26** Distribution of BLUEs by group and environment for kernel-to-shell shape ratios, part 1.

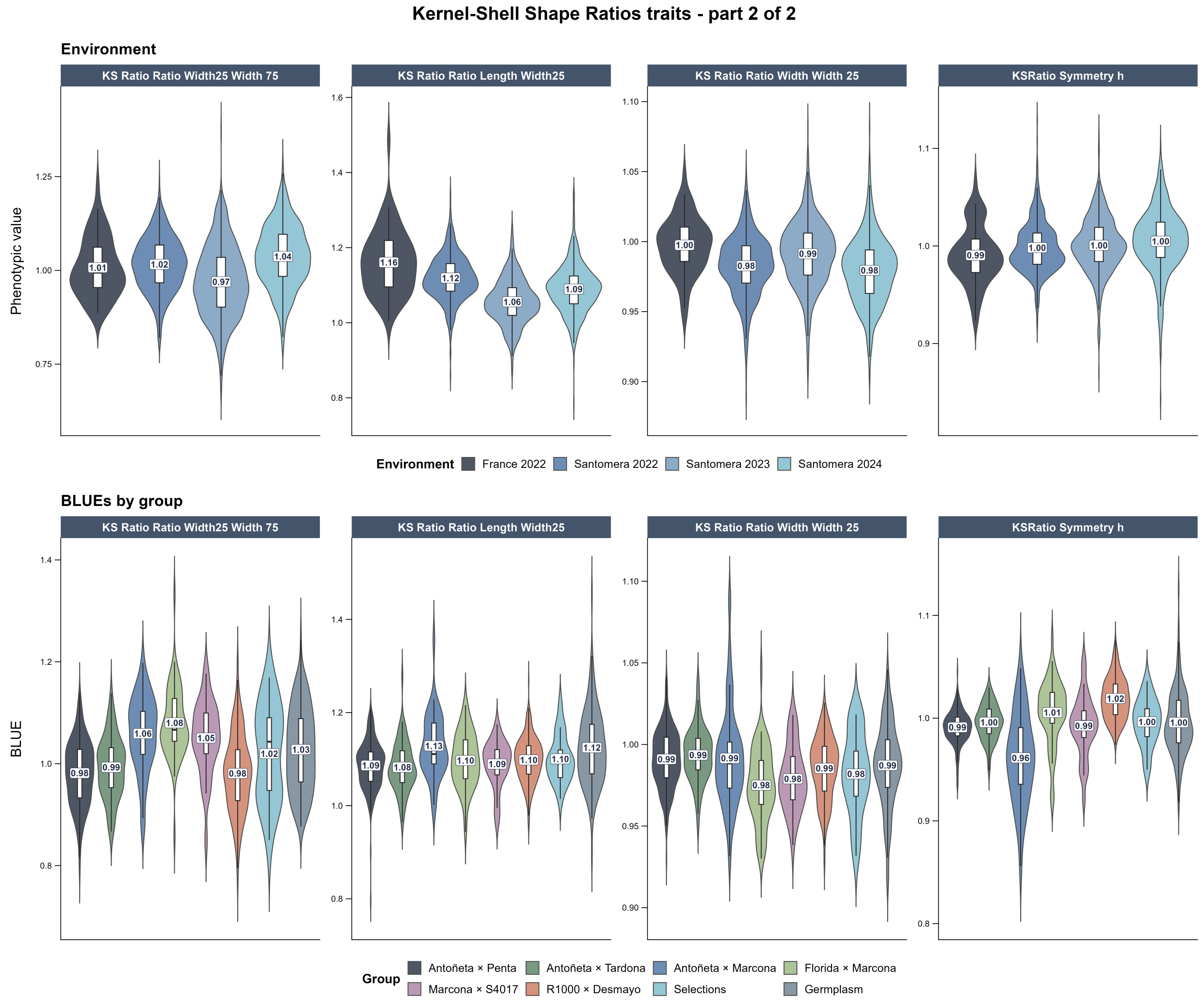


**Figure 27** Distribution of BLUEs by group and environment for kernel-to-shell shape ratios, part 2.

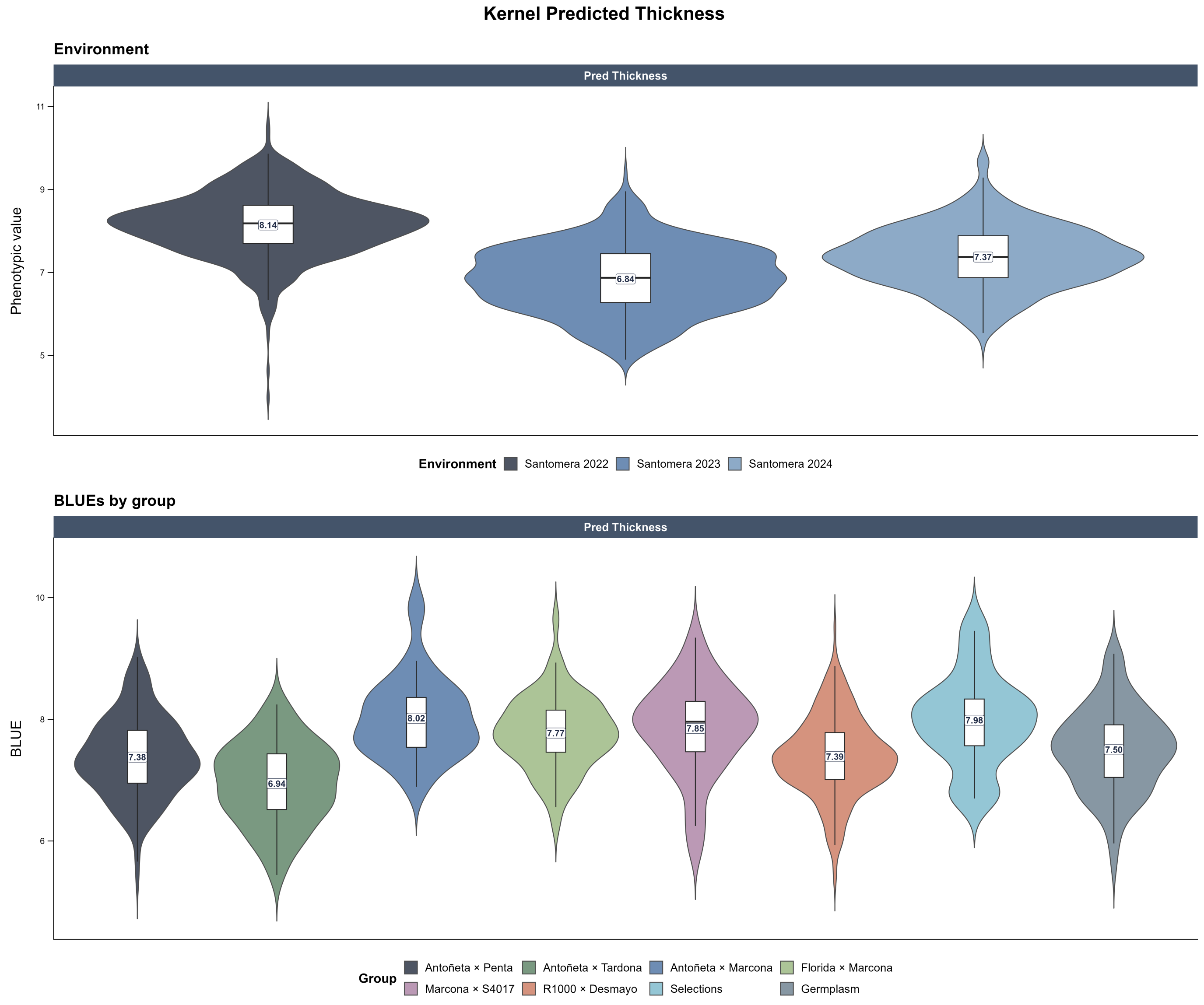


**Figure 28** Distribution of BLUEs by group and environment for predicted shell thickness.

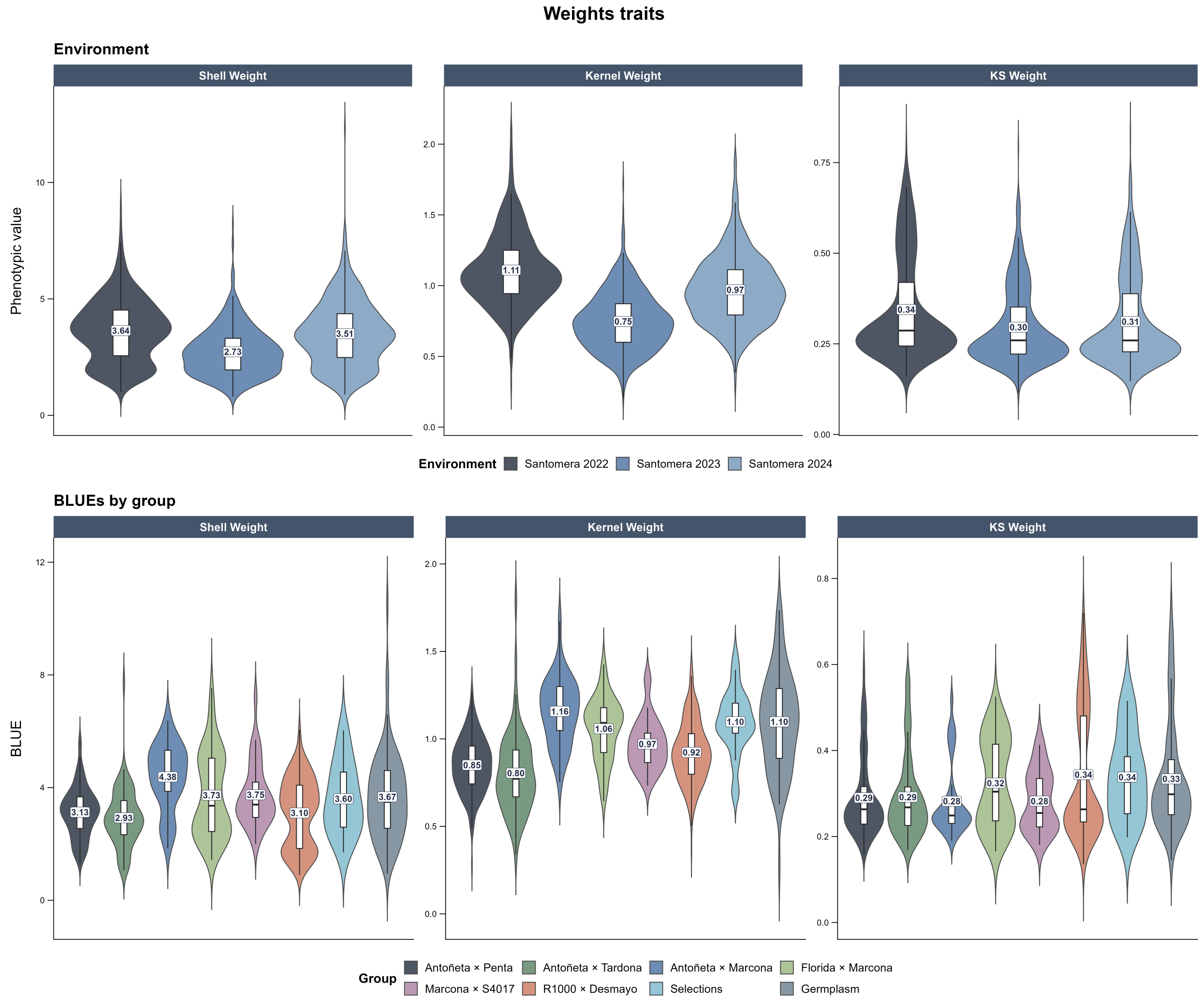


**Figure 29** Distribution of BLUEs by group and environment for kernel and shell weight traits.

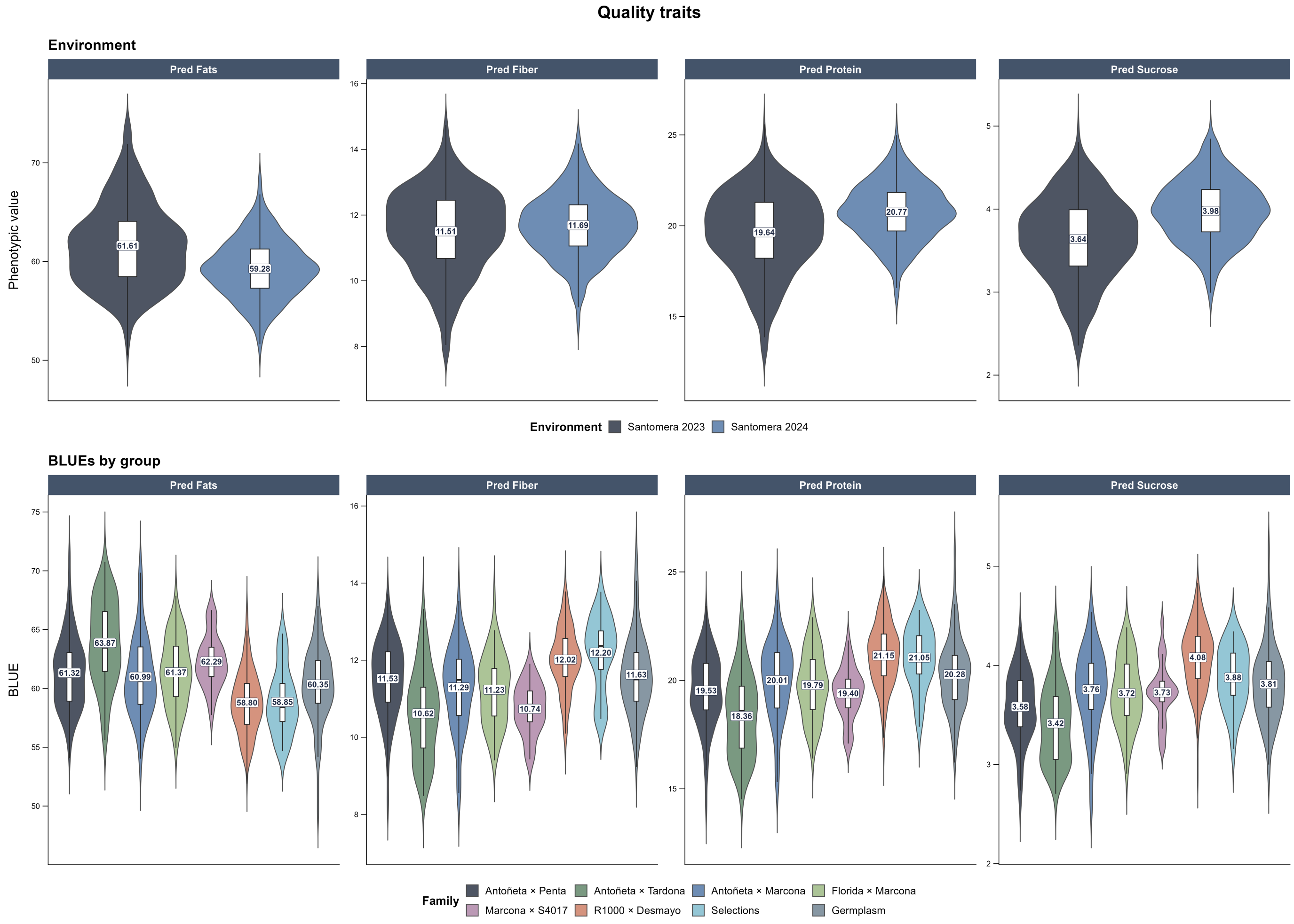


**Figure 30** Distribution of BLUEs by group and environment for almond quality traits.

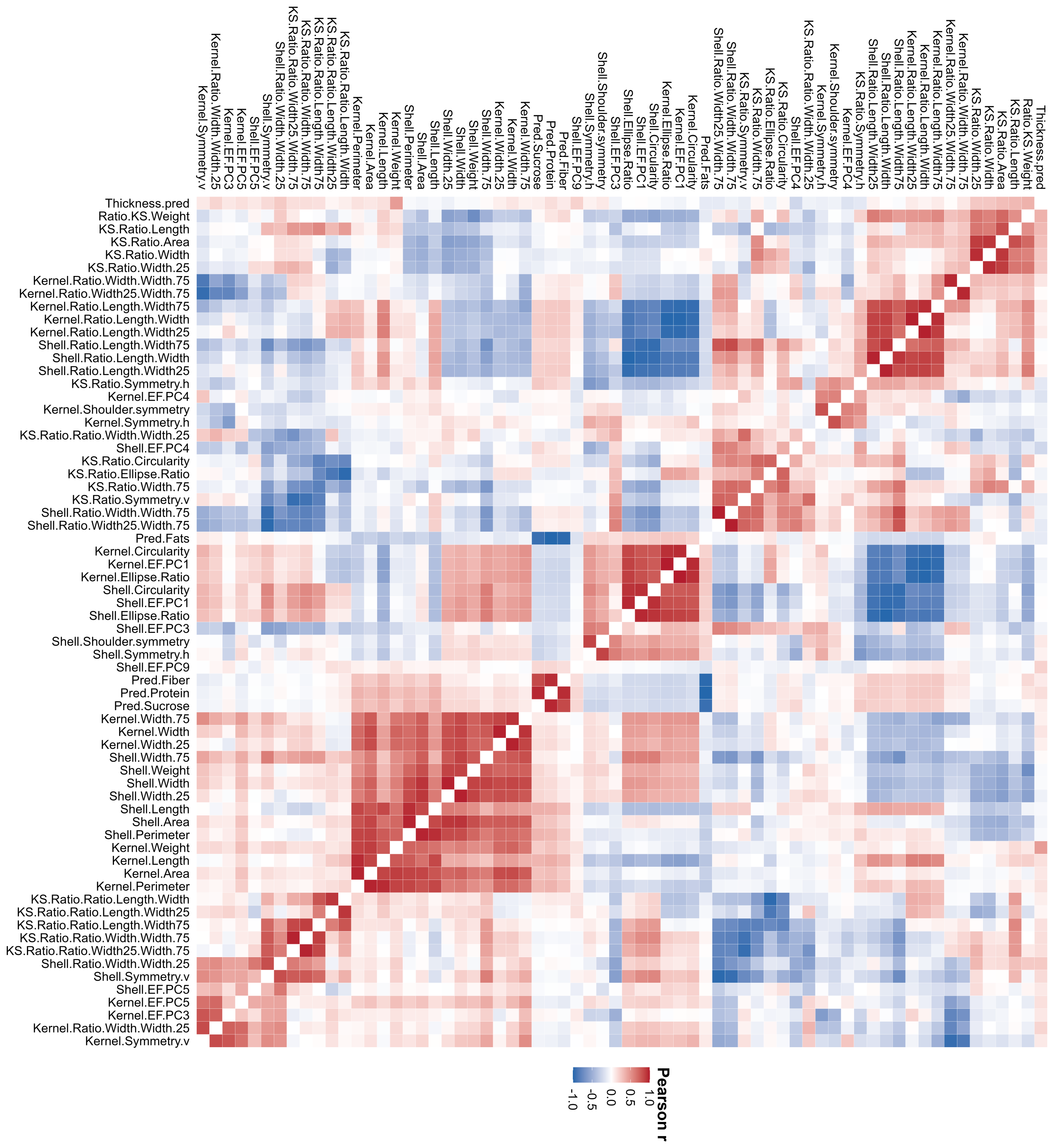

**Figure 31** Correlation heatmap among the evaluated morphological and quality-related traits.

**Appendix 5**

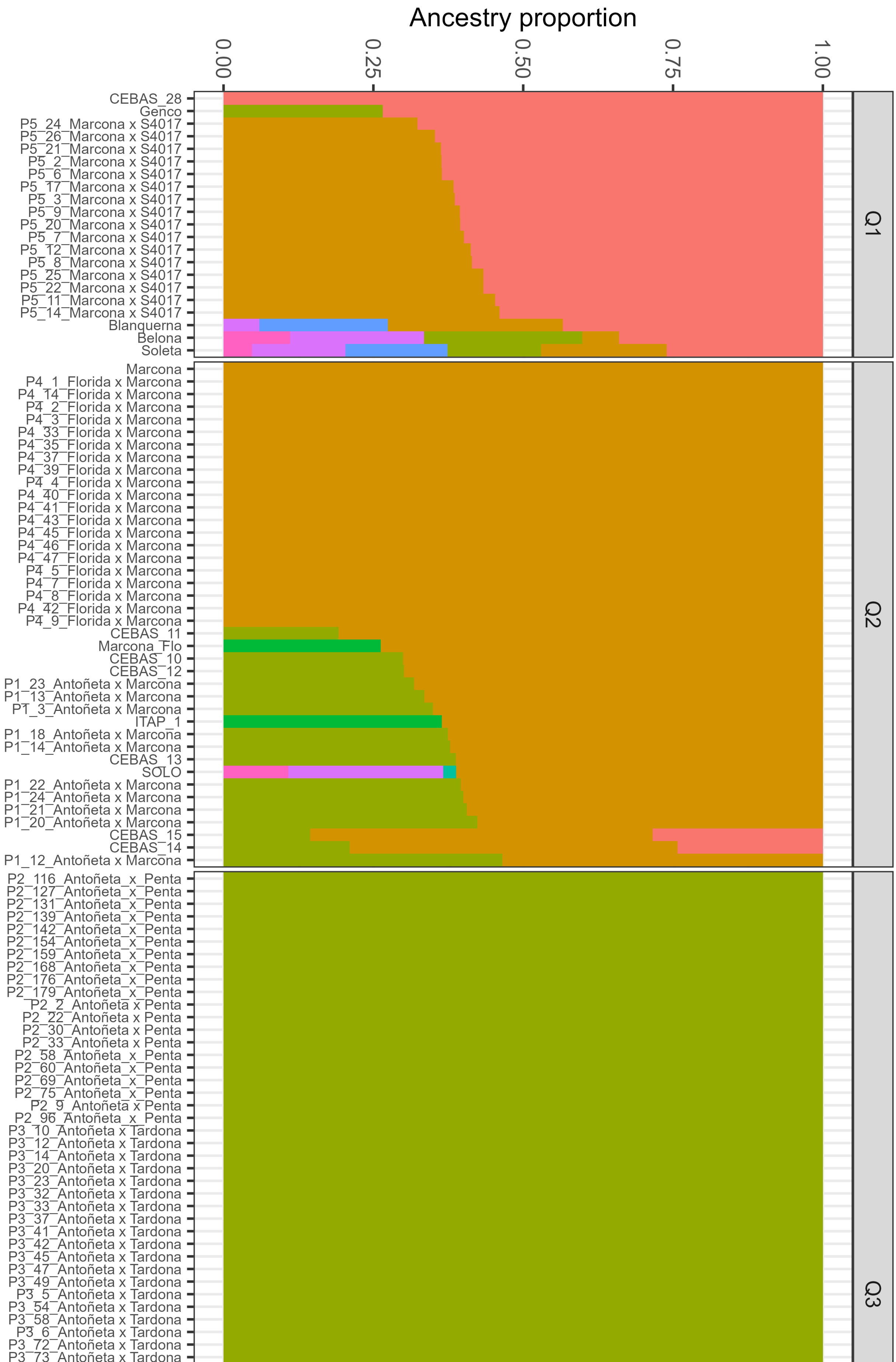
Ancestry proportion
0.00
0.25
0.50
0.75
1.00
Q1
Q2
Q3
CEBAS_28
Genco
P5_24_Marcona x S4017
P5_26_Marcona x S4017
P5_21_Marcona x S4017
P5_2_Marcona x S4017
P5_6_Marcona x S4017
P5_17_Marcona x S4017
P5_3_Marcona x S4017
P5_9_Marcona x S4017
P5_20_Marcona x S4017
P5_7_Marcona x S4017
P5_12_Marcona x S4017
P5_8_Marcona x S4017
P5_25_Marcona x S4017
P5_22_Marcona x S4017
P5_11_Marcona x S4017
P5_14_Marcona x S4017
Blanquerna
Belona
Soleta
Marcona
P4_1_Florida x Marcona
P4_14_Florida x Marcona
P4_2_Florida x Marcona
P4_3_Florida x Marcona
P4_33_Florida x Marcona
P4_35_Florida x Marcona
P4_37_Florida x Marcona
P4_39_Florida x Marcona
P4_4_Florida x Marcona
P4_40_Florida x Marcona
P4_41_Florida x Marcona
P4_43_Florida x Marcona
P4_45_Florida x Marcona
P4_46_Florida x Marcona
P4_47_Florida x Marcona
P4_5_Florida x Marcona
P4_7_Florida x Marcona
P4_8_Florida x Marcona
P4_42_Florida x Marcona
P4_9_Florida x Marcona
CEBAS_11
Marcona_Flo
CEBAS_10
CEBAS_12
P1_23_Antoñeta x Marcona
P1_13_Antoñeta x Marcona
P1_3_Antoñeta x Marcona
ITAP_1
P1_18_Antoñeta x Marcona
P1_14_Antoñeta x Marcona
CEBAS_13
SOLO
P1_22_Antoñeta x Marcona
P1_24_Antoñeta x Marcona
P1_21_Antoñeta x Marcona
P1_20_Antoñeta x Marcona
CEBAS_15
CEBAS_14
P1_12_Antoñeta x Marcona
P2_116_Antoñeta_x_Penta
P2_127_Antoñeta_x_Penta
P2_131_Antoñeta_x_Penta
P2_139_Antoñeta_x_Penta
P2_142_Antoñeta_x_Penta
P2_154_Antoñeta_x_Penta
P2_159_Antoñeta_x_Penta
P2_168_Antoñeta_x_Penta
P2_176_Antoñeta_x_Penta
P2_179_Antoñeta_x_Penta
P2_2_Antoñeta x Penta
P2_22_Antoñeta x Penta
P2_30_Antoñeta x Penta
P2_33_Antoñeta x Penta
P2_58_Antoñeta_x_Penta
P2_60_Antoñeta_x_Penta
P2_69_Antoñeta_x_Penta
P2_75_Antoñeta_x_Penta
P2_9_Antoñeta x Penta
P2_96_Antoñeta_x_Penta
P3_10_Antoñeta x Tardona
P3_12_Antoñeta x Tardona
P3_14_Antoñeta x Tardona
P3_20_Antoñeta x Tardona
P3_23_Antoñeta x Tardona
P3_32_Antoñeta x Tardona
P3_33_Antoñeta x Tardona
P3_37_Antoñeta x Tardona
P3_41_Antoñeta x Tardona
P3_42_Antoñeta x Tardona
P3_45_Antoñeta x Tardona
P3_47_Antoñeta x Tardona
P3_49_Antoñeta x Tardona
P3_5_Antoñeta x Tardona
P3_54_Antoñeta x Tardona
P3_58_Antoñeta x Tardona
P3_6_Antoñeta x Tardona
P3_72_Antoñeta x Tardona
P3_73_Antoñeta x Tardona
